# HIF2α is a Direct Regulator of Neutrophil Motility

**DOI:** 10.1101/2020.06.05.137133

**Authors:** Sundary Sormendi, Mathieu Deygas, Anupam Sinha, Anja Krüger, Ioannis Kourtzelis, Gregoire Le Lay, Mathilde Bernard, Pablo J. Sáez, Michael Gerlach, Kristin Franke, Ana Meneses, Martin Kräter, Alessandra Palladini, Jochen Guck, Ünal Coskun, Triantafyllos Chavakis, Pablo Vargas, Ben Wielockx

## Abstract

Orchestrated recruitment of neutrophils to inflamed tissue is essential during initiation of inflammation. Inflamed areas are usually hypoxic, and adaptation to reduced oxygen pressure is typically mediated by hypoxia pathway proteins. However, it is still unclear how these factors influence the migration of neutrophils to and at the site of inflammation either during their transmigration through the blood-endothelial cell barrier, or their motility in the interstitial space. Here, we reveal that activation of the Hypoxia Inducible Factor-2 (HIF2α) due to deficiency of HIF-prolyl hydroxylase domain protein-2 (PHD2) boosts neutrophil migration specifically through highly confined microenvironments. *In vivo,* the increased migratory capacity of PHD2-deficient neutrophils resulted in massive tissue accumulation in models of acute local inflammation. Using systematic RNAseq analyses and mechanistic approaches, we identified RhoA, a cytoskeleton organizer, as the central downstream factor that mediates HIF2α-dependent neutrophil motility. Thus, we propose that the here identified novel PHD2-HIF2α-RhoA axis is vital to the initial stages of inflammation as it promotes neutrophil movement through highly confined tissue landscapes.

## Introduction

In the innate immune response, neutrophils represent the first line of protection against infections, extravasating quickly from circulation to inflamed tissues for fast pathogen elimination. This process necessitates transit from an oxygen-rich circulatory system to the inflammation site, which is typically hypoxic due to vasculature damage and/or high metabolic demand of pathogens and host cells.^1^ Thus, neutrophil adaptation to low oxygen levels is crucial during the early phases of the inflammatory response.

Under hypoxic conditions, the transcription factors Hypoxia Inducible Factor-1 (HIF1α) and its isoform HIF2α are key elements that control immune cell metabolism and function,^2–7^ and importantly, HIF activity is controlled by a class of oxygen sensors known as the HIF prolylhydroxylase domain enzymes (PHD1-3) (reviewed in^8,9^). When oxygen levels decrease, PHDs get inactivated, which results in HIFα stabilization and transcription of relevant target genes. Interestingly, HIF1α-deficiency results in subdued inflammation^2,3^ while, inversely, PHD inactivation and/or HIFα stabilization leads to enhanced neutrophil survival,^4,10^ chemotaxis and degranulation (reviewed in^11^). Although both HIFα subunits have overlapping activities, unique roles for HIF2α, including in neutrophil function, have been reported.^4–6^

Over the past decade, several mechanisms have been shown to participate in the multi-step recruitment of neutrophils from circulation to sites of infection or inflammation.^12–14^ The recruitment process requires cell plasticity because cells deform as they move through the blood-endothelial cell barrier and the confined areas of interstitial tissues. Leucocyte migration through these microenvironments is orchestrated by actin polymerization regulators, such as the rho GTPases RhoA, Cdc42 and Rac1.^15–18^ In this context, HIF1α expression has been suggested to modulate both functional changes in the cytoskeleton and metabolic reprograming^19–22^. Importantly, disruption of mechanisms that control neutrophil infiltration in tissues is associated with sepsis, a life-threatening condition with multi-organ failure and one of the leading causes of death in the intensive care unit (ICU).^23^ Conversely, till date, no effective therapeutic strategies are available for mitigating an uncontrolled neutrophilic inflammatory response.

In this study, we address the effects of PHD2-deficiency on the motility of neutrophils, including their recruitment during localized inflammation. Using *ex vivo* and *in vivo* imaging in a variety of highly confined microenvironments, we demonstrate for the first time that HIF2α over-activation enhances the migratory capacity of neutrophils in a chemotaxis-independent manner. Through whole transcriptome analysis and combined migratory regulation, we describe a role for the PHD2-HIF2α-RhoA axis in the prompt initiation of the innate immune response.

## Materials and methods

### Mice

All mouse strains were housed in our local mouse facility under specific pathogen-free conditions. Experiments were performed with male and female mice at the age of 8 to 12 weeks. Vav:cre-PHD2^f/f^ (cKO P2) and Vav:cre-PHD2/HIF2^ff/ff^ (cKO P2H2) mouse lines were created in our laboratory, using PHD2^f/f^,^24^ Vav:cre^25^ (generous gift from Dr. Graf, Spain) and/or HIF2α^f/f^.^26^ All offspring were born in normal Mendelian ratios and individual floxed lines have been previously backcrossed to C57BL/6J for at least 9 times. WT controls in all experiments were Cre-negative littermates without any chimerism (partial deletion of floxed genes in early blastomeres).^25^ Mice were genotyped using primers described in supplemental Table 1 and knock-down efficiency confirmed via qRT-PCR on isolated neutrophils (supplemental Figure 1A, C) and/or genomic PCR on ear biopsies.^27^ KRN TCR transgenic mice were inter-crossed with NOD Shilt/J mice (Charles River, Italy) to generate K/BxN mice as described previously.^28^ A detailed description of the inflammation models can be found as supplemental data. Breeding of all mouse lines and animal experiments were in accordance with the local guidelines on animal welfare and were approved by the Landesdirektion Sachsen, Germany.

### Histological analysis

5μm thick cryo-sections from 24-hour PMA-treated ears or 7μm thick cryo-sections of knees from 5-day K/BxN-treated mice were were incubated for 1 hour at 37 °C with primary antibodies to detect Gr1+ neutrophils or cCas3^+^ apoptotic cells. Imaging was performed using an epi-fluorescence microscope with Zeiss EC Plan-Neofluar objectives. Number of Gr1^+^ cells per tissue area and percentage of cCas3^+^ in Gr1^+^ cells were quantified using *Zen* software Version 3.1 (see supplemental Table 2 for more information on antibodies).

### Flow cytometry

Immune cell profile of synovial fluid from 5-day K/BxN-treated arthritic knee joints was assessed via FACS performed on LSRII (Becton Dickinson), and cell numbers were counted on MACS quant (Miltenyi). After knee isolation, digestion to extract the cellular compartment of the synovial cavity was performed using collagenase D, Dispase II, and DNAse I in DMEM. Knees were incubated for 30 minutes at 37 °C the supernatant was centrifuged, washed and single cells stained for specific myeloid cell markers using the following fluorophore-conjugated antibodies for 30 minutes at +4 °C (see supplemental Table 2 for more information on antibodies).

### Bone Marrow-Derived neutrophils (BMDN)

BMDNs were obtained by crushing long bones the bones in 5% FCS using a mortar and either isolated by negative selection using the EasySep Mouse Neutrophil Enrichment Kit (Stemcell Technologies) or by positive selection using biotinylated antibodies (see supplemental data for more details).

### 1-D and 2-D confined migration in micro-channels

Customized polydimethylsiloxane (PDMS) micro devices containing micro-channel areas and 2D-free areas were used to study cell migration in highly confined environments as described previously.^29^ PDMS micro-chips were coated with fibronectin (10μg/ml), their nuclei prelabelled with Hoechst for 30 minutes at 37°C and 10^5^ neutrophils (in 5μl) loaded in 3μm-diameter wells. Migrating neutrophils were imaged by video-microscopy (Leica DMI8). Images were analysed using *Fiji* software^30^ and a customised script was used to create kymographs for individual migrating cells within the micro-channels. Cell speed was calculated using *MATLAB* software (The MathWorks, Inc). 2D-confined random migration was analysed using *Imaris* (Bitplane) cell tracking software. Where indicated and prior to their loading in the PDMS micro-device, cells were resuspended in media containing 1 μM of CCG-100602 (from Sigma), 1 μM of ML141 (from Sigma) or 1 μM of the Cell permeant C3 transferase (from Cytoskeleton).

### Cell migration in 3D collagen matrices

Neutrophil migration in a 3D environment was evaluated in customized PDMS chambers filled with varying concentrations of a collagen matrix (3-5 mg/ml) containing 2×10^6^ neutrophils/ml. After polymerization of the collagen matrix, phase imaging was performed using videomicroscopy (DMI8). For the chemotaxis assay, CXCL2 (20ng/ml) was added locally to the matrix.

### Rho-GTPase activity assays

The activity of Activated RhoA, Rac and Cdc42 was measured in lysates from negatively sorted neutrophils from all genotypes by performing respective activation G-LISA assays (RhoA, Rac1,2,3 and Cdc42 G-LISA Activation Assays; Cytoskeleton) on fibronectin-coated plates as per manufacturer’s protocol.

### Statistics

Data and graphs represent mean ± SEM of representative experiments. Statistical significance was calculated using the Mann Whitney U test (unpaired) or the Wilcoxson matched-pairs signed rank test (paired) using GraphPad Prism (v7.02 or higher); *p<0.05 was considered statistically significant.

### Data Sharing Statement

RNAseq data are available at GEO (**GSE151703**).

Additional data may be found in a data supplement available with the online version of this article.

For original data, please contact

## Results

### PHD2-deficient neutrophils display enhanced migration in highly confined environments in a HIF-2α-dependent manner

Although changes in the hypoxia pathway are involved in multiple stages of the inflammatory response, details on how the PHD/HIF axis governs neutrophil migration remain elusive. Given that PHD2 is a central regulator of the hypoxia response, we studied the motility of PHD2-deficient BMDNs isolated from vav:cre-PHD2^f/f^ mice (henceforth denoted as cKO P2; supplementary Figure 1A). Initially, *1D migration assays* in polydimethylsiloxane (PDMS) micro-channel devices of different levels of constriction (channel widths of 3, 4 or 5μm) were used to characterize the migratory capacity of individual neutrophils (Figure 1A).^17,18,31–34^ Interestingly, cKO P2 neutrophils moved significantly faster than their WT counterparts, but only in the most confined channels (Figure 1B and Supplemental Figure 1B). To identify downstream effectors of this phenotype, we evaluated the contributions of HIF2α, a PHD2 target and a central factor in inflammation^4,35^, in cKO P2H2 neutrophils compared to their littermate controls (Supplemental Figure 1C). Interestingly, there were no differences in speed at any of the degrees of confinement tested (Figure 1C). These data strongly suggest that enhanced HIF2α activation regulates neutrophil motion in very confined microenvironments.

**Figure 1.**
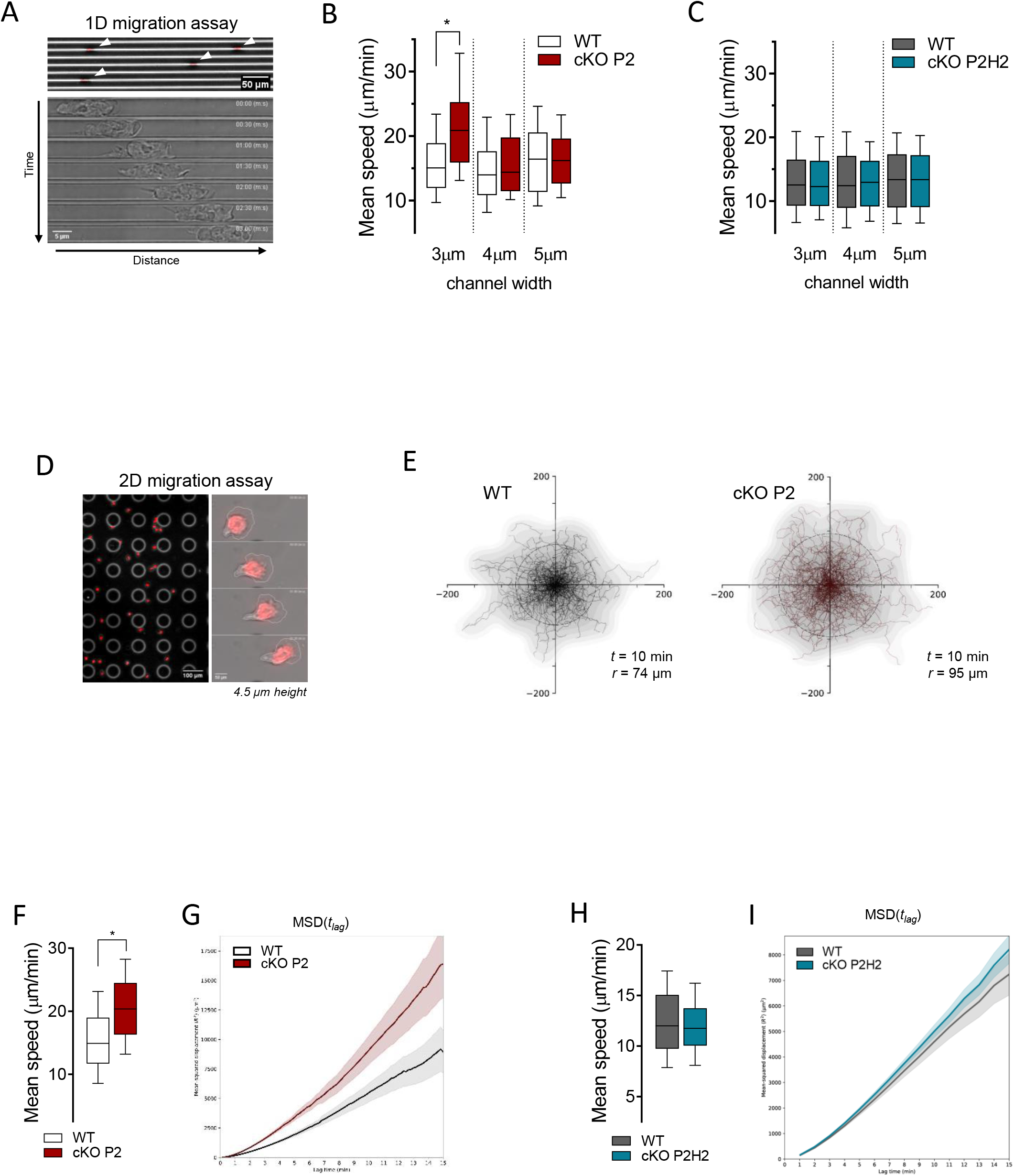

We extended our analysis to evaluate neutrophil migration in a *2D confined microenvironment* (4.5 μm height) (Figure 1D). Similar to the results obtained in the 1D migration assay, neutrophils from cKO P2 mice showed increased motility compared to their WT counterparts, as evidenced by longer trajectories of equivalent durations (Figure 1E), as well as greater speed (Figure 1F), along with higher mean square displacement (MSD) values (Figure 1G). On the other hand, under identical conditions, cKO P2H2 neutrophils did not show any difference in speed or MSD compared to their WT counterparts (Figure 1H and 1I). Interestingly, cell migration in a non-confining 2D chamber (12μm height) showed no difference in speed, trajectories, or MSD (Supplemental Figure 1D-F). Thus, these data indicate that the PHD2-HIF2α pathway regulates cell migration by facilitating mobility strictly in confined spaces.

### PHD2-deficient neutrophils display enhanced non-directed motility in complex environments

We used *3D-collagen matrices* to confirm the role of PHD2 in neutrophil migration in a microenvironment of fibers and different pore sizes, adequately mimicking the tissue complexity *in vivo.* Therefore, migration of freshly-isolated BMDNs from cKO P2 mice and WT littermates was compared in dense 3D collagen gels (4mg/ml) (Figure 2A) during which cKO P2 neutrophils showed greater motility, as evidenced by a higher displacement radius (Figure 2A). Detailed analysis of their random trajectories showed that cKO P2 neutrophils displayed greater speed and MSD values compared to WT cells (Figure 2B, C). Interestingly, this difference was completely lost in less dense collagen gels (2mg/ml) (Figure 2B and Supplemental Figure 2A).

**Figure 2.**
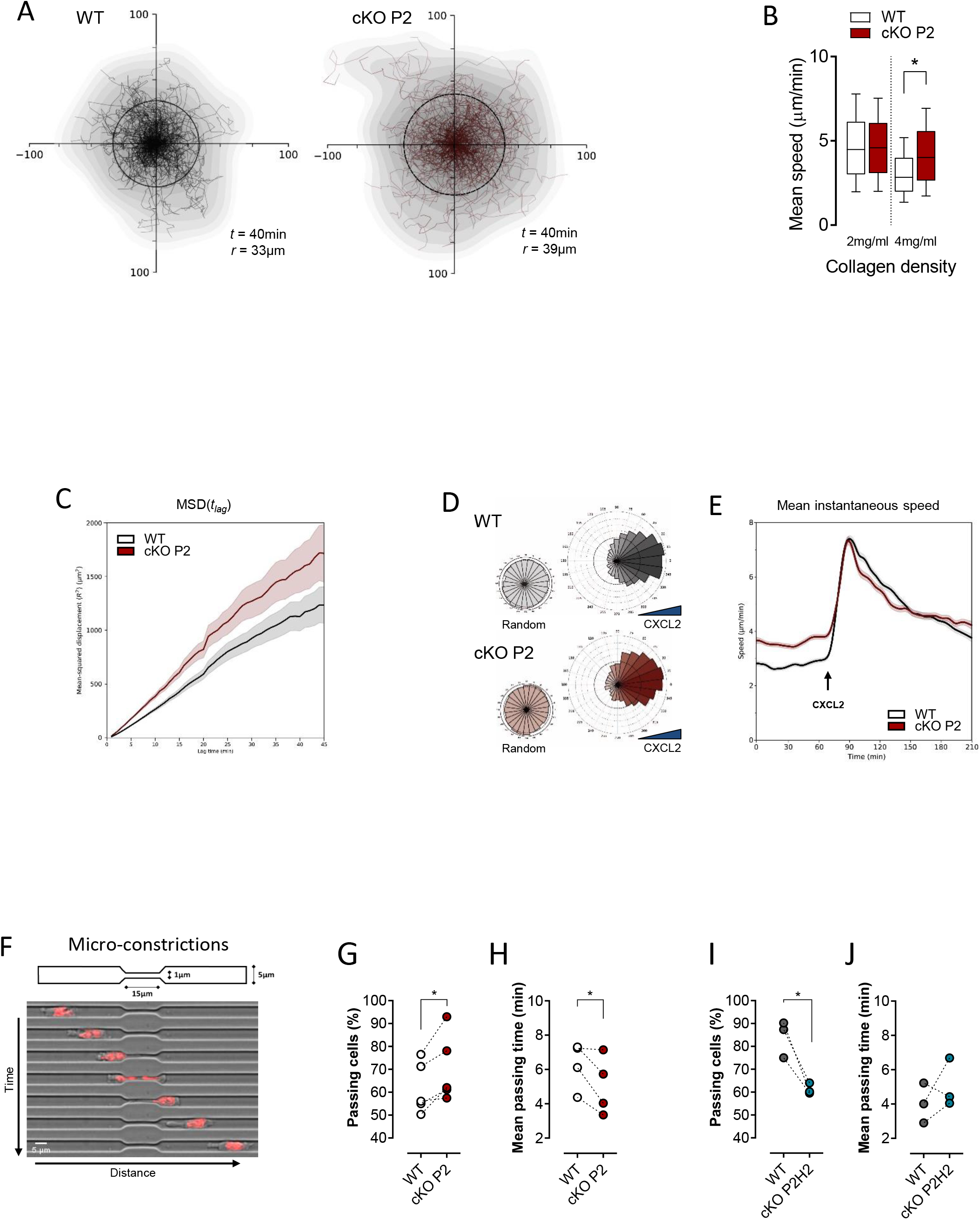

As it has been suggested that silencing of PHD2 in neutrophils leads to their enhanced chemotaxis,^36^ we assessed this effect in our complex 3D collagen matrix setup using CXCL2 as a classical neutrophil chemokine. Neutrophil trajectory analysis in dense collagen gels (4mg/ml) showed that absence of PHD2 did not affect neutrophil chemokine sensing because their directionality towards CXCL2 remained unaltered (Figure 2D) and a similar strong increase in cell speed was found in both PHD2-deficient and WT neutrophils (Figure 2E). Thus, these results show that PHD2-deficient neutrophils display an enhanced migratory capacity in dense 3D collagen gels and that PHD2 loss does not affect CXCL2-induced chemotactic capacity. In other words, the faster migration of cKO P2 neutrophils is independent of chemotaxis induction and is rather linked to enhanced undirected motility or chemokinesis.

The migratory capacity of several cell types in complex microenvironments is highly dependent on their capacity to deform when encountering narrow pores.^37,38^ Therefore, we evaluated whether cKO P2 neutrophils can overcome severely constricted spaces of only 1μm width (Figure 2F).^17,39^ Remarkably, PHD2-deficient neutrophils showed an enhanced preference to pass through these constrictions (Figure 2G) and were also faster compared to WT neutrophils (Figure 2H). Interestingly, under these conditions, cKO P2H2 neutrophils showed reduced migration; and, similar migration kinetics than WT cells (Figure 2I, J), again suggesting a PHD2/HIF2α-dependent axis in migration through extreme narrow constrictions.

Next, we studied whether the ability of cKO P2 neutrophils to pass through small confinements is related to changes in their deformability when an external force is applied. For this, we first analyzed neutrophil deformability using *real time fluorescence and deformability cytometry* (RT-FDC) (see supplemental data), which can extract the stiffness of cells (Young’s Modulus) in high-throughput, without contact at ms-timescales.^40,41^ We used steady-state BMDNs, Phorbol 12-Myristate 13-Acetate (PMA)-activated BMDNs, and peripheral blood neutrophils isolated at 6h after thioglycolate-induced peritonitis. However, no differences were observed between cKO P2 and WT neutrophils under any of the conditions tested (supplemental Figure 2B). Likewise, a *microcapillary microcirculation mimetic* (MMM) assay^42,43^ using peritonitis neutrophils showed no difference in their ability to passively navigate through multiple constrictions at high speed (supplemental Figure 2C). Taken together, these assays strongly suggest that loss of PHD2 does not affect neutrophil deformability under externally applied stress without confinement.

### PHD2-deficient neutrophils extravasate faster *in vivo* and accumulate in inflamed tissue

Based on the enhanced ability of PHD2-deficient neutrophils to overcome very small constrictions, we decided to study the behavior of these cells *in vivo*; specifically, in a more complex setting of sterile skin inflammation. Ear lobes from cKO P2 and WT littermate mice that displayed no difference in total numbers of hematopoietic stem cells, myeloid progenitors or mature neutrophils (supplemental Figure 3A), were ectopically treated with PMA and the recruitment of Ly6G^+^ cells was visualized using intra-vital 2-photon microscopy (Figure 3A). In line with our previous experiments, we found that PHD2-deficient neutrophils were able to extravasate about 30% faster from the vessel into the ear tissue than their WT counterparts (Figure 3B-C). Furthermore, the cumulative effect of faster neutrophil extravasation time resulted in an anticipated increase in Gr1^+^ cells in the inflamed cKO P2 ear compared to that in WT littermates at 24 hours after PMA-treatment (Figure 3D). Conversely but consistently, this difference in migration was abolished in cKO P2H2 mice (Figure 3E), further confirming a role for HIF2α activity in driving increased migration capacity of these neutrophils.

**Figure 3.**
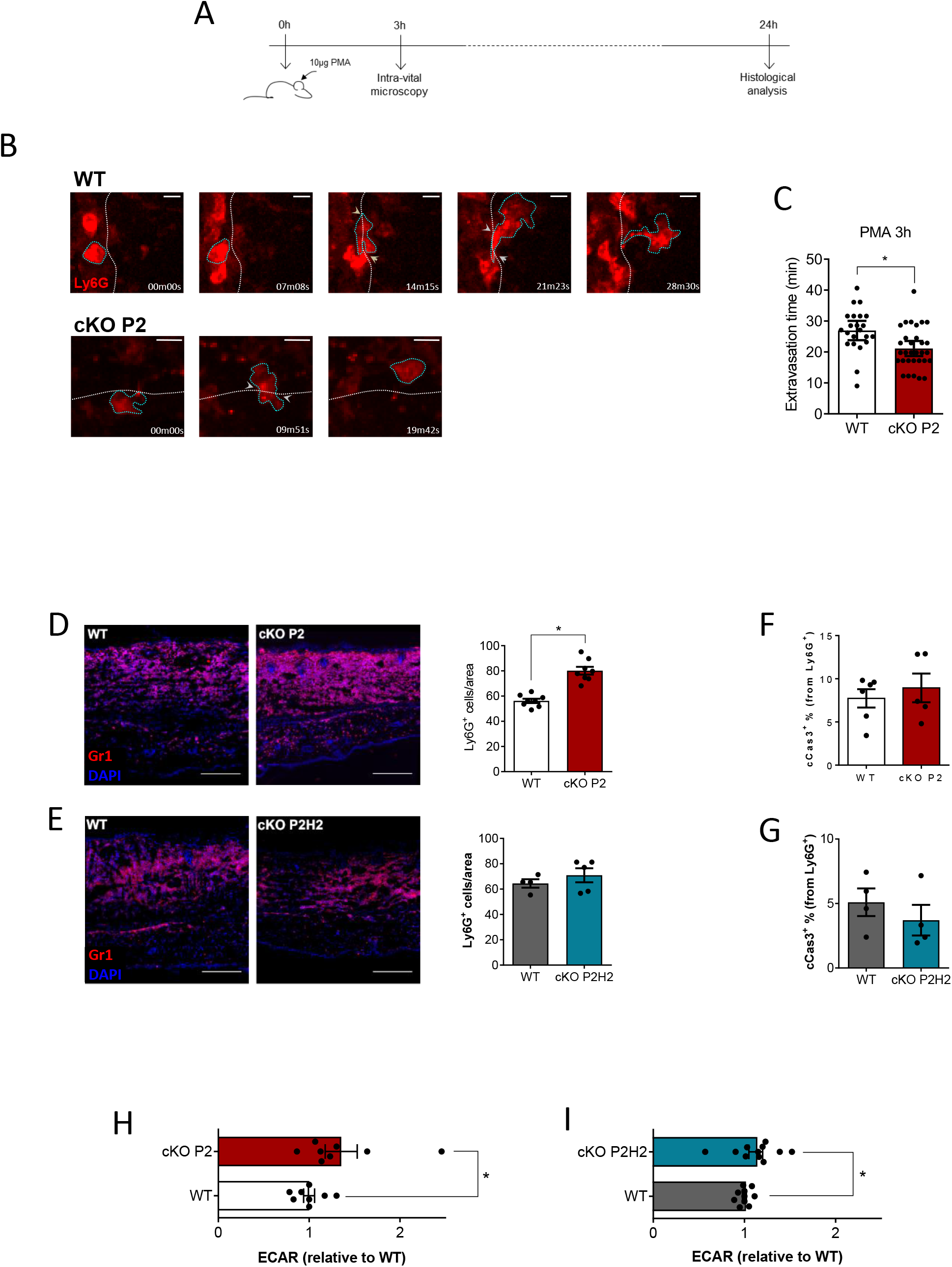

As previous studies have described PHD2-related improved survival of neutrophils during inflammation,^4,44^ we evaluated the level of apoptotic cells in 24 hour PMA-treated ears, but found no difference in cleaved caspase-3^+^ cell numbers (cCas3^+^) between the different genotypes (Figure 3F, G; supplemental Figure 3B, C). Additionally, as recent work has associated PHD2 with enhanced neutrophil glycolysis and their recruitment to sites of inflammation,^36^ we assessed the glycolytic capacity of BMDNs from cKO P2, cKO P2H2, and their respective WT counterparts by measuring extracellular acidification rate (ECAR). In line with previous reports, PHD2-deficient neutrophils appeared to be significantly more glycolytic than their respective WT counterparts (Figure 3H). However, neutrophils lacking both PHD2 and HIF2α also showed significantly higher glycolysis (Figure 3I). Taken together, although HIF2α directly controls the migration speed of neutrophils in confined spaces and inflamed tissues, this effect is independent of their survival or glycolytic activity.

### HIF2α stabilization upon loss of PHD2 affects cytoskeletal gene expression profiles

It is well-accepted that the functionality of innate immune cells varies depending on the lipid-type composition of its cytoplasmic membrane.^45,46^ Therefore, we evaluated if altered membrane lipid composition of the cKO P2 neutrophils could account for their different migratory ability, by performing high-throughput lipidomic analysis of freshly isolated BMDNs (see supplemental data). However, as there were no significant alterations between the cKO P2 and WT BMDNs (supplemental Figure 2D and Table 3), it appears unlikely that differences in the lipid composition are directly responsible for the dramatic difference in the migratory capacity of the cKOP2 neutrophils.

Next, to further characterize the molecular underpinnings of the HIF2α-driven neutrophil migration phenotype, we used *next generation sequencing* (NGS) wherein the steady state transcriptome of BMDNs derived from cKO P2 and cKO P2H2 mice were analyzed and compared to that from their respective WT counterparts (Figure 4A). Gene signatures of various lineages were evaluated using gene set enrichment analyses (GSEA) as described previously.^47–49^ In line with our *in vivo* cCas3+ results, we detected no significant apoptosis signatures among any of the genotypes (Figure 4B) and NGS confirmed a significant enrichment of glycolysis/gluconeogenesis related genes in both cKO P2 and cKO P2H2 BMDNs (Figure 4C). Strikingly, steady state BMDNs lacking PHD2, with or without HIF2α, displayed a significant reduction in genes related to the innate immune response but not the chemokine signaling pathway (Figure 4D). Together, these observations suggest that significant HIF2α-independent changes in glycolytic capacity and immune response of PHD2-deficient neutrophils can be likely linked to HIF1α activity, as previously suggested.^2,36^ Conversely, a number of HIF2α-dependent gene signatures associated with PHD2 deficiency related to function and structure of the neutrophil cytoskeleton, including Rho GTPase activity (Figure 4E). Additionally, using an integrative method, we identified a number of HIF2α-associated *master regulators* that could potentially control cellular cytoskeletal rearrangements through transcriptional or protein regulation (supplemental Figure 4A).

**Figure 4.**
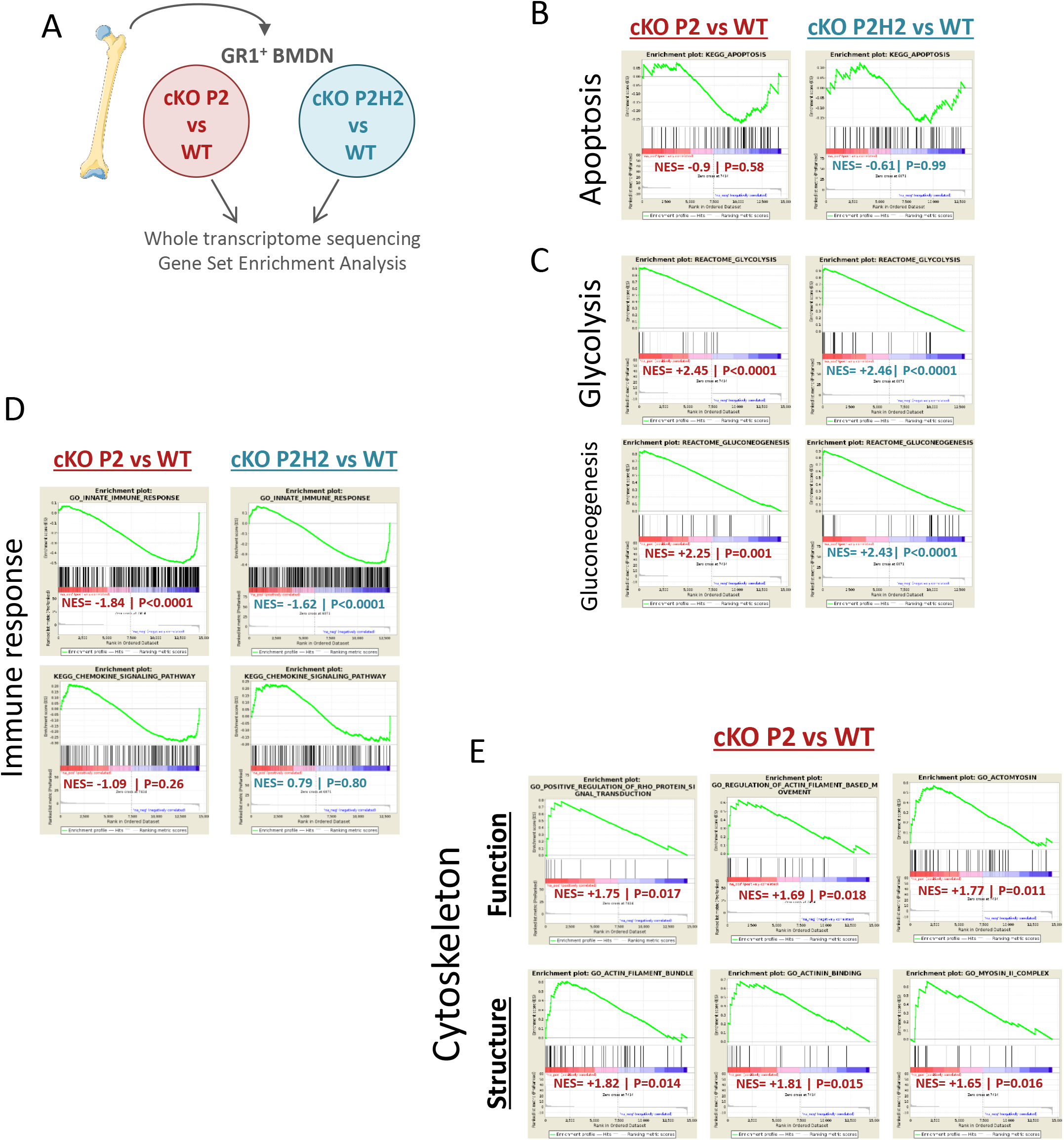

### Diminished RhoA GTPase activity underlies the faster HIF2α-dependent migration of PHD2-deficient neutrophils

Small Rho GTPases (RhoA, cdc42 and Rac) are the final molecular effectors that steer cytoskeletal dynamics. In line with this, we identified numerous potential associations (direct and/or indirect) among 49 genes/proteins and with RhoA and/or Cdc42 (bold lines), but not Rac GTPase (Figure 5A). Notably, 7 of these genes have been previously identified as being associated with HIF2α binding sites (supplemental Figure 4B).^50^

To substantiate this link between PHD2/HIF2α and Rho GTPases, we used an *ex vivo* enzymatic assay to quantify the activity of these Rho GTPases in untreated freshly-isolated BMDNs from cKO P2 and P2H2 mice. Interestingly, cKO P2 neutrophils exhibited diminished RhoA and Cdc42 GTPase activity (Figure 5B, C), while Rac GTPase activity was comparable with WT neutrophils (Figure 5D). Further, cKO P2H2 neutrophils displayed no significant reduction in either RhoA, Cdc42 or Rac GTPase activity, suggesting that regulation of RhoA and/or Cdc42 is dependent on the PHD2/HIF2α-axis (Figure 5B-D).

Given this reduction in RhoA and Cdc42 GTPase activity in PHD2-deficient neutrophils, we examined whether their direct inhibition in WT neutrophils can mimic the motility phenotype displayed by cKO P2 neutrophils. Therefore, we performed a series of *ex vivo* 1D-migration assays using the RhoA inhibitor CCG100602 (CCG) or the Rho inhibitor Exoenzyme C3 Transferase (C3) and found that while the use of low doses of CCG or C3 enhanced the speed of migrating neutrophils in 3μm micro-channels (Figure 5E), treatment of cells with a Cdc42 inhibitor (ML141) did not have any effect on the velocity of BMDNs (Figure 5F). Taken together, our data strongly argue for a PHD2/HIF2α–orchestrated regulatory loop in RhoA GTPase activity-dependent motility of BMDNs.

**Figure 5.**
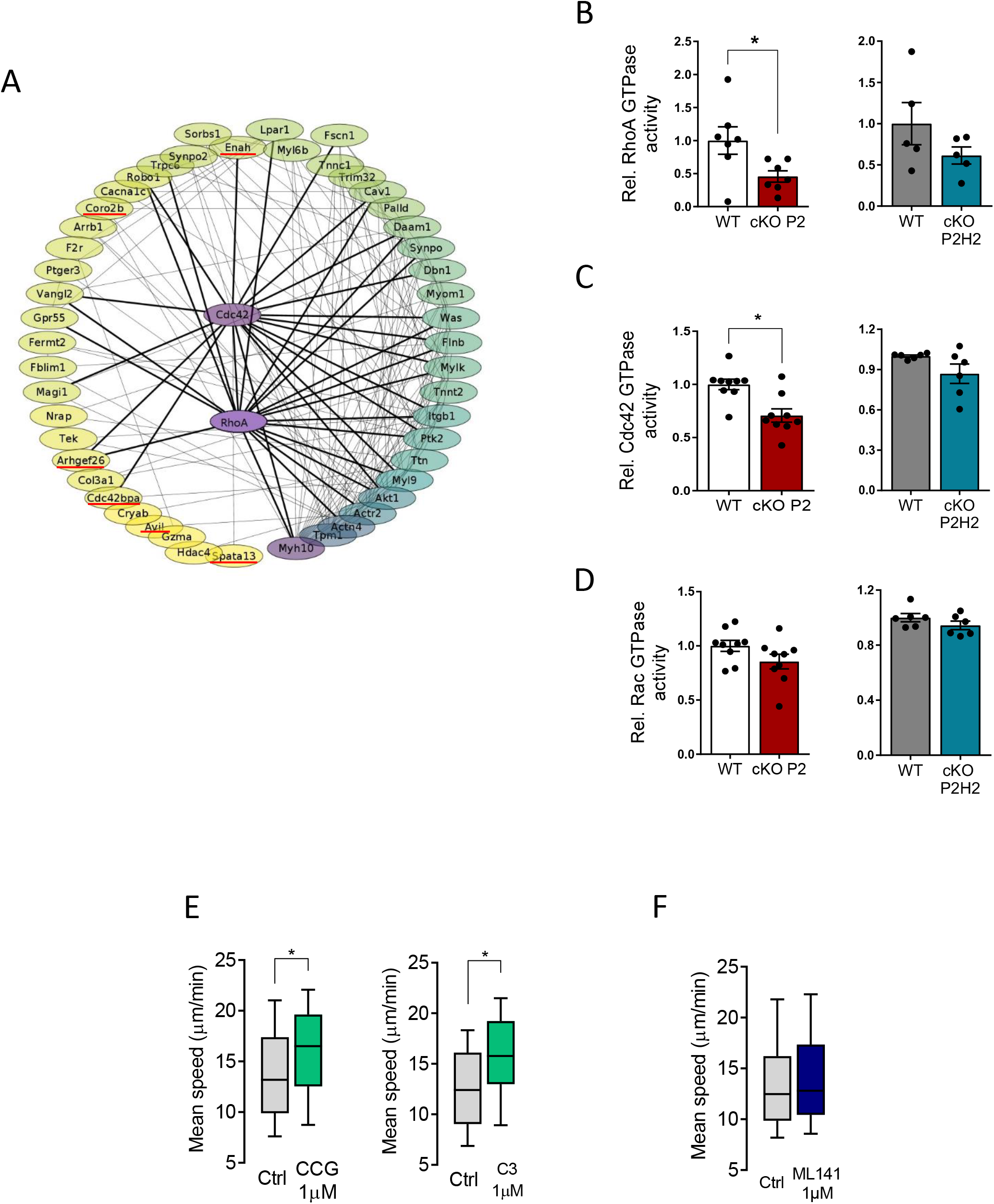

### The PHD2/HIF2α-axis controls neutrophil accumulation in joints during severe inflammatory arthritis

To test the biological effects of the enhanced migratory capacity of PHD2-deficient neutrophils, we subjected the different mouse strains to an autoantibody-induced inflammatory arthritis model (K/BxN), which has been shown to be myeloid dependent (Figure 6A).^51,52^ cKO P2 mice displayed enhanced swelling of the hind limbs compared to their WT littermates (Figure 6B) and this effect was sustained throughout the first 2 weeks of the experiment. In line with our previous results, cKO P2H2 and their WT littermates displayed no difference in swelling (Figure 6C). To characterize the myeloid composition of the inflamed knee joints, we performed flow cytometry analysis of the synovial fluid drawn on day 5, which revealed much higher accumulation of neutrophils in cKO P2 knee joints (>3-fold increase versus WT), along with slightly enhanced macrophages (Figure 6D); immunofluorescence for Gr1 on knee joints further confirmed this observation (Figure 6E). Conversely, although no differences were observed in joint swelling between cKO P2H2 mice and their WT littermates, their synovial fluid showed a slight but significant reduction in neutrophil numbers at day 5 compared to cKO P2 mice (Figure 6F). Thus, also in arthritic joints PHD2/HIF2α is a central axis during the initial stages of the inflammation.

**Figure 6.**
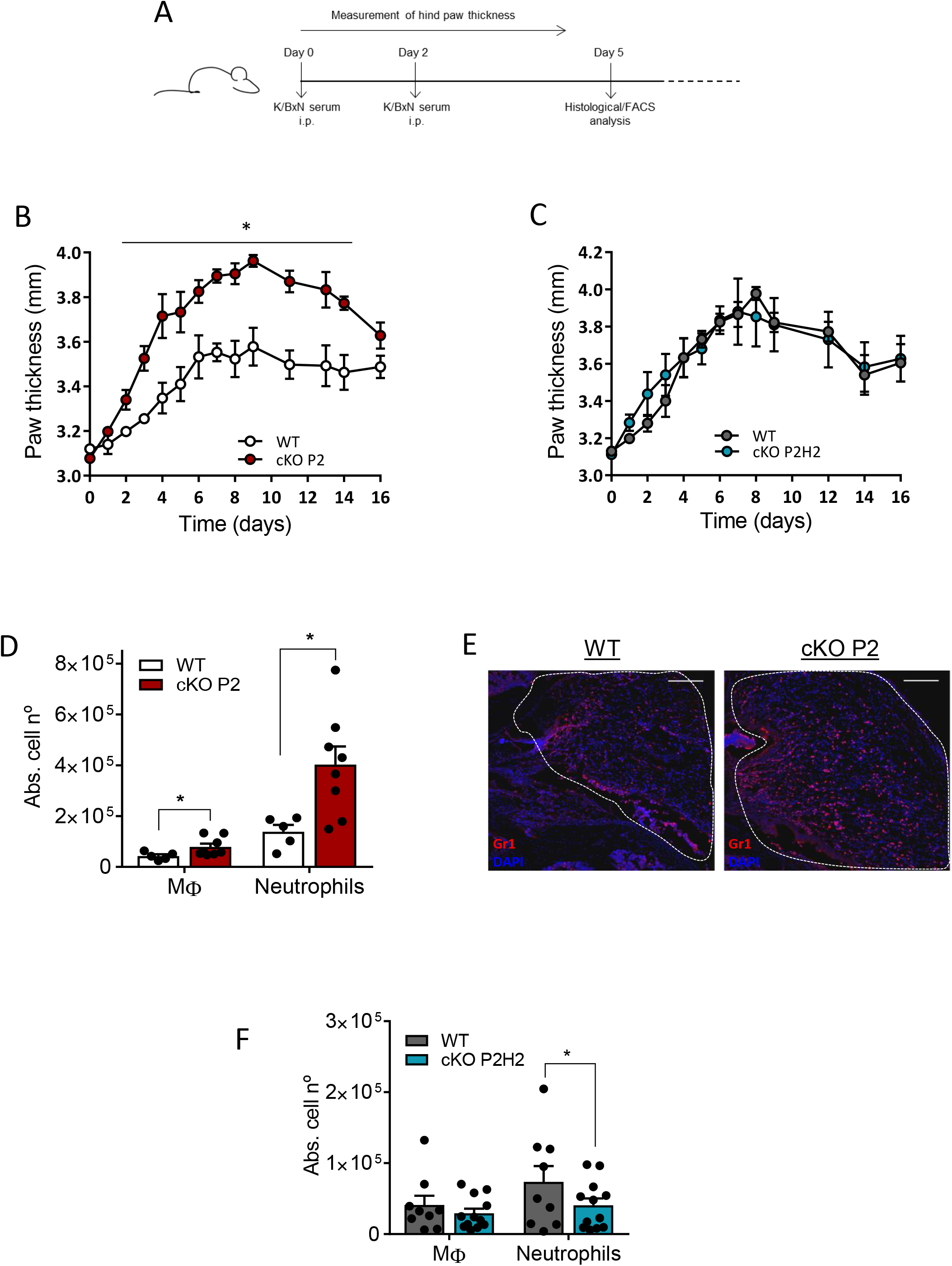

## Discussion

In the current work we have explored if hypoxia pathway proteins can directly regulate neutrophil motility, and reveal that activation of HIF2α in mouse neutrophils due to constitutive PHD2 loss enhances neutrophil migration through very confined environments independent of chemotactic -, glycolytic-or apoptotic-activity. Using a combination of *in vivo, ex vivo* and deep sequencing approaches, we provide evidence that these neutrophils have the capacity to migrate faster than their WT counterparts, and that this phenotype may be directly related to changes in their cytoskeleton mediated by a substantial reduction in RhoA GTPase activity.

Although it is generally accepted that neutrophils are the first immune cells to arrive in the tissue during inflammation, the molecular basis of neutrophil recruitment, which encompasses extravasation and interstitial migration, remain elusive. Further, neutrophil recruitment has been evaluated using a variety of migration assays in multiple studies related to the innate immune response,^53–56^ including in the context of hypoxia pathway proteins,^2,4^ but these studies call into debate the role of adhesion molecules.^57,58^ Here, we consistently show in 1D, 2D and 3D assays that neutrophils lacking PHD2 alone, and not both PHD2 and HIF2α, display enhanced cell motility and that only in severely confined environments. This difference in chemokinesis between cKO P2 and WT remained in a comparable setup using a chemokine as attractant (chemotaxis), demonstrating that the enhanced migratory capacity regulated by the PHD2-HIF2α axis is probably a cell intrinsic characteristic.

An important process during neutrophil recruitment is the final and time-limiting step of trans-endothelial migration (TEM), which is, in part, mediated by mechanical forces generated by the migrating neutrophil itself.^59–61^ We reveal a central role for HIF2α in this process. Indeed, considering the narrow pores between neighboring endothelial cells during the early phase of neutrophil diapedesis,^60^ our results from multiple approaches reiterate two main observations, viz., that greater numbers of cKO P2 neutrophils pass through small constrictions with enhanced speed. Intuitively, these observations account for the shorter TEM-time in a local ear inflammation model. The cumulative effects of such enhanced transmigration into inflamed tissues that were observable even at later time points in two completely independent *in vivo* models, i.e., inflammatory skin lesions and sterile arthritis. Indeed, it is possible that the enormous increase in cKO P2 neutrophils is positively affected by the fact that once a pore is opened, successive neutrophils are more likely to extravasate at this spot, enabling more neutrophils to enter the interstitium (skin) or the synovium (joint) of the inflamed tissue.^60^

Previous studies in a model of acute lung injury have reported enhanced glycolytic capacity of PHD2-deficient neutrophils, potentially due to HIF1α stabilization, which also enhanced neutrophil recruitment to the inflammatory site.^36^ Here, we confirm enhanced glycolysis in cKO P2 neutrophils and show that it is HIF2α-independent, strongly suggesting that glycolytic metabolism does not underlie the chemokinesis phenotype described here. The absence of differences in neutrophil apoptosis *in vivo* was corroborated by the RNAseq data from both single and double knock-out neutrophils, implying that the prolonged inflammation phenotype in cKO P2 mice was probably not due to persistence of the neutrophils. This is in contrast to results obtained using *in vitro* approaches that describe reduced apoptosis in HIF2α overexpressing neutrophils, which then resulted in delayed resolution of the inflammatory response.^4^ This group also reported delayed apoptosis in PHD2-deficient neutrophils and connected this to persistent inflammation.^36^ We believe these discrepancies are related to differences in the experimental models used.

Although several studies have linked the hypoxia pathway to the migratory capacity of a cell, only a few have suggested a role for the PHD/HIF axis in regulating cell migration through changes in cytoskeletal function.^20–22^ In migrating neutrophils *in vivo,* dynamic polymerized actin converges at the leading-edge of pseudopods, while stable actin with high acto-myosin contractility assemble at the rear. Both polarization and maintenance of this cytoskeletal asymmetry strongly rely on Rho GTPase activity.^62,63^ Using deep sequencing data from neutrophils of single and double transgenic lines, we show that a vast number of genes associated with Rho GTPase signaling are either directly or indirectly regulated by HIF2α. Interestingly, cKO P2 neutrophils displayed a significant downregulation of RhoA GTPase and we show this to be directly associated with enhanced motility because RhoA-inhibitor treated WT neutrophils behaved similarly in confined environments. These findings are similar to those reported earlier, i.e., increased flux of RhoA-deficient neutrophils and aggravated tissue injury in LPS-induced acute lung injury.^64^ A potential explanation is that the partial RhoA inhibition would primarily decrease dynamic cell protrusions, known to restrict cell migration by competing with stable actin cables at the cell rear.^18^ Alternatively, the HIF2α axis could be directly involved in the induction of cell contractility, which promotes neutrophil and DCs migration under strong confinement.^17,34^ However, further efforts are required to identify the specific molecular mechanism.

In conclusion, our results demonstrate that HIF2α-activation, due to constitutive loss of PHD-2, enhances the motility of neutrophils in highly confined surroundings, also during inflammation. Importantly, this phenotype is independent of chemotaxis signaling, glycolysis or apoptosis. Mechanistically, it is the reduction of RhoA GTPase activity that enhances the motility of PHD2 deficient neutrophils through very confined microenvironments. These findings highlight the potential deleterious effects of sustained HIF2α activity and may have important implications for the uncontrolled use of hypoxia mimetic agents that are currently licensed or are in phase II and III clinical trials.

## Acknowledgments

S.S. received financial support from the Dresden International Graduate School for Biomedicine and Bioengineering (DIGS-BB), B.W. was supported by the Heisenberg program (Deutsche Forschungsgemeinschaft – DFG, Germany; WI3291/5-1 and 12-1). This work was supported by grants from the DFG (TRR-CRC 205 Die Nebenniere: Zentrales Relais in Gesundheit und Krankheit (A02) to B.W. and T.C.; CRC 1181 (C7) to T.C.; the Alexander von Humboldt Foundation (AvH Professorship to J.G.). PV received financial support from the Association Nationale pour la Recherche (MOTILE project, ANR-16-CE13-0009), the Emergences Canceropole (SYNTEC project) and Labex-IPGG, as well as from “Institut Pierre-Gilles de Gennes” (laboratoire d’excellence, “Investissements d’avenir” program ANR-10-IDEX-0001-02 PSL and ANR-10-LABX-31). We would like to thank Silke Tulok and Dr. Anja Nobst from Core Facility Cellular Imaging (CFCI-MTZ-Dresden) for excellent assistance, Dr. Graf (CRG, Barcelona, Spain) for the Vav:cre mouse line and Dr. Vasuprada Iyengar for English Language and content editing.

## Authorship

S.S. designed the study, performed the majority of experiments, analysed data, and contributed in writing the manuscript. M.D., I.K., K.F. and M.K. designed and performed experiments, analysed data and contributed to the discussions. P.J.S., GLL and MB analysed data and contributed to the discussions. A.S. performed deep sequencing analysis and A.P. performed lipidomic analysis. M.G. performed intravital microscopy and contributed to the discussions.

A.K. and A.M. performed experiments and analysed data. J.G., Ü.C. provided tools and contributed to the discussions. T.C. provided tools, contributed to the discussions and edited the manuscript. P.V. designed and supervised the ex vivo migration studies, performed experiments, analysed data, and contributed in writing the manuscript. B.W. designed and supervised the overall study, analysed data, and wrote the manuscript.

## Current affiliation

I.K.: Hull York Medical School, York Biomedical Research Institute, University of York, York, United Kingdom. M.K. and J.G.: Max Planck Institute for the Science of Light & Max-Planck-Zentrum für Physik und Medizin, 91058 Erlangen, Germany.

