## Supplemental data, figures, Table for "HIF2α is a Direct Regulator of Neutrophil Motility"

### Supplemental Figure 1

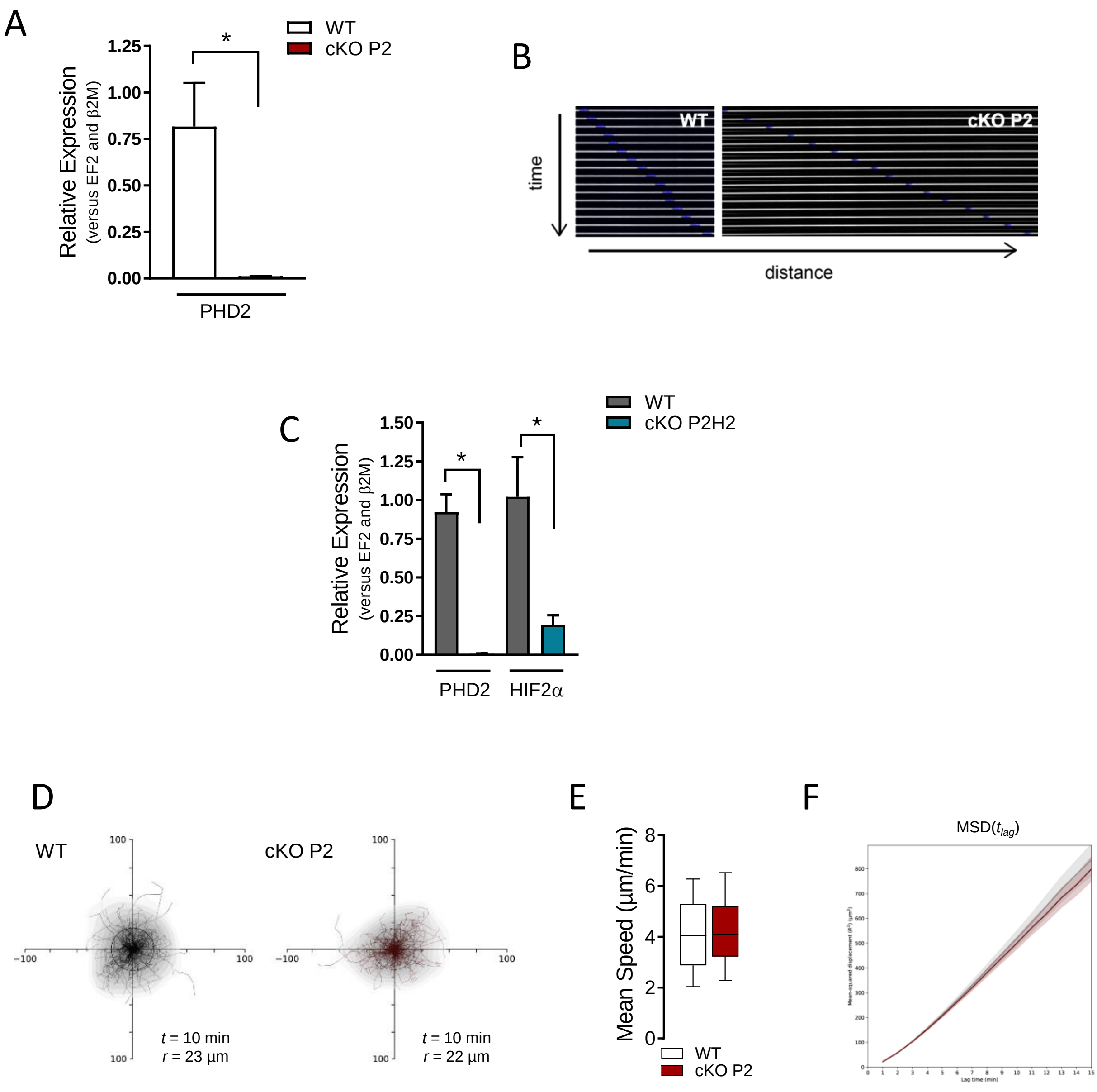

**Supplemental Figure 1. PHD2- and PHD2/HIF2-deficient neutrophils migrate differently in confined 2D environments.** (A) QPCR analysis of PHD2-deficient neutrophils confirmed significant downregulation of PHD2 compared to WT counterparts (n=3). (B) A representative montage of a single neutrophil (nucleus labeled with Hoechst) migrating through a 1D micro-channel of 3µm width. (C) QPCR analysis of PHD2/HIF2α-deficient neutrophils demonstrated significant downregulation of both genes compared to WT counterparts (n=3). (D) Representative neutrophil tracks during un-confined random 2D-migration (12 µm height). *t* indicates the period of the tracking, and *r* the mean displacement radius for this period. (E) Mean speed of neutrophil migration tracked in Panel D. (F) Mean squared displacement of neutrophil data depicted in Panel E.

Supplemental Figure 2

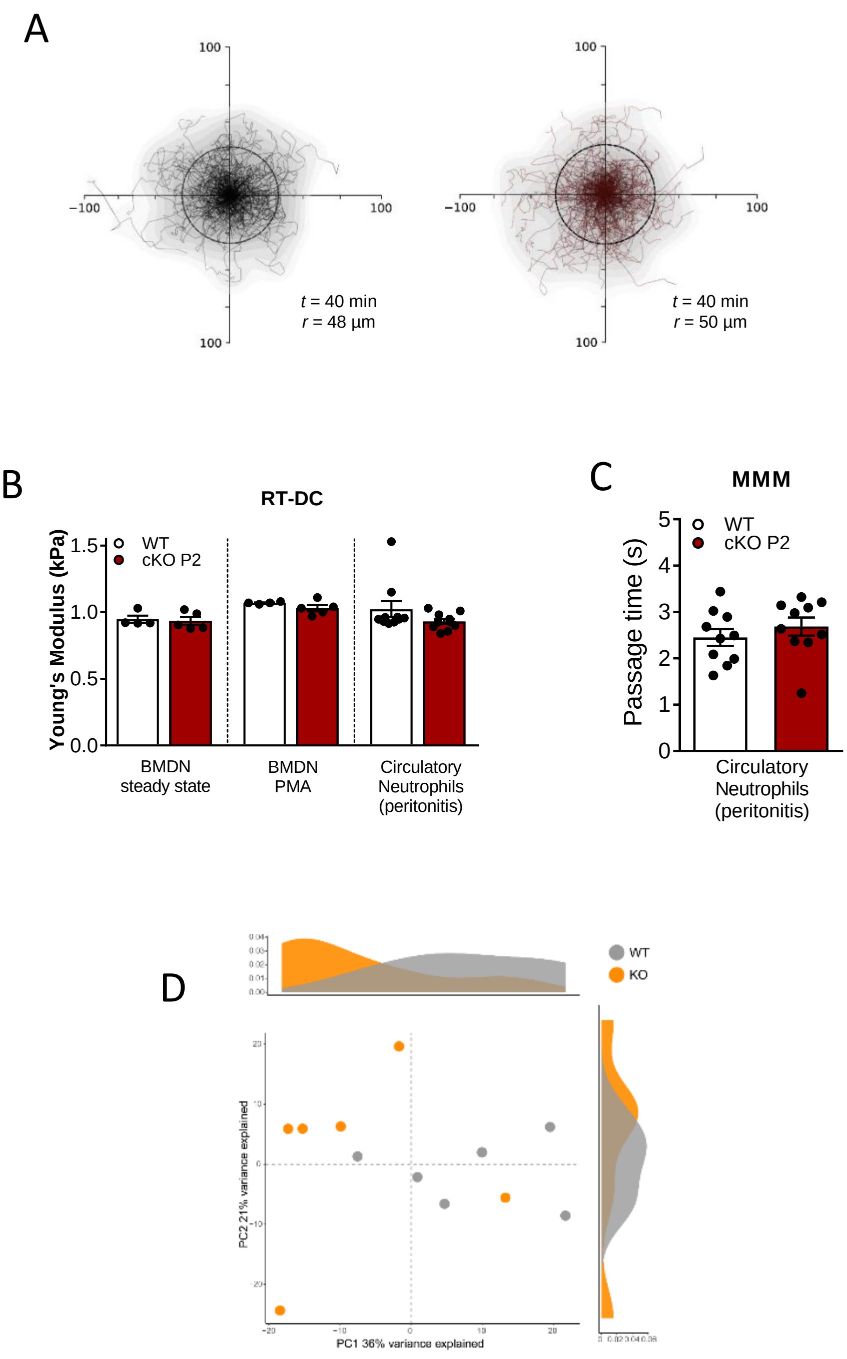

**Supplemental Figure 2. Loss of PHD2 does not affect neutrophil deformability under shear stress conditions or their lipid composition.**

(A) Representative neutrophil tracks during random 3D-migration in a 2mg/ml collagen gel. *t* indicates the period of the tracking and *r* the mean displacement radius for this period. (B) Real-time fluorescence and deformability cytometry (RT-C) was performed using a laminar flow of 0.08µl/s. Ly6G+ neutrophil´s deformability was measured and the Young‘s Modulus (kPa) was extracted in untreated BMDNs (steady state), PMA-treated, and circulatory neutrophils at 4h after thioglycolate-induced peritonitis (n=4-10). (C) Microfluidic microcirculation mimetic (MMM) assay was performed under constant pressure drop of 50 mBar between inlet and outlet. Neutrophil passage time through 180 micro-constrictions smaller than the cell diameter was evaluated as passage time (s). Data are represented as mean ± SEM. Statistical significance was defined using the Mann-Whitney U test. Data points in the graphs represent individual mice (n=10).(D) Principal component analysis (PCA) of the lipid composition of neutrophils from cKO P2 mice versus their WT counterparts.

Supplemental Figure 3

A

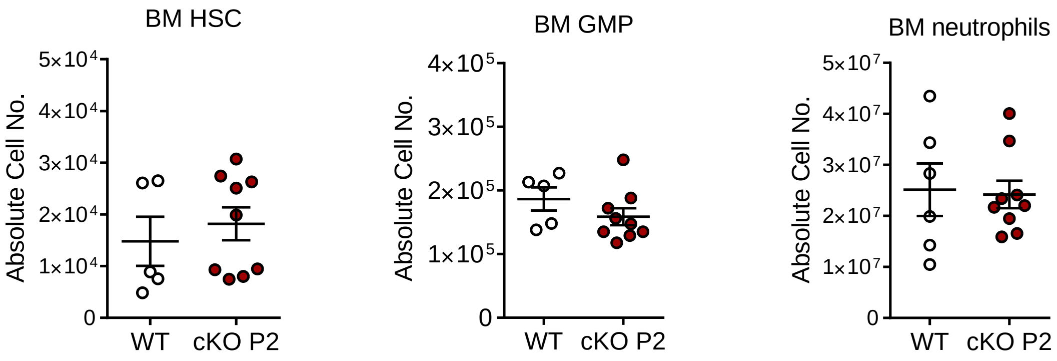

B

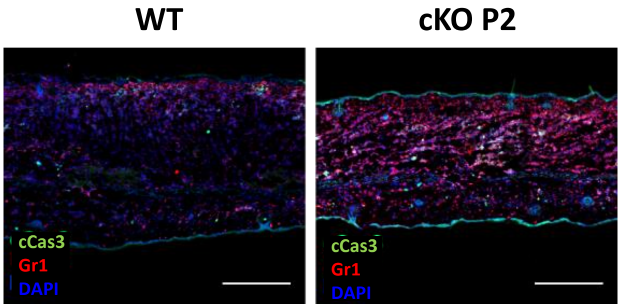

C

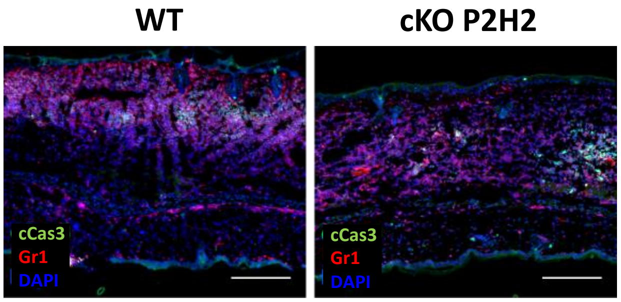

**Supplemental Figure 3. Loss of PHD2 in the hematopoietic compartment does not affect normal development of neutrophils or its progenitors.** Absolute cell numbers, enumerated through FACS analysis, for (A) hematopoietic stem cells (HSC), granulocyte-monocyte progenitors (GMP), or mature neutrophils from bone marrow (BM). In all graphs, data points represent individual mice (n=5-9). (B-C) Representative images of histological analysis of ear cryosections at 24 hours after PMA treatment that were stained for cleaved caspase 3 (cCas3) and Gr1 (neutrophils) in WT vs cKO P2 (B) and WT vs cKO P2H2 (C); all scale bars represent 50µm. Data are presented as mean ± SEM. Statistical significance was determined using Mann-Whitney U test; (*p<0.05).

### Supplemental Figure 4

A

| Regulon | NES | p-value | Amount of regulated genes |
|---|---|---|---|
| Slc2a13 | 3.4 | 0.0007 | 3448 |
| **Sgsm1** | **3.4** | **0.0007** | **3444** |
| **Cdc42bpg** | **3.4** | **0.0007** | **3444** |
| **Coro2b** | **3.31** | **0.0009** | **3444** |
| **Ggnbp2** | **3.14** | **0.0017** | **3620** |
| Xlr4b | 3.14 | 0.0017 | 3620 |
| Sh3bp5l | 3.09 | 0.002 | 4271 |
| Ormdl1 | 3.09 | 0.002 | 4271 |
| Zfp174 | 3.06 | 0.0022 | 3273 |
| Hist1h2bl | 3.06 | 0.0022 | 3273 |
| Cdkn2aip | 3.06 | 0.0022 | 3273 |
| Adgrg2 | 3.06 | 0.0022 | 3273 |
| Olfml1 | 3.03 | 0.0024 | 3348 |
| **Syt7** | **3.03** | **0.0025** | **3348** |
| Zfp958 | 3.01 | 0.0026 | 4237 |
| A830010M20Rik | 3.01 | 0.00264 | 3784 |
| **Avil** | **3.01** | **0.0027** | **3784** |
| Wif1 | 3 | 0.0027 | 4326 |
| **Spata13** | **3** | **0.0027** | **4326** |
| **Aldoc** | **3** | **0.0027** | **4326** |
| Tmem18 | 3 | 0.0027 | 4326 |
| Smarca1 | 3 | 0.0027 | 4326 |
| **Eps8l1** | **2.98** | **0.0029** | **3395** |
| **Eml5** | **2.94** | **0.0032** | **3920** |
| Zfp458 | 2.94 | 0.0033 | 3102 |
| **Enah** | **2.94** | **0.0033** | **3888** |
| Aff1 | 2.94 | 0.0033 | 3888 |
| 2610306M01Rik | 2.94 | 0.0033 | 3102 |
| Tcim | 2.94 | 0.0033 | 3888 |
| Zfp654 | 2.94 | 0.0033 | 3092 |
| Gm29243 | 2.93 | 0.0034 | 2971 |
| Epc1 | 2.93 | 0.0034 | 3960 |
| **Gad1** | **2.93** | **0.0034** | **2971** |
| Trmt12 | 2.93 | 0.0034 | 3960 |
| Mamld1 | 2.91 | 0.0036 | 3395 |
| C920021L13Rik | 2.91 | 0.0036 | 2888 |
| Mief1 | 2.91 | 0.0036 | 2888 |
| Prg4 | 2.91 | 0.0036 | 3395 |
| Wisp2 | 2.9 | 0.0037 | 2888 |
| Nrn1 | 2.89 | 0.0039 | 3172 |
| Fut1 | 2.89 | 0.0039 | 3858 |
| **Stard4** | **2.89** | **0.0039** | **3858** |
| Kcnf1 | 2.89 | 0.0039 | 3173 |
| Abraxas1 | 2.88 | 0.0039 | 2802 |
| Tnxb | 2.88 | 0.004 | 4132 |
| **Arhgef26** | **2.88** | **0.004** | **4132** |
| Clcf1 | 2.88 | 0.004 | 3309 |
| Zfp655 | 2.88 | 0.004 | 3102 |

| Regulon | NES | p.value | Amount of regulated genes |
|---|---|---|---|
| Izumo1r | -2.89 | 0.0039 | 3184 |
| Apod | -2.89 | 0.0039 | 3184 |
| Timm17b | -2.89 | 0.0039 | 3860 |
| Mms19 | -2.89 | 0.0039 | 3184 |
| Il15ra | -2.89 | 0.0039 | 3184 |
| **Gzma** | **-2.91** | **0.0037** | **3397** |
| Dlx1 | -2.91 | 0.0037 | 3397 |
| Lipt1 | -2.91 | 0.0037 | 3397 |
| **Ift81** | **-2.93** | 0.0037 | **2972** |
| Ndufs8 | -2.93 | 0.0037 | 2972 |
| Pex3 | -2.93 | 0.0037 | 3951 |
| S100a1 | -2.93 | 0.0037 | 3951 |
| Klrb1c | -2.94 | 0.0037 | 3937 |
| Tmem126a | -2.94 | 0.0037 | 3937 |
| Pofut2 | -2.94 | 0.0037 | 3937 |
| Phf11d | -2.94 | 0.0037 | 3055 |
| Fez2 | -2.94 | 0.0038 | 3055 |
| Dact3 | -2.94 | 0.0038 | 3913 |
| Rcan1 | -2.94 | 0.0038 | 3913 |
| Smim20 | -2.94 | 0.0038 | 3913 |
| Atp1b3 | -2.94 | 0.0038 | 3913 |
| Kxd1 | -2.94 | 0.0038 | 3913 |
| Rpl19 | -2.94 | 0.0038 | 3913 |
| Btbd2 | -2.94 | 0.0038 | 3913 |
| Lbh | -2.98 | 0.0038 | 3139 |
| Gm11944 | -2.98 | 0.0038 | 3139 |
| Lrp11 | -2.99 | 0.0039 | 3774 |
| Gns | -3 | 0.0039 | 3774 |
| Abhd12 | -3 | 0.0039 | 3774 |
| Lgals3bp | -3 | 0.0039 | 3774 |
| Ncs1 | -3.01 | 0.0039 | 4331 |
| Fam98c | -3.01 | 0.0039 | 4331 |
| Slc6a1 | -3.01 | 0.0039 | 3440 |
| P2ry6 | -3.01 | 0.0039 | 4331 |
| **Ctss** | **-3.01** | 0.0039 | **4331** |
| Tprn | -3.01 | 0.0039 | 4331 |
| Slc25a35 | -3.01 | 0.004 | 3440 |
| Ly86 | -3.01 | 0.004 | 4331 |
| Spaca9 | -3.01 | 0.004 | 4331 |
| Phf11b | -3.02 | 0.004 | 4331 |
| Hspb11 | -3.02 | 0.004 | 2922 |
| Klra9 | -3.03 | 0.004 | 3357 |
| Prf1 | -3.03 | 0.004 | 3357 |
| Il2rb | -3.03 | 0.004 | 3357 |
| Gm12185 | -3.03 | 0.004 | 3357 |
| Gnptg | -3.08 | 0.004 | 3774 |
| D230025D16Rik | -3.08 | 0.0041 | 3774 |
| Arl4d | -3.09 | 0.0041 | 4276 |

B

| Motif: HIF2α(bHLH)/785_O-HIF2α-ChIP-Seq(GSE34871) | | | |
|---|---|---|---|
| Name | Refseq | Sequence | MotifScore |
| Mylk | NM_139300 | TGGTACGTGC | 9,357758 |
| Myh10 | NM_175260 | AGAGACGTGA | 7,650727 |
| Arrb1 | NM_177231 | AAAGACGTGC | 7,280568 |
| Cav1 | NM_007616 | GCACGTCCTA | 7,62973 |
| Trim32 | NM_053084 | AAGGACGTGC | 7,806661 |
| Ptger3 | NM_011196 | TCACGTAACC | 7,256074 |
| Ttn | NM_011652 | TCACGTTTGC | 7,071815 |

**Supplemental Figure 4. RNAseq analysis**

(A) The most significant master regulators (pos. and neg.) in PHD2-deficient neutrophils that can potentially regulate the expression/activity of a vast number of genes or gene products (=amount of regulated genes). Genes highlighted in bold belong to cytoskeleton remodeling signatures. (B) Set of genes involved in cytoskeleton modelling and identified as harboring Hypoxia Responsive Elements (HRE) in their promoter/enhancer regions.

### Supplemental data

#### Transcriptome Mapping

Low quality nucleotides were removed using the Illumina Fastq filter (http://cancan.cshl.edu/labmembers/gordon/fastq_illumina_filter/). Reads were further subjected to adaptor trimming using cutadapt [1]. Alignment of the reads to the Mouse genome was done using STAR Aligner [2] with the parameters: "--runMode alignReads --outSAMstrandField intronMotif --outSAMtype BAM SortedByCoordinate --readFilesCommand zcat". Mouse Genome version GRCm38 (release M12 GENCODE) was used for alignment and average concordant mapping rates for the samples were between 91-96 %.

#### Read Quantification

The parameters: 'htseq-count -f bam -s reverse -m union -a 20', HTSeq-0.6.1p1 [3] were used to count the reads to be mapped to the genes in the aligned sample files. The same GTF file was used for mapping and read quantification.

#### Differential Expression Analysis

Gene-centric differential expression analysis was performed using DESeq2_1.8.1 [4]. The raw read counts for genes across samples were normalized using the 'rlog' command of DESeq2, and subsequently, these values were used to render a PCA plot using ggplot2_1.0.1 [5].

#### Functional Analyses

Pathway and functional analyses of the genes were performed using GSEA [6] and EGSEA [7]. GSEA is a stand-alone software with a GUI. To run GSEA, a ranked list of all the genes from DESeq2-based calculations was created using the -log10 of the p-value and multiplying it with the direction of fold change, i.e., '+' for positive or '-' for negative. This ranked list was then queried against the following repositories: Molecular Signatures Database (MSigDB), Reactome, KEGG, and GO (gene ontology).

EGSEA is an R/Bioconductor-based command-line package. For functional analyses using EGSEA, a differentially expressed list of genes with parameters, log2foldchange > 0.3 and padj <0.05, was used. Functional analyses used the same data repositories as above.

#### Master Regulators

The program ARACNE [8] was used to construct a gene network in the form of an adjacency list using normalized gene expression values as the input. This network, along with a normalized expression matrix, was further used as an input for the program VIPER [9], which uses these two inputs to compute a list of master expression regulators (genes).

#### Network Construction

A union of Protein-Protein Interactions (PPIs) for mouse was extracted from BioGrid [10] and STRING [11] databases. Genes from the GSEA pathway (cytoskeleton remodeling), the GTPases, RhoA and Cdc42, along with the master regulators involved in cytoskeleton remodeling, were mapped on to the PPI network. The network was constructed using Cytoscape v3.6 [12]. A degree-sorted, anticlockwise circular layout, starting from highest degree node/gene: Myh10, to the lowest: Spata13, was chosen to depict the interaction network. In this depiction, the nodes (genes) are colored according to their degree of connectivity, wherein highly connected nodes are purple while less well-connected nodes in lighter shades. The edges involving RhoA and Cdc42 have been darkened.

**Motif Enrichment Analysis**

HOMER (v4.10) [13] was used to detect the presence of Hypoxia Responsive Elements (HRE) in the promoter/enhancer regions of the genes involved in cytoskeleton modelling.

**Lipid extraction for mass spectrometry lipidomics**

BMDNs were obtained via positive selection and cell pellets (6 individual samples per genotype) were immediately frozen at -80 degrees C. Mass spectrometry-based lipid analysis was performed at Lipotype GmbH (Dresden, Germany) as previously described [14]. Lipids were extracted using a two-step chloroform/methanol procedure [15]. Samples were spiked with internal lipid standard mixture containing: cardiolipin 16:1/15:0/15:0/15:0 (CL), ceramide 18:1;2/17:0 (Cer), diacylglycerol 17:0/17:0 (DAG), hexosylceramide 18:1; 2/12:0 (HexCer), lyso-phosphatidate 17:0 (LPA), lyso-phosphatidylcholine 12:0 (LPC), lysophosphatidylethanolamine 17:1 (LPE), lyso-phosphatidylglycerol 17:1 (LPG), lyso-phosphatidylinositol 17:1 (LPI), lyso-phosphatidylserine 17:1 (LPS), phosphatidate 17:0/17:0 (PA), phosphatidylcholine 17:0/17:0 (PC), phosphatidylethanolamine 17:0/17:0 (PE), phosphatidylglycerol 17:0/17:0 (PG), phosphatidylinositol 16:0/16:0 (PI), phosphatidylserine 17:0/17:0 (PS), cholesterol ester 20:0 (CE), sphingomyelin 18:1;2/12:0;0 (SM), triacylglycerol 17:0/17:0/17:0 (TAG) and cholesterol D6 (Chol). After extraction, the organic phase was transferred to an infusion plate and dried in a speed vacuum concentrator. The 1$^{st}$ step dry extract was re-suspended in 7.5 mM ammonium acetate in chloroform/methanol/propanol (1:2:4, V:V:V) while the 2$^{nd}$ step dry extract was in 33% ethanol solution of methylamine in chloroform/methanol (0.003:5:1; V:V:V). All liquid handling steps were performed using Hamilton Robotics STARlet robotic platform with the Anti Droplet Control feature for organic solvent pipetting.

MS data acquisition - Samples were analyzed by direct infusion on a QExactive mass spectrometer (Thermo Scientific) equipped with a TriVersa NanoMate ion source (Advion Biosciences). Samples were analyzed in both positive and negative ion modes with a resolution of $R_{m/z=200}=280000$ for MS and $R_{m/z=200}=17500$ for MS-MS experiments, in a single acquisition. MS-MS was triggered by an inclusion list encompassing corresponding MS mass ranges scanned in 1 Da increments [16]. Both MS and MS-MS data were combined to monitor CE, DAG and TAG ions as ammonium adducts; PC, PC O-, as acetate adducts; and CL, PA, PE, PE O-, PG, PI and PS as deprotonated anions. MS-only was used to monitor LPA, LPE, LPE O-, LPI and LPS as deprotonated anions; Cer, HexCer, SM, LPC and LPC O- as acetate adducts, and cholesterol as ammonium adduct of an acetylated derivative [17].

Data analysis and post-processing were conducted with in-house developed lipid identification software based on LipidXplorer [18, 19]. Post-processing and normalization of data were performed using an in-house developed data management system. Only lipid identifications with a signal-to-noise ratio >5, and a signal intensity 5-fold higher than in corresponding blank samples were considered for further data analysis.

Lipidomics data downstream analysis: The data matrix provided by Lipotype GmbH contained 633 different lipids measured across 12 samples (picomol/microgram). The dataset was filtered by removing all lipids that were measured in a single sample (n= 86). The original dataset, its filtered version, and the subset of lipids removed by filtering are provided in Suppl. ”Lipidomics_data_pmol.xlsx”. The filtered data was log-transformed before undergoing Principal Component Analysis (scaled and centered PCA). Variations between the two conditions (wild type versus knockout) were statistically tested by means of the Wilcoxon rank sum test, which was calculated for lipid species, lipid classes, lipids regrouped by their total chain length, and lipids regrouped by the number of double bonds. Resulting p-values were adjusted according to the Benjamini-Hochberg correction. Analyses were performed in R (ver. 3.6.1) [20] and plots were generated using ggplot2 [21].

**Intravital microscopy**

Ear inflammation was initiated (10µl of 1mg/ml PMA diluted in ethanol) 3h before the actual start of imaging. Mice were anaesthetized with 3% isoflurane, endotracheally intubated, and actively ventilated for anesthesia maintenance with 2% isoflurane. The left ear was gently immobilized on a heat-regulated glass platform via double-sided tape and covered in sterile eye-gel (Vidisic, Bausch+Lomb) for objective immersion. Anti-Ly6G-PE and FITC-Dextran (MW 150.000Da) (2.5mg/ml, Sigma-Aldrich) were simultaneously injected via a tail vein catheter to label neutrophils and the vasculature, respectively.

Imaging was performed with an upright Leica SP8 multiphoton laser scanning microscope at the Core Facility Cellular Imaging Dresden (CFCI). A Chamaeleon II (Coherent) was used as laser emitter, while non-descanned HyD and PMT detectors (Leica) were utilized for signal collection. 3D images (Z-stacks) in the 70-100µm range were acquired using a 25x 0.95NA water immersion objective with long working distance (Leica). Pixel size corresponded to 0.433/0.433/2.0µm (X/Y/Z). For imaging, 3 Positions were repeatedly imaged for a duration of 30 min (8 frames each, interleaved acquisition) from 3 hours after stimulation.

Imaging data were acquired by simultaneous excitation/detection with a laser wavelength of 960nm. FITC-Dextran (500-550nm) and PE signal (580-654nm) were detected with HyD detectors (Leica). Second harmonic generation signal (415-474nm) was detected with the PMT detector (Leica).

**Real-Time Deformability Assay (RT-DC)**

RT-DC was performed as previously described [22,23] using steady state BMDNs, BMDNs previously treated with PMA (200ng/ml) for 30 minutes, and peripheral blood neutrophils at 4h after Thioglycolate-induced sterile peritonitis. Cells were resuspended in 0.6% methylcellulose containing PBS and flushed through a narrow 20 µm wide and 20 µ high channel constriction. Cell deformability was recorded using high speed microscopy at a constant flow rate of 0.08µl/s. Prior to the measurement all isolated neutrophils were labelled with APC-conjugated anti-Ly6G antibody to exclusively evaluate only Ly6G+ cells. Data analysis was performed using ShapeOut software (Zellmechanik Dresden) and Young´s Modulus of every single cell was extracted using an analytical model[24] and numerical simulations[25].

**Microfluidic Microcirculation Mimetic (MMM) Assay**

MMM was done as described previously[26,27]. In brief, MMM is based on a microfluidic chip made from PDMS. An inlet and an outlet region connected by a single channel (20 µm in width) with 180 successive constrictions (5 µm in width). Peripheral blood neutrophils isolated at 4h after Thioglycolate-induced sterile peritonitis were resuspended to a final density of $3 \times 10^4$ cells in 1 ml in PBS and driven through the constriction at a constant pressure drop between inlet and outlet of 50 mBar using an air pressure control system (MFCS-FLEX; Fluigent, Villejuif, France). The microfluidic chip was mounted to an inverted microscope, which was connected to a camera (The Imaging Source, Bremen, Germany) to record videos at a frame-rate of 120 frames $sec^{-1}$. The total passage time was extracted from the videos using custom-made written codes in Python.

**TABLE 1.** LIST OF PRIMERS USED TO GENOTYPE MICE

| | |
|---|---|
| Vav_1 | 5´-CCATGGCACCCAAGAAGAAG-3´ |
| Vav_2 | 5´-GCTTAGTTTTCCTGCAGCGG-3´ |
| PHD2, exon2 | 5’-CGCATCTTCCATCTCCATTT-3’ |
| PHD2, intron3 | 5’-GGCAGTGATAACAGGTGCAA-3’ |
| PHD2, intron1 | 5’-CTCACTGACCTACGCCGTGT-3’ |
| HIF2α, intron2 | 5’-CAGGCAGTATGCCTGGCTAATTCCAGTT-3’ |
| HIF2α, intron2 | 5'-CTTCTTCCATCATCTGGGATCTGGGACT-3' |
| HIF2α, intron1 | 5’-GCTAACACT GTACTGTCTGAAAGAGTAGC-3’ |

**TABLE 2.** LIST OF ANTIBODIES

| **Primary Antibodies** | | | | | |
|---|---|---|---|---|---|
| **Antibody** | **Dilution** | **Specificity** | **Host** | **Catalog number** | **Company** |
| Gr1 | 1:200 | mouse | rat | 14-5931-81 | Thermo Fischer |
| Cleaved Caspase 3 (cCas3) | 1:300 | mouse | rabbit | 9661S | Cell signalling |
| CD45 PE | 1:200 | mouse | rat | 12-0451-82 | eBioscience |
| CD11b FITC | 1:800 | mouse | rat | 11-0112-81 | eBioscience |
| F4/80 PE-Cy7 | 1:100 | mouse | rat | 25-4801-82 | eBioscience |
| Ly6G APC | 1:200 | mouse | rat | 17-9668-80 | eBioscence |
| Ly6G biotin | 1:800 | mouse | rat | 13-5931-82 | eBioscience |
| **Secondary Antibodies** | | | | | |
| **Antibody** | **Dilution** | **Specificity** | **Host** | **Catalog number** | **Company** |
| Anti-Rat-IgG A555 | 1:500 | rat | goat | 4417 | Cell signalling |
| Anti-Rabbit-IgG A488 | 1:500 | rabbit | goat | 4412 | Cell signalling |

| | class | functional category | total length | totaldb | totaloh | 205wt | 632wt | 634wt | 980wt | 981wt | 635wt | 203ko | 479ko | 480ko | 824ko | 874ko | 875ko | |
|---|---|---|---|---|---|---|---|---|---|---|---|---|---|---|---|---|---|---|
| CE_16:0 | CE | STO | 16 | 0 | 0 | 8,08737 | 8,04288 | 9,56237 | 3,71014 | 7,96068 | 7,03083 | 5,36145 | 4,89195 | 10,345 | 6,20611 | 5,41328 | 0 | CE_16:0 |
| CE_16:1 | CE | STO | 16 | 1 | 0 | 0 | 4,12311 | 5,38192 | 0 | 0 | 0 | 0 | 0 | 0 | 0 | 2,97626 | 0 | CE_16:1 |
| CE_18:1 | CE | STO | 18 | 1 | 0 | 11,9491 | 10,1119 | 11,2602 | 6,87398 | 11,4341 | 11,3181 | 8,64011 | 6,88753 | 13,1547 | 9,15929 | 8,28613 | 7,52029 | CE_18:1 |
| CE_18:2 | CE | STO | 18 | 2 | 0 | 4,21073 | 4,6784 | 0 | 0 | 4,533 | 0 | 0 | 3,53417 | 3,56836 | 0 | 0 | 0 | CE_18:2 |
| Cer_34:0;2 | Cer | MEM | 34 | 0 | 2 | 0,280019 | 0,243851 | 0,361386 | 0,151278 | 0,217711 | 0,34586 | 0,159043 | 0,132746 | 0,21574 | 0,238347 | 0,166654 | 0,167995 | Cer_34:0;2 |
| Cer_34:2;2 | Cer | MEM | 34 | 2 | 2 | 0,170626 | 0 | 0,170415 | 0,081007 | 0,139043 | 0,16263 | 0,087226 | 0 | 0 | 0,149324 | 0 | 0,081283 | Cer_34:2;2 |
| Cer_36:1;2 | Cer | MEM | 36 | 1 | 2 | 0,767257 | 0,718147 | 0,877771 | 0 | 0,64303 | 0,877429 | 0 | 0 | 0 | 0,825154 | 0 | 0 | Cer_36:1;2 |
| Cer_36:2;2 | Cer | MEM | 36 | 2 | 2 | 0 | 0,089125 | 0 | 0 | 0 | 0,136558 | 0 | 0 | 0 | 0 | 0 | 0 | Cer_36:2;2 |
| Cer_38:1;2 | Cer | MEM | 38 | 1 | 2 | 0,330736 | 0,260558 | 0,288986 | 0 | 0,266419 | 0,329284 | 0 | 0 | 0 | 0,312174 | 0 | 0 | Cer_38:1;2 |
| Cer_40:1;2 | Cer | MEM | 40 | 1 | 2 | 1,487 | 1,26371 | 1,57997 | 0,989208 | 1,33238 | 1,3707 | 0,768301 | 0,851495 | 1,37775 | 1,51125 | 0,919427 | 0,930481 | Cer_40:1;2 |
| Cer_40:2;2 | Cer | MEM | 40 | 2 | 2 | 0,302791 | 0,256762 | 0,295167 | 0,204427 | 0,261572 | 0,321412 | 0,193249 | 0,161479 | 0,161702 | 0,285758 | 0,124116 | 0,218418 | Cer_40:2;2 |
| Cer_42:0;1 | Cer | MEM | 42 | 0 | 1 | 0 | 0,111905 | 0,109947 | 0 | 0 | 0 | 0,099156 | 0 | 0,147809 | 0 | 0,096451 | 0 | Cer_42:0;1 |
| Cer_42:1;1 | Cer | MEM | 42 | 1 | 1 | 0,223948 | 0,194348 | 0,230997 | 0,147674 | 0,186291 | 0,119523 | 0,201961 | 0,123762 | 0,241721 | 0,198499 | 0,169975 | 0 | Cer_42:1;1 |
| Cer_42:1;2 | Cer | MEM | 42 | 1 | 2 | 3,1231 | 2,62213 | 3,80978 | 1,91136 | 2,67489 | 2,90709 | 2,17416 | 1,65393 | 2,52852 | 3,10277 | 1,72875 | 1,52404 | Cer_42:1;2 |
| Cer_42:2;1 | Cer | MEM | 42 | 2 | 1 | 0,096688 | 0,091807 | 0 | 0 | 0 | 0 | 0 | 0 | 0 | 0 | 0,095256 | 0 | Cer_42:2;1 |
| Cer_42:2;2 | Cer | MEM | 42 | 2 | 2 | 6,45272 | 5,50003 | 6,89831 | 4,44999 | 5,73904 | 6,21874 | 3,593 | 3,42876 | 4,442 | 6,39024 | 3,41437 | 3,71789 | Cer_42:2;2 |
| Chol | Chol | MEM | 27 | 1 | 0 | 882,539 | 892,966 | 1039,9 | 685,422 | 823,72 | 958,42 | 636,771 | 624,349 | 831,967 | 904,306 | 541,57 | 542,006 | Chol |
| CL_68:2 | CL | MEM | 68 | 2 | 0 | 0 | 0,248116 | 0,693169 | 0,499814 | 0,619435 | 0,720866 | 0,701935 | 0,80386 | 0 | 1,0394 | 0 | 0,416004 | CL_68:2 |
| CL_70:2 | CL | MEM | 70 | 2 | 0 | 1,29367 | 0,522866 | 0 | 0,726732 | 0,320084 | 1,91855 | 0,799931 | 0,7228 | 0 | 2,43398 | 1,75555 | 0,283551 | CL_70:2 |
| CL_70:4 | CL | MEM | 70 | 4 | 0 | 0 | 0,584094 | 1,10609 | 0 | 0 | 1,33798 | 0,785897 | 0 | 0 | 0,975488 | 0 | 0,267792 | CL_70:4 |
| CL_70:5 | CL | MEM | 70 | 5 | 0 | 0 | 0,358112 | 1,28652 | 0,370554 | 0 | 1,18236 | 0,580767 | 0 | 0 | 0,854512 | 0 | 0 | CL_70:5 |
| CL_70:6 | CL | MEM | 70 | 6 | 0 | 0 | 0,335122 | 1,60574 | 0 | 0 | 1,50947 | 0,582578 | 0 | 0 | 0,778276 | 0 | 0 | CL_70:6 |
| CL_70:7 | CL | MEM | 70 | 7 | 0 | 0 | 0,662965 | 1,12076 | 0,717959 | 0,536445 | 1,14433 | 0,64769 | 0 | 0 | 1,36362 | 0 | 0,600516 | CL_70:7 |
| CL_72:6 | CL | MEM | 72 | 6 | 0 | 3,93311 | 1,8772 | 3,77247 | 1,67438 | 1,36657 | 5,06944 | 2,77055 | 1,52633 | 0 | 3,90162 | 2,66653 | 1,08479 | CL_72:6 |
| CL_72:7 | CL | MEM | 72 | 7 | 0 | 6,07884 | 3,2573 | 6,01018 | 3,04255 | 3,30365 | 6,5729 | 2,92285 | 2,22344 | 0 | 7,10365 | 5,06566 | 2,36446 | CL_72:7 |
| CL_72:8 | CL | MEM | 72 | 8 | 0 | 5,14721 | 3,56667 | 5,92782 | 3,75534 | 3,39608 | 6,24397 | 2,54276 | 3,06786 | 0 | 8,27565 | 6,44473 | 3,02897 | CL_72:8 |
| CL_74:6 | CL | MEM | 74 | 6 | 0 | 7,16925 | 2,29729 | 6,52778 | 2,73855 | 1,99986 | 5,72078 | 4,53859 | 3,25442 | 0 | 10,7487 | 7,34457 | 0,753239 | CL_74:6 |
| CL_74:8 | CL | MEM | 74 | 8 | 0 | 1,7825 | 1,66495 | 2,92006 | 1,36534 | 1,42336 | 3,28782 | 1,27356 | 0 | 0 | 2,87245 | 2,40561 | 0,87738 | CL_74:8 |
| CL_74:9 | CL | MEM | 74 | 9 | 0 | 0 | 1,18065 | 1,64716 | 0 | 0,959347 | 0 | 0 | 0 | 0 | 1,21842 | 0 | 0,587475 | CL_74:9 |
| DAG_16:0-18:1 | DAG | MEM | 34 | 1 | 0 | 5,45518 | 4,25363 | 5,08834 | 3,7489 | 4,61363 | 6,45917 | 4,66171 | 2,83475 | 0 | 4,47026 | 2,68693 | 2,77517 | DAG_16:0-18:1 |
| DAG_16:0-18:2 | DAG | MEM | 34 | 2 | 0 | 2,94586 | 2,52775 | 2,30188 | 1,96765 | 2,36579 | 3,27588 | 2,57843 | 1,86197 | 1,31309 | 2,54984 | 0 | 1,59609 | DAG_16:0-18:2 |
| DAG_16:0-20:2 | DAG | MEM | 36 | 2 | 0 | 0,283339 | 0 | 0,194233 | 0,067941 | 0,181918 | 0,290575 | 0,13861 | 0 | 0 | 0,218189 | 0 | 0 | DAG_16:0-20:2 |
| DAG_16:0-20:3 | DAG | MEM | 36 | 3 | 0 | 0,381326 | 0,256187 | 0,365607 | 0,173654 | 0,317494 | 0,523365 | 0 | 0 | 0 | 0,273704 | 0,144378 | 0 | DAG_16:0-20:3 |
| DAG_16:0-20:4 | DAG | MEM | 36 | 4 | 0 | 1,10295 | 1,4743 | 1,11726 | 1,00498 | 1,54638 | 1,49718 | 2,28653 | 0 | 0 | 1,25087 | 0,893836 | 0 | DAG_16:0-20:4 |
| DAG_16:0-22:4 | DAG | MEM | 38 | 4 | 0 | 0,350571 | 0,175227 | 0,359485 | 0 | 0,390054 | 0,533773 | 0,383113 | 0 | 0 | 0,31218 | 0,157027 | 0 | DAG_16:0-22:4 |
| DAG_16:1-18:0 | DAG | MEM | 34 | 1 | 0 | 0,122853 | 0,096084 | 0,171144 | 0 | 0 | 0,150081 | 0,081142 | 0 | 0 | 0,137991 | 0 | 0 | DAG_16:1-18:0 |
| DAG_16:1-18:1 | DAG | MEM | 34 | 2 | 0 | 0,502792 | 0,495351 | 0,517893 | 0,337892 | 0,399799 | 0,587559 | 0,500404 | 0,37776 | 0,198406 | 0,537753 | 0 | 0,251212 | DAG_16:1-18:1 |
| DAG_18:0-18:2 | DAG | MEM | 36 | 2 | 0 | 1,51889 | 0 | 1,06502 | 0,783693 | 1,14702 | 1,35756 | 0,649292 | 0 | 0 | 1,19964 | 0,749805 | 0 | DAG_18:0-18:2 |
| DAG_18:0-20:4 | DAG | MEM | 38 | 4 | 0 | 3,80201 | 3,31696 | 3,98185 | 2,67531 | 3,85247 | 4,58981 | 3,1726 | 2,00173 | 0 | 3,61266 | 1,7633 | 2,18386 | DAG_18:0-20:4 |
| DAG_18:1-18:1 | DAG | MEM | 36 | 2 | 0 | 1,30229 | 0 | 1,3153 | 0,918604 | 1,31096 | 1,48113 | 0,776251 | 0 | 0 | 1,23663 | 0,888817 | 0 | DAG_18:1-18:1 |
| DAG_18:1-18:2 | DAG | MEM | 36 | 3 | 0 | 1,48365 | 1,3977 | 2,37912 | 1,41561 | 2,09573 | 2,28386 | 0 | 1,4422 | 0 | 1,46047 | 1,41084 | 0 | DAG_18:1-18:2 |
| DAG_18:1-18:3 | DAG | MEM | 36 | 4 | 0 | 0,295024 | 0 | 0,943338 | 0 | 0 | 0,441538 | 0 | 0 | 0 | 0 | 0 | 0 | DAG_18:1-18:3 |
| DAG_18:1-20:3 | DAG | MEM | 38 | 4 | 0 | 0 | 0 | 0,148717 | 0 | 0,207445 | 0 | 0 | 0 | 0 | 0,146777 | 0 | 0 | DAG_18:1-20:3 |
| DAG_18:2-18:2 | DAG | MEM | 36 | 4 | 0 | 0,389521 | 0,60658 | 0,565946 | 0,590461 | 0,576925 | 0,481091 | 0,597354 | 0 | 0 | 0,438999 | 0,531162 | 0 | DAG_18:2-18:2 |
| DAG_18:2-20:2 | DAG | MEM | 38 | 4 | 0 | 0 | 0 | 0,111 | 0 | 0,119923 | 0 | 0 | 0 | 0 | 0 | 0 | 0 | DAG_18:2-20:2 |
| HexCer_34:1;2 | HexCer | MEM | 34 | 1 | 2 | 2,47358 | 2,75506 | 2,62985 | 1,73746 | 1,91034 | 2,14716 | 1,26557 | 1,71972 | 2,22921 | 2,37816 | 1,35189 | 1,57475 | HexCer_34:1;2 |
| HexCer_36:1;2 | HexCer | MEM | 36 | 1 | 2 | 0,419814 | 0 | 0,384955 | 0 | 0,242471 | 0 | 0 | 0,23051 | 0 | 0,3887 | 0 | 0 | HexCer_36:1;2 |
| HexCer_38:1;2 | HexCer | MEM | 38 | 1 | 2 | 0,359424 | 0 | 0,296181 | 0,232665 | 0,269884 | 0,479867 | 0 | 0,28673 | 0 | 0,296714 | 0 | 0 | HexCer_38:1;2 |
| HexCer_40:1;2 | HexCer | MEM | 40 | 1 | 2 | 2,36078 | 2,37027 | 2,67559 | 1,59384 | 1,8463 | 2,48145 | 1,63546 | 1,78047 | 1,75327 | 2,40637 | 1,26519 | 1,29038 | HexCer_40:1;2 |
| HexCer_42:1;2 | HexCer | MEM | 42 | 1 | 2 | 3,44491 | 3,37835 | 4,25874 | 0 | 2,8068 | 4,02599 | 3,0573 | 2,63652 | 2,28589 | 3,35315 | 0 | 0 | HexCer_42:1;2 |
| HexCer_42:2;2 | HexCer | MEM | 42 | 2 | 2 | 3,53467 | 3,50295 | 4,16283 | 2,49592 | 2,99233 | 4,14014 | 2,55372 | 2,63265 | 2,14976 | 3,43587 | 1,68828 | 1,92369 | HexCer_42:2;2 |
| LPA_18:0 | LPA | LYS | 18 | 0 | 0 | 0,295452 | 0 | 0 | 0 | 0 | 0 | 0 | 0 | 6,10159 | 0 | 0 | 0 | LPA_18:0 |
| LPC_16:0 | LPC | LYS | 16 | 0 | 0 | 4,52142 | 3,06641 | 4,03935 | 2,64283 | 3,48455 | 5,54746 | 6,05725 | 2,50021 | 2,0072 | 5,23956 | 3,45274 | 2,15928 | LPC_16:0 |
| LPC_18:0 | LPC | LYS | 18 | 0 | 0 | 1,75841 | 1,43002 | 1,7567 | 1,07563 | 1,47638 | 1,68393 | 1,40284 | 0,842563 | 1,1641 | 1,71565 | 1,30841 | 1,07604 | LPC_18:0 |

| | | | | | | | | | | | | | | | | | | |
|---|---|---|---|---|---|---|---|---|---|---|---|---|---|---|---|---|---|---|
| LPC_18:1 | LPC | LYS | 18 | 1 | 0 | 1,37261 | 1,01876 | 1,17299 | 0,646876 | 1,08887 | 1,64035 | 1,95659 | 0,674452 | 0,72287 | 1,22643 | 0,809225 | 0,485442 | LPC_18:1 |
| LPC_18:2 | LPC | LYS | 18 | 2 | 0 | 0,938115 | 0,698783 | 0,869121 | 0,591317 | 0,671928 | 1,23253 | 1,38166 | 0,626445 | 0,360073 | 0,970524 | 0,549064 | 0,339273 | LPC_18:2 |
| LPC_20:4 | LPC | LYS | 20 | 4 | 0 | 0,892053 | 0 | 0,711604 | 0,344047 | 0,422036 | 1,01387 | 1,40632 | 0,360432 | 0 | 0,707847 | 0 | 0 | LPC_20:4 |
| LPCO_16:0 | LPC O- | LYS | 16 | 0 | 0 | 1,69178 | 1,6456 | 1,72896 | 1,52839 | 1,94757 | 1,33932 | 1,09254 | 1,01115 | 3,13278 | 2,12808 | 1,81754 | 1,09077 | LPCO_16:0 |
| LPCO_18:0 | LPC O- | LYS | 18 | 0 | 0 | 0,299723 | 0 | 0 | 0 | 0 | 0 | 0 | 0 | 0,539423 | 0 | 0 | 0 | LPCO_18:0 |
| LPCO_18:1 | LPC O- | LYS | 18 | 1 | 0 | 0,627732 | 0,661494 | 0,709699 | 0,597928 | 0,823487 | 0,604096 | 0,546995 | 0,505028 | 1,08267 | 0,912482 | 0,732811 | 0,544283 | LPCO_18:1 |
| LPE_18:0 | LPE | LYS | 18 | 0 | 0 | 1,7993 | 1,12036 | 1,68091 | 1,0037 | 1,34087 | 1,69881 | 1,21715 | 0,874462 | 1,15359 | 1,66734 | 0,958907 | 0,847605 | LPE_18:0 |
| LPE_18:1 | LPE | LYS | 18 | 1 | 0 | 0,765774 | 0,497581 | 0,719105 | 0,487967 | 0,634627 | 0,856561 | 1,0447 | 0,51689 | 0,363466 | 0,783964 | 0,50661 | 0,370258 | LPE_18:1 |
| LPE_18:2 | LPE | LYS | 18 | 2 | 0 | 0,457979 | 0,378679 | 0,492821 | 0,399077 | 0,441819 | 0,485487 | 0,440862 | 0,435501 | 0,430503 | 0,436274 | 0,430567 | 0,375289 | LPE_18:2 |
| LPE_20:3 | LPE | LYS | 20 | 3 | 0 | 0,167225 | 0 | 0 | 0,140418 | 0 | 0,187309 | 0,134294 | 0,145895 | 0,119923 | 0,146508 | 0 | 0,108171 | LPE_20:3 |
| LPE_20:4 | LPE | LYS | 20 | 4 | 0 | 1,49452 | 1,18555 | 1,47966 | 0,997753 | 1,30589 | 1,49866 | 1,56288 | 1,01799 | 1,01407 | 1,25869 | 1,02743 | 0,857484 | LPE_20:4 |
| LPE_22:4 | LPE | LYS | 22 | 4 | 0 | 0,216561 | 0 | 0,178591 | 0,12529 | 0,186882 | 0,217601 | 0,289145 | 0 | 0 | 0,156889 | 0 | 0 | LPE_22:4 |
| LPE_22:6 | LPE | LYS | 22 | 6 | 0 | 0,307012 | 0 | 0,237287 | 0,169599 | 0,213764 | 0,446988 | 0,344769 | 0,217346 | 0 | 0,22097 | 0,135949 | 0,141844 | LPE_22:6 |
| LPEO_16:0 | LPE O- | LYS | 16 | 0 | 0 | 0,121947 | 0 | 0 | 0 | 0 | 0 | 0,089125 | 0 | 0,116681 | 0 | 0 | 0 | LPEO_16:0 |
| LPEO_16:1 | LPE O- | LYS | 16 | 1 | 0 | 0,828221 | 0,617796 | 0,801639 | 0,504707 | 0,693658 | 0,818997 | 0,660192 | 0,508072 | 0,943505 | 0,815455 | 0,591863 | 0,407943 | LPEO_16:1 |
| LPEO_18:1 | LPE O- | LYS | 18 | 1 | 0 | 0,578492 | 0,49529 | 0,629421 | 0,43612 | 0,501382 | 0,567518 | 0,3787 | 0,356606 | 0,719554 | 0,647093 | 0,415535 | 0,351086 | LPEO_18:1 |
| LPG_18:1 | LPG | LYS | 18 | 1 | 0 | 0,220385 | 0,16985 | 0,245769 | 0,122543 | 0,173545 | 0,254292 | 0,203991 | 0,124915 | 0 | 0,375121 | 0,131379 | 0,088634 | LPG_18:1 |
| LPG_18:2 | LPG | LYS | 18 | 2 | 0 | 0,186453 | 0,146593 | 0,14719 | 0,1195 | 0,155075 | 0,193291 | 0,174107 | 0,141195 | 0 | 0,263835 | 0,106347 | 0,091833 | LPG_18:2 |
| LPG_20:4 | LPG | LYS | 20 | 4 | 0 | 0,129243 | 0,079332 | 0,097564 | 0,08258 | 0,091718 | 0,126784 | 0,113808 | 0,056201 | 0 | 0,164491 | 0 | 0,046185 | LPG_20:4 |
| LPG_22:6 | LPG | LYS | 22 | 6 | 0 | 0,117559 | 0,083802 | 0,130078 | 0,074909 | 0,083932 | 0,098523 | 0,092301 | 0 | 0 | 0,187528 | 0 | 0,051817 | LPG_22:6 |
| LPI_18:0 | LPI | LYS | 18 | 0 | 0 | 0,74896 | 0,756442 | 0,99398 | 0,521705 | 0,691776 | 0,820694 | 0,501349 | 0,446108 | 2,08441 | 1,19256 | 0,703725 | 0,376908 | LPI_18:0 |
| LPI_20:4 | LPI | LYS | 20 | 4 | 0 | 0,462778 | 0,252492 | 0,403838 | 0,185141 | 0,23583 | 0,470774 | 0,500524 | 0,195664 | 0 | 0,414101 | 0,217245 | 0 | LPI_20:4 |
| LPS_18:0 | LPS | LYS | 18 | 0 | 0 | 0,86863 | 0,572725 | 0,824233 | 0,484884 | 0,702198 | 0,84721 | 0,831895 | 0,404149 | 1,07833 | 1,22563 | 0,633964 | 0,347068 | LPS_18:0 |
| LPS_18:2 | LPS | LYS | 18 | 2 | 0 | 0,217156 | 0 | 0,1706 | 0 | 0 | 0 | 0 | 0 | 0 | 0,180188 | 0 | 0 | LPS_18:2 |
| LPS_20:4 | LPS | LYS | 20 | 4 | 0 | 0 | 0,124358 | 0,184891 | 0 | 0,105366 | 0,230374 | 0,290663 | 0 | 0 | 0,21944 | 0 | 0 | LPS_20:4 |
| LPS_22:6 | LPS | LYS | 22 | 6 | 0 | 0 | 0 | 0 | 0 | 0 | 0 | 0,159967 | 0 | 0 | 0,12646 | 0 | 0 | LPS_22:6 |
| PA_16:0-16:0 | PA | MEM | 32 | 0 | 0 | 0,87952 | 0,885601 | 1,44007 | 0,78266 | 0,85191 | 1,37328 | 0,453958 | 0,614308 | 0 | 1,79429 | 1,38651 | 0,579794 | PA_16:0-16:0 |
| PA_16:0-18:1 | PA | MEM | 34 | 1 | 0 | 1,13437 | 0,7216 | 2,03652 | 0,927059 | 0,919276 | 1,11852 | 0,640597 | 0,654896 | 0 | 1,85646 | 1,36529 | 0,422077 | PA_16:0-18:1 |
| PA_16:0-18:2 | PA | MEM | 34 | 2 | 0 | 0 | 0,590264 | 0,953694 | 0,580524 | 0,667321 | 0,606197 | 0 | 0 | 0 | 1,25096 | 1,01524 | 0 | PA_16:0-18:2 |
| PA_16:1-18:0 | PA | MEM | 34 | 1 | 0 | 0 | 0,158668 | 0 | 0 | 0 | 0 | 0 | 0 | 0 | 0,326001 | 0 | 0 | PA_16:1-18:0 |
| PA_17:0-17:1 | PA | MEM | 34 | 1 | 0 | 0 | 0 | 0 | 0 | 0 | 0,29395 | 0 | 0 | 0 | 0 | 0,408579 | 0,143931 | PA_17:0-17:1 |
| PA_17:1-17:1 | PA | MEM | 34 | 2 | 0 | 0 | 0,014025 | 0,053664 | 0,012226 | 0 | 0,012676 | 0 | 0 | 0 | 0 | 0,041615 | 0 | PA_17:1-17:1 |
| PA_18:0-18:1 | PA | MEM | 36 | 1 | 0 | 0 | 0,246278 | 0 | 0 | 0 | 0 | 0 | 0 | 0 | 0,533975 | 0 | 0 | PA_18:0-18:1 |
| PA_18:0-18:2 | PA | MEM | 36 | 2 | 0 | 0 | 0,431162 | 0,90145 | 0,342319 | 0 | 0,451853 | 0 | 0 | 0 | 0,894942 | 0 | 0 | PA_18:0-18:2 |
| PA_18:0-20:4 | PA | MEM | 38 | 4 | 0 | 1,81589 | 1,61996 | 2,54166 | 1,25235 | 1,2867 | 1,99753 | 0,820451 | 1,11023 | 0 | 3,43698 | 1,73259 | 0,603531 | PA_18:0-20:4 |
| PA_18:1-18:1 | PA | MEM | 36 | 2 | 0 | 0 | 0,047867 | 0,180447 | 0,027569 | 0 | 0,054798 | 0 | 0 | 0 | 0,126981 | 0 | 0 | PA_18:1-18:1 |
| PA_18:1-18:2 | PA | MEM | 36 | 3 | 0 | 0 | 0,227506 | 0,637916 | 0,345924 | 0 | 0 | 0 | 0 | 0 | 0,630839 | 0 | 0 | PA_18:1-18:2 |
| PA_18:2-18:2 | PA | MEM | 36 | 4 | 0 | 0 | 0,406425 | 0,927299 | 0,371421 | 0,351512 | 0,770724 | 0,308043 | 0 | 0 | 1,06144 | 0,650528 | 0 | PA_18:2-18:2 |
| PC_14:0-16:0 | PC | MEM | 30 | 0 | 0 | 18,5445 | 14,4522 | 26,7813 | 9,13162 | 11,1844 | 26,9142 | 17,4576 | 11,5632 | 7,06925 | 16,6269 | 6,61961 | 7,3592 | PC_14:0-16:0 |
| PC_14:0-17:0 | PC | MEM | 31 | 0 | 0 | 0,595754 | 0,453159 | 0,700724 | 0,320837 | 0,331038 | 1,08894 | 0,775336 | 0,351163 | 0,152043 | 0,520583 | 0,279471 | 0,331662 | PC_14:0-17:0 |
| PC_14:0-18:0 | PC | MEM | 32 | 0 | 0 | 6,15121 | 5,36487 | 8,05074 | 3,89459 | 4,15256 | 8,22065 | 4,81376 | 3,79218 | 2,67782 | 6,10347 | 2,84809 | 3,45887 | PC_14:0-18:0 |
| PC_14:0-18:1 | PC | MEM | 32 | 1 | 0 | 9,50619 | 8,01954 | 13,4495 | 5,52919 | 6,5066 | 13,6339 | 10,2767 | 5,47556 | 4,3598 | 8,62853 | 4,37484 | 4,52304 | PC_14:0-18:1 |
| PC_14:0-18:2 | PC | MEM | 32 | 2 | 0 | 24,0854 | 31,7391 | 27,227 | 37,84 | 29,4123 | 38,6719 | 19,3668 | 21,3991 | 31,1332 | 35,7747 | 24,9863 | 24,5981 | PC_14:0-18:2 |
| PC_14:0-20:2 | PC | MEM | 34 | 2 | 0 | 0 | 0 | 0,680895 | 0 | 0 | 0,64481 | 0 | 0 | 0,304748 | 0 | 0 | 0 | PC_14:0-20:2 |
| PC_14:0-20:3 | PC | MEM | 34 | 3 | 0 | 0 | 0 | 1,06768 | 0 | 0 | 1,25224 | 0,85851 | 0,344028 | 0 | 0 | 0 | 0 | PC_14:0-20:3 |
| PC_14:0-20:4 | PC | MEM | 34 | 4 | 0 | 0 | 16,5182 | 16,6155 | 19,7729 | 0 | 22,2101 | 15,4332 | 14,2306 | 0 | 17,9961 | 11,9761 | 12,3221 | PC_14:0-20:4 |
| PC_15:0-15:0 | PC | MEM | 30 | 0 | 0 | 0,857025 | 0,896009 | 1,12744 | 0,583266 | 0,744354 | 1,03861 | 0,533861 | 0,703282 | 0,998313 | 0,904547 | 0,506433 | 0,409695 | PC_15:0-15:0 |
| PC_15:0-16:0 | PC | MEM | 31 | 0 | 0 | 13,2447 | 8,08584 | 14,8838 | 8,36465 | 9,48983 | 19,4134 | 15,154 | 7,73245 | 4,74813 | 9,8619 | 5,47229 | 6,26222 | PC_15:0-16:0 |
| PC_15:0-16:1 | PC | MEM | 31 | 1 | 0 | 0 | 0 | 6,24168 | 0 | 0 | 6,69504 | 6,09016 | 0 | 0 | 0 | 3,12914 | 0 | PC_15:0-16:1 |
| PC_15:0-17:0 | PC | MEM | 32 | 0 | 0 | 0,918991 | 0,989395 | 0,580894 | 0,70628 | 0,736477 | 0 | 0,501221 | 0,414536 | 0,424889 | 0 | 0,564539 | 0,868893 | PC_15:0-17:0 |
| PC_15:0-17:1 | PC | MEM | 32 | 1 | 0 | 0,488527 | 0,573147 | 0,831071 | 0,592859 | 0 | 0,947107 | 0,397677 | 0,383005 | 0,611158 | 0 | 0,289711 | 0,273187 | PC_15:0-17:1 |
| PC_15:0-18:0 | PC | MEM | 33 | 0 | 0 | 0,609737 | 0,333289 | 0,507446 | 0 | 0 | 0,635558 | 0,517117 | 0,367373 | 0 | 0 | 0 | 0,312044 | PC_15:0-18:0 |
| PC_15:0-18:1 | PC | MEM | 33 | 1 | 0 | 5,4427 | 3,15026 | 6,28497 | 10,7909 | 13,6904 | 7,04014 | 6,12117 | 2,82644 | 0 | 4,42351 | 2,46561 | 2,28217 | PC_15:0-18:1 |
| PC_16:0-16:0 | PC | MEM | 32 | 0 | 0 | 187,053 | 134,892 | 195,437 | 101,648 | 130,289 | 247,934 | 172,706 | 106,948 | 53,8133 | 155,763 | 71,0703 | 82,6165 | PC_16:0-16:0 |
| PC_16:0-16:1 | PC | MEM | 32 | 1 | 0 | 40,5822 | 30,5281 | 50,5924 | 21,479 | 25,8987 | 50,7723 | 42,1951 | 25,757 | 14,7458 | 41,6652 | 15,7948 | 17,1425 | PC_16:0-16:1 |

| | | | | | | | | | | | | | | | | | | |
|---|---|---|---|---|---|---|---|---|---|---|---|---|---|---|---|---|---|---|
| PC_16:0-16:2 | PC | MEM | 32 | 2 | 0 | 4,81105 | 5,66436 | 5,6063 | 6,67751 | 4,8438 | 9,75885 | 4,3299 | 2,98341 | 0 | 4,99991 | 0 | 4,23591 | PC_16:0-16:2 |
| PC_16:0-17:1 | PC | MEM | 33 | 1 | 0 | 13,3875 | 8,37478 | 14,5113 | 0 | 0 | 17,9304 | 14,1131 | 7,68596 | 0 | 11,4864 | 5,56524 | 5,9623 | PC_16:0-17:1 |
| PC_16:0-18:0 | PC | MEM | 34 | 0 | 0 | 38,3729 | 30,2175 | 41,7911 | 21,4179 | 30,0646 | 44,5748 | 31,5336 | 20,0837 | 15,5462 | 33,2486 | 17,5653 | 18,269 | PC_16:0-18:0 |
| PC_16:0-18:1 | PC | MEM | 34 | 1 | 0 | 243,565 | 176,834 | 278,625 | 136,069 | 173,375 | 306,624 | 240,224 | 119,087 | 85,5642 | 217,382 | 97,2817 | 100,892 | PC_16:0-18:1 |
| PC_16:0-18:2 | PC | MEM | 34 | 2 | 0 | 164,191 | 137,171 | 163,857 | 95,4315 | 113,868 | 195,604 | 144,054 | 98,9235 | 85,7754 | 143,963 | 71,8942 | 76,6521 | PC_16:0-18:2 |
| PC_16:0-18:3 | PC | MEM | 34 | 3 | 0 | 6,32907 | 5,24272 | 7,36421 | 3,63883 | 4,39683 | 11,3064 | 9,37767 | 4,98328 | 0 | 5,98338 | 2,59692 | 2,97963 | PC_16:0-18:3 |
| PC_16:0-19:1 | PC | MEM | 35 | 1 | 0 | 1,15423 | 0 | 1,08896 | 0,658829 | 0,754389 | 1,2361 | 0,649972 | 0,550145 | 0 | 0,874632 | 0 | 0,401601 | PC_16:0-19:1 |
| PC_16:0-20:1 | PC | MEM | 36 | 1 | 0 | 4,05521 | 3,06249 | 5,32462 | 2,43095 | 5,78007 | 5,94396 | 3,59587 | 2,59687 | 1,99715 | 5,14958 | 2,21759 | 1,90915 | PC_16:0-20:1 |
| PC_16:0-20:2 | PC | MEM | 36 | 2 | 0 | 12,7629 | 9,43209 | 11,6766 | 6,39689 | 8,48327 | 15,1863 | 8,21149 | 6,99028 | 5,1538 | 9,82235 | 4,64069 | 5,36576 | PC_16:0-20:2 |
| PC_16:0-20:3 | PC | MEM | 36 | 3 | 0 | 16,3408 | 13,2955 | 18,9867 | 8,10703 | 11,657 | 21,0644 | 15,8103 | 9,30352 | 6,45756 | 14,7935 | 6,63539 | 7,0405 | PC_16:0-20:3 |
| PC_16:0-20:4 | PC | MEM | 36 | 4 | 0 | 168,874 | 120,964 | 162,022 | 77,457 | 107,004 | 192,957 | 146,281 | 78,6808 | 36,8109 | 132,977 | 57,4672 | 63,2975 | PC_16:0-20:4 |
| PC_16:0-20:5 | PC | MEM | 36 | 5 | 0 | 0 | 0 | 8,05681 | 6,64039 | 0 | 10,7799 | 6,55332 | 0 | 0 | 0 | 2,56404 | 3,92041 | PC_16:0-20:5 |
| PC_16:0-22:2 | PC | MEM | 38 | 2 | 0 | 0 | 0 | 0,778455 | 0 | 0 | 0,758834 | 0,71955 | 0 | 0 | 0 | 0 | 0 | PC_16:0-22:2 |
| PC_16:0-22:3 | PC | MEM | 38 | 3 | 0 | 1,13311 | 0,845914 | 1,14521 | 0,497205 | 0 | 1,26052 | 0,822305 | 0 | 0 | 0,963119 | 0,430987 | 0,410995 | PC_16:0-22:3 |
| PC_16:0-22:4 | PC | MEM | 38 | 4 | 0 | 17,764 | 11,1355 | 15,7473 | 7,28802 | 10,7589 | 21,2976 | 12,9644 | 6,3809 | 3,66041 | 13,1519 | 5,70238 | 6,49001 | PC_16:0-22:4 |
| PC_16:0-22:5 | PC | MEM | 38 | 5 | 0 | 6,97528 | 5,79437 | 9,92596 | 4,26714 | 5,96607 | 10,1992 | 7,5821 | 4,56291 | 2,10967 | 6,98239 | 3,08686 | 3,70486 | PC_16:0-22:5 |
| PC_16:0-22:6 | PC | MEM | 38 | 6 | 0 | 14,0238 | 9,60177 | 0 | 6,49106 | 8,71745 | 19,1921 | 12,434 | 6,74138 | 0 | 10,933 | 5,07799 | 5,45945 | PC_16:0-22:6 |
| PC_16:0-24:4 | PC | MEM | 40 | 4 | 0 | 1,64417 | 1,19482 | 2,09918 | 0,962586 | 1,18236 | 2,45125 | 2,34373 | 1,48853 | 0 | 1,66719 | 0,879402 | 0,962291 | PC_16:0-24:4 |
| PC_16:0-24:5 | PC | MEM | 40 | 5 | 0 | 1,31344 | 0 | 0,888115 | 0 | 0 | 1,43523 | 1,60653 | 0 | 0 | 0 | 0 | 0 | PC_16:0-24:5 |
| PC_16:1-16:1 | PC | MEM | 32 | 2 | 0 | 3,36428 | 3,22063 | 5,96282 | 5,16815 | 3,35409 | 6,17356 | 4,94326 | 3,06137 | 2,39845 | 6,54689 | 2,47489 | 3,51401 | PC_16:1-16:1 |
| PC_16:1-17:1 | PC | MEM | 33 | 2 | 0 | 0 | 0 | 11,6649 | 6,86634 | 0 | 0 | 0,661529 | 0 | 0 | 0 | 0 | 5,85698 | PC_16:1-17:1 |
| PC_16:1-18:0 | PC | MEM | 34 | 1 | 0 | 3,28174 | 3,04788 | 5,2014 | 2,39534 | 3,04426 | 4,46414 | 2,76574 | 2,20909 | 2,0348 | 4,00147 | 1,61432 | 1,8798 | PC_16:1-18:0 |
| PC_16:1-18:1 | PC | MEM | 34 | 2 | 0 | 11,4382 | 9,95655 | 19,2614 | 7,81244 | 8,43785 | 16,4492 | 12,2948 | 6,79046 | 7,57892 | 13,3185 | 5,55655 | 5,78349 | PC_16:1-18:1 |
| PC_16:1-18:2 | PC | MEM | 34 | 3 | 0 | 7,68979 | 7,84205 | 10,361 | 6,46602 | 6,48855 | 10,7015 | 6,70684 | 6,76832 | 7,31625 | 8,43745 | 5,04639 | 4,77479 | PC_16:1-18:2 |
| PC_16:1-20:2 | PC | MEM | 36 | 3 | 0 | 0 | 0 | 0,502554 | 0 | 0 | 0,476607 | 0 | 0 | 0,430999 | 0 | 0 | 0 | PC_16:1-20:2 |
| PC_16:1-20:3 | PC | MEM | 36 | 4 | 0 | 0 | 0 | 1,77377 | 0,505401 | 0 | 2,00029 | 1,96147 | 1,28372 | 0 | 0,866232 | 0 | 0,523976 | PC_16:1-20:3 |
| PC_16:1-20:4 | PC | MEM | 36 | 5 | 0 | 23,6219 | 21,9903 | 16,03 | 16,0818 | 19,171 | 25,4843 | 12,6785 | 11,7196 | 10,1197 | 0 | 8,45663 | 10,1723 | PC_16:1-20:4 |
| PC_17:0-18:1 | PC | MEM | 35 | 1 | 0 | 3,70198 | 3,29073 | 3,18855 | 2,20228 | 2,87782 | 4,49469 | 3,17258 | 1,79389 | 2,36271 | 3,27252 | 1,89799 | 1,52647 | PC_17:0-18:1 |
| PC_17:0-18:2 | PC | MEM | 35 | 2 | 0 | 8,82591 | 7,70372 | 6,00985 | 6,47934 | 7,41182 | 6,85768 | 6,96743 | 4,89008 | 6,73839 | 4,50657 | 3,0247 | 2,98302 | PC_17:0-18:2 |
| PC_17:0-20:4 | PC | MEM | 37 | 4 | 0 | 11,3957 | 7,96699 | 11,0961 | 5,51004 | 7,70943 | 9,97709 | 7,90443 | 5,63602 | 0 | 9,00182 | 0 | 5,17891 | PC_17:0-20:4 |
| PC_17:0-22:4 | PC | MEM | 39 | 4 | 0 | 2,68216 | 1,50336 | 2,09209 | 1,73522 | 1,85797 | 2,42283 | 2,21699 | 1,68506 | 0 | 2,17249 | 1,31881 | 1,91744 | PC_17:0-22:4 |
| PC_17:1-18:0 | PC | MEM | 35 | 1 | 0 | 0 | 0 | 0 | 0 | 0 | 0 | 0 | 0 | 0 | 0 | 0,262957 | 0,185447 | PC_17:1-18:0 |
| PC_17:1-18:1 | PC | MEM | 35 | 2 | 0 | 0 | 0 | 4,24195 | 0 | 0 | 3,59524 | 0 | 0 | 0 | 2,95784 | 1,48209 | 1,72477 | PC_17:1-18:1 |
| PC_17:1-22:4 | PC | MEM | 39 | 5 | 0 | 0 | 1,11161 | 1,42701 | 0 | 0 | 0 | 0 | 1,2255 | 0 | 0 | 0,995392 | 1,1958 | PC_17:1-22:4 |
| PC_17:1-22:5 | PC | MEM | 39 | 6 | 0 | 0 | 0 | 0 | 0 | 1,50033 | 2,64631 | 0 | 0 | 0 | 1,76799 | 0 | 0 | PC_17:1-22:5 |
| PC_18:0-18:1 | PC | MEM | 36 | 1 | 0 | 17,9843 | 15,6513 | 21,7352 | 11,404 | 12,4055 | 17,5134 | 12,9886 | 7,9336 | 11,945 | 15,6537 | 8,24949 | 9,08243 | PC_18:0-18:1 |
| PC_18:0-18:2 | PC | MEM | 36 | 2 | 0 | 59,6216 | 59,4811 | 64,0306 | 37,3789 | 45,8313 | 63,3301 | 32,5847 | 33,2665 | 45,1483 | 56,895 | 29,9394 | 30,5763 | PC_18:0-18:2 |
| PC_18:0-18:3 | PC | MEM | 36 | 3 | 0 | 0,46703 | 0 | 0,633996 | 0,315022 | 0 | 0,926173 | 0 | 0 | 0 | 0 | 0 | 0 | PC_18:0-18:3 |
| PC_18:0-20:1 | PC | MEM | 38 | 1 | 0 | 0 | 0 | 0 | 0 | 0 | 0 | 0 | 0 | 0,718325 | 0,889414 | 0 | 0 | PC_18:0-20:1 |
| PC_18:0-20:2 | PC | MEM | 38 | 2 | 0 | 1,86088 | 1,55715 | 1,6755 | 0,911265 | 1,44718 | 1,33549 | 0,99776 | 1,19953 | 1,37546 | 1,49388 | 0,853977 | 0,737832 | PC_18:0-20:2 |
| PC_18:0-20:3 | PC | MEM | 38 | 3 | 0 | 5,42051 | 5,00721 | 7,13853 | 3,2128 | 4,45344 | 5,71998 | 4,0767 | 3,08142 | 3,66618 | 4,88586 | 2,59873 | 2,42908 | PC_18:0-20:3 |
| PC_18:0-20:4 | PC | MEM | 38 | 4 | 0 | 84,7968 | 68,528 | 80,8591 | 36,9382 | 54,9975 | 83,8103 | 47,528 | 34,3633 | 24,581 | 69,3973 | 29,6499 | 35,5874 | PC_18:0-20:4 |
| PC_18:0-22:4 | PC | MEM | 40 | 4 | 0 | 4,08691 | 3,54765 | 2,97948 | 1,52049 | 2,79321 | 2,92247 | 1,73888 | 0,981811 | 1,32581 | 2,82678 | 1,16032 | 1,68091 | PC_18:0-22:4 |
| PC_18:0-22:5 | PC | MEM | 40 | 5 | 0 | 2,93531 | 2,75516 | 2,50301 | 1,36303 | 0 | 2,26184 | 1,91711 | 1,34238 | 1,76944 | 1,71559 | 1,16648 | 1,34342 | PC_18:0-22:5 |
| PC_18:1-18:1 | PC | MEM | 36 | 2 | 0 | 29,7573 | 28,2241 | 44,4934 | 20,89 | 26,9531 | 35,6467 | 24,0965 | 15,7407 | 21,621 | 33,4385 | 15,5937 | 16,0023 | PC_18:1-18:1 |
| PC_18:1-18:2 | PC | MEM | 36 | 3 | 0 | 43,1529 | 42,7988 | 58,5284 | 32,191 | 35,4174 | 52,027 | 33,4243 | 27,37 | 34,4892 | 44,4142 | 24,3283 | 23,7718 | PC_18:1-18:2 |
| PC_18:1-18:3 | PC | MEM | 36 | 4 | 0 | 1,50586 | 1,36243 | 3,23216 | 0 | 1,06683 | 3,78641 | 3,12817 | 1,32749 | 0,51428 | 1,16901 | 0,664757 | 0,59163 | PC_18:1-18:3 |
| PC_18:1-20:1 | PC | MEM | 38 | 2 | 0 | 1,97086 | 1,62107 | 2,35333 | 1,28968 | 1,74947 | 1,54724 | 0,85483 | 1,0082 | 1,66841 | 1,94872 | 0,910445 | 1,06102 | PC_18:1-20:1 |
| PC_18:1-20:2 | PC | MEM | 38 | 3 | 0 | 2,41834 | 3,10018 | 4,17569 | 2,18891 | 2,73999 | 3,22333 | 2,2384 | 2,04868 | 3,26903 | 3,01022 | 1,86706 | 1,79506 | PC_18:1-20:2 |
| PC_18:1-20:3 | PC | MEM | 38 | 4 | 0 | 7,72119 | 6,94819 | 11,1082 | 4,82826 | 6,58199 | 9,97406 | 7,68349 | 4,87388 | 4,81296 | 7,69814 | 3,61455 | 3,8709 | PC_18:1-20:3 |
| PC_18:1-20:4 | PC | MEM | 38 | 5 | 0 | 42,5723 | 31,8703 | 46,9661 | 21,4242 | 27,6516 | 47,1073 | 32,5834 | 19,2508 | 12,4387 | 37,4773 | 14,2246 | 16,4764 | PC_18:1-20:4 |
| PC_18:1-22:4 | PC | MEM | 40 | 5 | 0 | 3,16265 | 1,9813 | 3,43926 | 1,46478 | 4,36557 | 3,55745 | 1,8428 | 1,90621 | 2,19394 | 2,62264 | 1,43671 | 1,57236 | PC_18:1-22:4 |
| PC_18:1-22:5 | PC | MEM | 40 | 6 | 0 | 0 | 0 | 2,50626 | 0 | 0 | 3,48094 | 0 | 2,10738 | 0 | 0 | 0 | 2,21922 | PC_18:1-22:5 |
| PC_18:2-18:2 | PC | MEM | 36 | 4 | 0 | 23,3151 | 23,5198 | 25,3346 | 17,1654 | 18,158 | 27,104 | 17,1997 | 18,6319 | 15,1375 | 21,8185 | 11,9511 | 13,2423 | PC_18:2-18:2 |
| PC_18:2-18:3 | PC | MEM | 36 | 5 | 0 | 0 | 0 | 0 | 0 | 0 | 0 | 2,66287 | 3,66704 | 0 | 0 | 1,23784 | 0 | PC_18:2-18:3 |

| | | | | | | | | | | | | | | | | | | |
|---|---|---|---|---|---|---|---|---|---|---|---|---|---|---|---|---|---|---|
| PC_18:2-20:1 | PC | MEM | 38 | 3 | 0 | 1,6026 | 1,3487 | 2,94746 | 1,34854 | 1,37999 | 2,13786 | 1,02711 | 1,1821 | 2,04876 | 1,48756 | 1,01993 | 0,858353 | PC_18:2-20:1 |
| PC_18:2-20:2 | PC | MEM | 38 | 4 | 0 | 8,32837 | 8,7131 | 9,53515 | 5,38303 | 6,85653 | 9,90175 | 4,16904 | 5,9039 | 7,58387 | 7,76146 | 4,12463 | 4,44423 | PC_18:2-20:2 |
| PC_18:2-20:3 | PC | MEM | 38 | 5 | 0 | 4,67188 | 4,80225 | 5,80303 | 2,68732 | 2,99934 | 5,30267 | 4,36207 | 3,80143 | 3,40182 | 3,83109 | 2,81395 | 2,3741 | PC_18:2-20:3 |
| PC_18:2-20:4 | PC | MEM | 38 | 6 | 0 | 17,5227 | 13,6452 | 31,1279 | 8,71474 | 12,0644 | 22,7379 | 14,0093 | 10,3563 | 9,03954 | 14,8641 | 6,67934 | 7,96855 | PC_18:2-20:4 |
| PC_18:2-22:4 | PC | MEM | 40 | 6 | 0 | 0 | 4,05568 | 2,21792 | 0 | 0 | 3,95645 | 2,96374 | 0 | 0 | 0 | 0 | 0 | PC_18:2-22:4 |
| PC_20:1-20:4 | PC | MEM | 40 | 5 | 0 | 1,3703 | 1,45939 | 2,55162 | 1,41631 | 1,6281 | 1,6907 | 1,23881 | 1,12214 | 0 | 2,03488 | 0,919444 | 0,583245 | PC_20:1-20:4 |
| PC_20:2-20:2 | PC | MEM | 40 | 4 | 0 | 0 | 0 | 0,220658 | 0,224831 | 0 | 0 | 0 | 0,192976 | 0,436895 | 0 | 0,187839 | 0 | PC_20:2-20:2 |
| PC_20:2-20:4 | PC | MEM | 40 | 6 | 0 | 12,5006 | 6,79533 | 4,40515 | 6,20818 | 9,39512 | 7,39774 | 5,06698 | 4,18422 | 5,94728 | 11,0066 | 5,12094 | 4,04044 | PC_20:2-20:4 |
| PC_20:3-20:4 | PC | MEM | 40 | 7 | 0 | 6,3595 | 0 | 7,18049 | 2,95514 | 0 | 7,5734 | 5,08976 | 3,27355 | 0 | 0 | 2,08701 | 2,37483 | PC_20:3-20:4 |
| PC_20:4-20:4 | PC | MEM | 40 | 8 | 0 | 2,99318 | 2,1359 | 3,34534 | 0 | 0 | 3,02617 | 2,03232 | 1,48267 | 0 | 2,20983 | 0 | 0 | PC_20:4-20:4 |
| PCO_16:0-16:0 | PC O- | MEM | 32 | 0 | 0 | 64,9206 | 83,9239 | 89,4435 | 72,5374 | 76,4488 | 61,4637 | 30,7713 | 65,737 | 127,815 | 91,4066 | 64,3251 | 61,988 | PCO_16:0-16:0 |
| PCO_16:0-16:1 | PC O- | MEM | 32 | 1 | 0 | 7,3436 | 9,60585 | 11,0568 | 8,33643 | 7,54756 | 5,92309 | 3,48574 | 6,22114 | 12,8222 | 10,0347 | 6,25568 | 6,75987 | PCO_16:0-16:1 |
| PCO_16:0-16:2 | PC O- | MEM | 32 | 2 | 0 | 0 | 0 | 0 | 0,469528 | 0 | 0 | 0 | 0,215291 | 0 | 0 | 0 | 0,313179 | PCO_16:0-16:2 |
| PCO_16:0-17:0 | PC O- | MEM | 33 | 0 | 0 | 0,002087 | 0,004827 | 0,002085 | 0,00361 | 0,005325 | 0 | 0 | 0,003454 | 0,007263 | 0 | 0,003855 | 0,007548 | PCO_16:0-17:0 |
| PCO_16:0-17:1 | PC O- | MEM | 33 | 1 | 0 | 0,036327 | 0,06492 | 0,022114 | 0,097443 | 0 | 0,03682 | 0,011486 | 0,029427 | 0,221308 | 0,035828 | 0,037908 | 0,036972 | PCO_16:0-17:1 |
| PCO_16:0-18:0 | PC O- | MEM | 34 | 0 | 0 | 0,369473 | 0,30585 | 0,392223 | 0,189064 | 0,290475 | 0,2541 | 0,145393 | 0,180371 | 0,430069 | 0,436164 | 0,161395 | 0,169271 | PCO_16:0-18:0 |
| PCO_16:0-18:1 | PC O- | MEM | 34 | 1 | 0 | 22,8739 | 23,9953 | 30,1637 | 20,0243 | 22,9685 | 19,8757 | 12,4178 | 14,7814 | 32,4321 | 26,8343 | 16,1925 | 15,6922 | PCO_16:0-18:1 |
| PCO_16:0-18:2 | PC O- | MEM | 34 | 2 | 0 | 39,7749 | 57,4112 | 56,9911 | 46,3329 | 49,9755 | 38,038 | 18,254 | 40,1612 | 85,4784 | 55,6485 | 40,3553 | 37,4105 | PCO_16:0-18:2 |
| PCO_16:0-18:3 | PC O- | MEM | 34 | 3 | 0 | 0,129873 | 0,295072 | 0,313153 | 0,1964 | 0,257643 | 0,349499 | 0,128162 | 0,200406 | 0,157354 | 0,331793 | 0 | 0,110091 | PCO_16:0-18:3 |
| PCO_16:0-20:1 | PC O- | MEM | 36 | 1 | 0 | 0 | 0 | 0,120165 | 0,073629 | 0 | 0,039812 | 0,034858 | 0,0522 | 0,139077 | 0,120534 | 0,046247 | 0 | PCO_16:0-20:1 |
| PCO_16:0-20:2 | PC O- | MEM | 36 | 2 | 0 | 1,32122 | 0,979099 | 1,20633 | 0,986526 | 0,978729 | 1,15299 | 0,361488 | 0,76843 | 1,75716 | 1,45388 | 0,760682 | 0,712959 | PCO_16:0-20:2 |
| PCO_16:0-20:3 | PC O- | MEM | 36 | 3 | 0 | 3,58549 | 4,36506 | 5,85038 | 3,52186 | 3,88375 | 3,85428 | 2,50505 | 2,86017 | 5,64193 | 4,13251 | 2,52819 | 2,56475 | PCO_16:0-20:3 |
| PCO_16:0-20:4 | PC O- | MEM | 36 | 4 | 0 | 80,2258 | 72,9217 | 77,6893 | 51,1635 | 67,8201 | 65,4753 | 35,0127 | 42,2655 | 68,8597 | 77,3355 | 40,9747 | 37,6705 | PCO_16:0-20:4 |
| PCO_16:0-22:4 | PC O- | MEM | 38 | 4 | 0 | 5,46218 | 4,59813 | 4,6428 | 3,85896 | 4,67247 | 4,72786 | 2,3671 | 2,75024 | 3,14426 | 4,91052 | 2,10778 | 2,81085 | PCO_16:0-22:4 |
| PCO_16:0-22:5 | PC O- | MEM | 38 | 5 | 0 | 0,984214 | 0,966353 | 2,32303 | 0,432006 | 1,04012 | 3,12936 | 0,89238 | 0,958896 | 0 | 1,48504 | 0,553257 | 0,439544 | PCO_16:0-22:5 |
| PCO_16:0-22:6 | PC O- | MEM | 38 | 6 | 0 | 2,12876 | 0 | 1,25925 | 0,797297 | 1,01752 | 1,06735 | 0 | 0 | 0 | 0 | 0 | 0 | PCO_16:0-22:6 |
| PCO_16:0-24:4 | PC O- | MEM | 40 | 4 | 0 | 2,34302 | 1,07373 | 1,85382 | 1,59303 | 1,72682 | 1,82956 | 0,911662 | 1,6808 | 1,94688 | 1,3147 | 1,57776 | 0,971067 | PCO_16:0-24:4 |
| PCO_16:1-16:0 | PC O- | MEM | 32 | 1 | 0 | 3,38136 | 3,7798 | 4,98887 | 3,16824 | 3,46601 | 3,60792 | 2,79101 | 3,14596 | 4,49097 | 4,21986 | 2,38972 | 2,78066 | PCO_16:1-16:0 |
| PCO_16:1-16:1 | PC O- | MEM | 32 | 2 | 0 | 0 | 0 | 0,341219 | 0 | 0 | 0 | 0 | 0,326667 | 0,581697 | 0,582505 | 0 | 0,419069 | PCO_16:1-16:1 |
| PCO_16:1-17:0 | PC O- | MEM | 33 | 1 | 0 | 0,197797 | 0,248497 | 0,216411 | 0,308733 | 0,247107 | 0,183533 | 0,104689 | 0,194133 | 0,386165 | 0,182784 | 0,235167 | 0,250999 | PCO_16:1-17:0 |
| PCO_16:1-18:0 | PC O- | MEM | 34 | 1 | 0 | 0,896516 | 1,20996 | 0,386272 | 0,514744 | 0,551272 | 0,316519 | 0,344692 | 0,662638 | 0,313714 | 0,739294 | 0,318636 | 0,378551 | PCO_16:1-18:0 |
| PCO_16:1-18:1 | PC O- | MEM | 34 | 2 | 0 | 1,10291 | 0,991361 | 1,67964 | 0,920489 | 1,01967 | 1,15041 | 0,751856 | 0,944581 | 1,44106 | 1,33708 | 0,578742 | 0,786716 | PCO_16:1-18:1 |
| PCO_16:1-18:2 | PC O- | MEM | 34 | 3 | 0 | 1,15622 | 1,08496 | 1,90042 | 1,60628 | 1,24136 | 1,20255 | 0,745653 | 1,02815 | 2,76483 | 1,52365 | 1,4796 | 1,14645 | PCO_16:1-18:2 |
| PCO_16:1-20:4 | PC O- | MEM | 36 | 5 | 0 | 3,74067 | 2,27383 | 4,38076 | 3,32923 | 3,27449 | 3,91202 | 2,74051 | 2,32104 | 3,56847 | 3,89805 | 2,01785 | 2,53326 | PCO_16:1-20:4 |
| PCO_16:2-16:0 | PC O- | MEM | 32 | 2 | 0 | 2,2946 | 2,75373 | 1,65782 | 2,00281 | 1,85453 | 2,31242 | 1,00464 | 1,26916 | 1,24359 | 1,91917 | 1,32656 | 0,980039 | PCO_16:2-16:0 |
| PCO_16:2-18:0 | PC O- | MEM | 34 | 2 | 0 | 1,73792 | 1,42528 | 0,678226 | 0,553297 | 0,750023 | 0,817136 | 0,197346 | 1,16169 | 0,337123 | 0,294672 | 0,476712 | 0,204366 | PCO_16:2-18:0 |
| PCO_16:2-18:1 | PC O- | MEM | 34 | 3 | 0 | 0,56766 | 0,793724 | 0,647822 | 0,322362 | 0,338661 | 0,333742 | 0,173351 | 0,472205 | 0,265277 | 0,447397 | 0,231244 | 0,440209 | PCO_16:2-18:1 |
| PCO_16:2-18:2 | PC O- | MEM | 34 | 4 | 0 | 0 | 0 | 0 | 0 | 0 | 0 | 0 | 0 | 1,0889 | 0 | 0 | 0,612162 | PCO_16:2-18:2 |
| PCO_16:2-20:4 | PC O- | MEM | 36 | 6 | 0 | 0 | 0 | 1,98453 | 0 | 0 | 1,37015 | 0,936602 | 0,811327 | 1,88641 | 0,913361 | 0 | 0 | PCO_16:2-20:4 |
| PCO_17:0-15:0 | PC O- | MEM | 32 | 0 | 0 | 2,21415 | 1,96039 | 2,38784 | 1,72243 | 1,72366 | 3,18895 | 1,89717 | 1,42877 | 1,32005 | 2,34275 | 1,15842 | 1,17298 | PCO_17:0-15:0 |
| PCO_17:0-16:0 | PC O- | MEM | 33 | 0 | 0 | 1,60097 | 2,07999 | 1,77283 | 1,76011 | 1,73191 | 0 | 0 | 1,45151 | 2,71845 | 2,25968 | 1,35481 | 1,44538 | PCO_17:0-16:0 |
| PCO_17:0-16:1 | PC O- | MEM | 33 | 1 | 0 | 3,06629 | 3,48669 | 3,7724 | 2,58031 | 2,97874 | 3,13935 | 1,84452 | 2,20482 | 3,4292 | 3,64439 | 1,94226 | 2,06625 | PCO_17:0-16:1 |
| PCO_17:0-16:2 | PC O- | MEM | 33 | 2 | 0 | 0,461098 | 0,555218 | 0,921876 | 0,347751 | 0,577428 | 0,766248 | 0,276183 | 0,239354 | 0 | 0,410196 | 0 | 0,547196 | PCO_17:0-16:2 |
| PCO_17:0-17:0 | PC O- | MEM | 34 | 0 | 0 | 2,35497 | 2,30368 | 2,15096 | 2,04359 | 2,19066 | 1,70036 | 1,12445 | 1,74482 | 3,28169 | 2,41192 | 2,03407 | 2,21257 | PCO_17:0-17:0 |
| PCO_17:0-17:1 | PC O- | MEM | 34 | 1 | 0 | 0,718018 | 0,798506 | 0,996919 | 0,452383 | 0,334352 | 1,49228 | 0,679859 | 0,729149 | 0,3173 | 1,21307 | 0,51022 | 0,418581 | PCO_17:0-17:1 |
| PCO_17:0-18:1 | PC O- | MEM | 35 | 1 | 0 | 0 | 4,35264 | 0 | 1,4965 | 0 | 0 | 0 | 1,28963 | 0,962245 | 2,86368 | 0 | 0,637111 | PCO_17:0-18:1 |
| PCO_17:0-18:2 | PC O- | MEM | 35 | 2 | 0 | 2,20396 | 2,52398 | 2,28744 | 2,1655 | 2,29698 | 1,26996 | 0 | 1,55615 | 3,20825 | 2,51598 | 1,74277 | 1,53094 | PCO_17:0-18:2 |
| PCO_17:0-18:3 | PC O- | MEM | 35 | 3 | 0 | 0,56885 | 0,594408 | 0,840172 | 0,49226 | 0,550903 | 0,742508 | 0,472462 | 0,438437 | 0 | 0,646045 | 0,334926 | 0,404528 | PCO_17:0-18:3 |
| PCO_17:0-19:1 | PC O- | MEM | 36 | 1 | 0 | 0,120611 | 0 | 0,088124 | 0,051838 | 0,076654 | 0,157011 | 0,02829 | 0,070824 | 0 | 0,115763 | 0 | 0,046466 | PCO_17:0-19:1 |
| PCO_17:0-20:1 | PC O- | MEM | 37 | 1 | 0 | 1,94847 | 3,85984 | 0 | 0 | 0 | 1,85879 | 0 | 0,843909 | 0 | 1,6732 | 0 | 0 | PCO_17:0-20:1 |
| PCO_17:0-20:2 | PC O- | MEM | 37 | 2 | 0 | 1,99352 | 4,96041 | 0 | 1,00162 | 0 | 1,9318 | 0 | 1,33819 | 0 | 2,80751 | 0 | 0 | PCO_17:0-20:2 |
| PCO_17:0-20:3 | PC O- | MEM | 37 | 3 | 0 | 0 | 1,87853 | 0 | 0 | 0 | 0 | 0 | 0 | 0 | 1,3653 | 0 | 0 | PCO_17:0-20:3 |
| PCO_17:0-20:4 | PC O- | MEM | 37 | 4 | 0 | 3,73586 | 3,07852 | 3,4563 | 2,70353 | 3,39873 | 2,55644 | 0,875925 | 2,17655 | 3,50125 | 3,62246 | 2,2303 | 1,97054 | PCO_17:0-20:4 |
| PCO_17:0-20:5 | PC O- | MEM | 37 | 5 | 0 | 0 | 0 | 0,393976 | 0,213522 | 0 | 0,333278 | 0,283376 | 0 | 0 | 0 | 0,16067 | 0,209488 | PCO_17:0-20:5 |
| PCO_17:0-22:4 | PC O- | MEM | 39 | 4 | 0 | 0 | 0 | 1,44566 | 0 | 0,855494 | 1,38397 | 0 | 0 | 0 | 0 | 0 | 0 | PCO_17:0-22:4 |

| | | | | | | | | | | | | | | | | | | |
|---|---|---|---|---|---|---|---|---|---|---|---|---|---|---|---|---|---|---|
| PCO_17:0-22:5 | PC O- | MEM | 39 | 5 | 0 | 0 | 0 | 0 | 0 | 0,762494 | 0 | 0 | 0 | 0 | 0,953383 | 0 | 0 | PCO_17:0-22:5 |
| PCO_17:1-15:0 | PC O- | MEM | 32 | 1 | 0 | 0 | 0 | 0,199166 | 0 | 0 | 0,118974 | 0,096574 | 0 | 0 | 0 | 0,102464 | 0 | PCO_17:1-15:0 |
| PCO_17:1-16:0 | PC O- | MEM | 33 | 1 | 0 | 3,31031 | 3,90235 | 3,88656 | 3,04652 | 3,28029 | 3,3426 | 1,8595 | 2,48093 | 4,23823 | 4,04453 | 2,37825 | 2,44389 | PCO_17:1-16:0 |
| PCO_17:1-16:1 | PC O- | MEM | 33 | 2 | 0 | 1,66166 | 1,86699 | 2,43754 | 1,96469 | 1,72114 | 1,52189 | 1,12809 | 1,78794 | 3,14233 | 2,81965 | 1,61159 | 1,51078 | PCO_17:1-16:1 |
| PCO_17:1-17:1 | PC O- | MEM | 34 | 2 | 0 | 0 | 0 | 0,20403 | 0,215872 | 0 | 0 | 0,098473 | 0 | 0 | 0 | 0 | 0,196771 | PCO_17:1-17:1 |
| PCO_17:1-18:0 | PC O- | MEM | 35 | 1 | 0 | 0 | 0,053715 | 0 | 0,016454 | 0 | 0 | 0 | 0,016093 | 0,014682 | 0,033806 | 0 | 0,00801 | PCO_17:1-18:0 |
| PCO_17:1-18:1 | PC O- | MEM | 35 | 2 | 0 | 0,127509 | 0,163098 | 0,239458 | 0,15653 | 0,1397 | 0,097522 | 0 | 0,096909 | 0,249806 | 0,207632 | 0,117027 | 0,099196 | PCO_17:1-18:1 |
| PCO_17:1-18:2 | PC O- | MEM | 35 | 3 | 0 | 1,02944 | 1,6304 | 1,70901 | 1,37044 | 1,24049 | 1,00309 | 0,500946 | 1,07504 | 2,65398 | 1,35738 | 1,17834 | 1,14461 | PCO_17:1-18:2 |
| PCO_17:1-20:3 | PC O- | MEM | 37 | 4 | 0 | 0,023808 | 0,020444 | 0,059623 | 0,021213 | 0 | 0,035513 | 0,017502 | 0,057771 | 0,067582 | 0,028464 | 0,030173 | 0,016388 | PCO_17:1-20:3 |
| PCO_17:1-20:4 | PC O- | MEM | 37 | 5 | 0 | 4,43751 | 3,70738 | 3,56348 | 2,36046 | 3,20011 | 3,56987 | 2,04285 | 2,36041 | 3,1125 | 4,134 | 1,92792 | 2,17928 | PCO_17:1-20:4 |
| PCO_17:2-16:0 | PC O- | MEM | 33 | 2 | 0 | 1,91515 | 2,72841 | 1,36992 | 2,19073 | 1,90816 | 1,63946 | 0,711939 | 1,50306 | 3,64356 | 1,74318 | 2,1329 | 1,27396 | PCO_17:2-16:0 |
| PCO_17:2-17:0 | PC O- | MEM | 34 | 2 | 0 | 0,152031 | 0 | 0 | 0,17545 | 0 | 0 | 0,18506 | 0 | 0 | 0 | 0 | 0,09971 | PCO_17:2-17:0 |
| PCO_17:2-18:0 | PC O- | MEM | 35 | 2 | 0 | 0 | 0,009216 | 0,006949 | 0,029961 | 0,011186 | 0,003584 | 0 | 0,007023 | 0,039916 | 0,008728 | 0,018835 | 0,008648 | PCO_17:2-18:0 |
| PCO_17:2-18:1 | PC O- | MEM | 35 | 3 | 0 | 0 | 0 | 0,059942 | 0,095798 | 0,054862 | 0,040201 | 0,066544 | 0,029926 | 0,114814 | 0,088033 | 0,063157 | 0,090286 | PCO_17:2-18:1 |
| PCO_17:2-18:2 | PC O- | MEM | 35 | 4 | 0 | 6,31679 | 5,79693 | 5,66668 | 4,74808 | 5,60302 | 5,33445 | 3,29877 | 3,9082 | 6,21407 | 6,17786 | 3,56835 | 2,43759 | PCO_17:2-18:2 |
| PCO_18:0-14:0 | PC O- | MEM | 32 | 0 | 0 | 2,68708 | 4,49594 | 4,04216 | 2,5795 | 2,53618 | 3,34851 | 1,49276 | 2,46089 | 3,87777 | 4,67269 | 3,0421 | 2,91937 | PCO_18:0-14:0 |
| PCO_18:0-15:0 | PC O- | MEM | 33 | 0 | 0 | 0,005779 | 0,010429 | 0,003185 | 0,00862 | 0,004465 | 0 | 0 | 0,002172 | 0,014201 | 0,0061 | 0,006907 | 0,007653 | PCO_18:0-15:0 |
| PCO_18:0-16:0 | PC O- | MEM | 34 | 0 | 0 | 7,62721 | 9,6372 | 9,83617 | 6,62331 | 7,86897 | 7,6384 | 3,41636 | 5,20322 | 11,2101 | 10,0916 | 5,56914 | 5,61436 | PCO_18:0-16:0 |
| PCO_18:0-16:1 | PC O- | MEM | 34 | 1 | 0 | 1,20386 | 1,38948 | 1,73849 | 1,01076 | 1,14973 | 1,37527 | 0,637611 | 0,660357 | 1,1919 | 1,13076 | 0,720622 | 0,835785 | PCO_18:0-16:1 |
| PCO_18:0-18:0 | PC O- | MEM | 36 | 0 | 0 | 0 | 0 | 0 | 0 | 0,975414 | 0 | 0 | 0 | 1,23709 | 0,834058 | 0 | 0 | PCO_18:0-18:0 |
| PCO_18:0-18:1 | PC O- | MEM | 36 | 1 | 0 | 2,93385 | 4,3254 | 4,14328 | 2,70394 | 3,65765 | 3,01522 | 1,21363 | 1,99618 | 3,90207 | 4,04705 | 2,25968 | 2,40123 | PCO_18:0-18:1 |
| PCO_18:0-18:2 | PC O- | MEM | 36 | 2 | 0 | 10,1774 | 12,723 | 12,3666 | 10,247 | 11,2123 | 9,05579 | 3,77407 | 7,08488 | 16,5577 | 12,1362 | 8,69288 | 8,38739 | PCO_18:0-18:2 |
| PCO_18:0-20:1 | PC O- | MEM | 38 | 1 | 0 | 0 | 0,986604 | 1,31545 | 0 | 0 | 0 | 0 | 0 | 0 | 0 | 0 | 0 | PCO_18:0-20:1 |
| PCO_18:0-20:2 | PC O- | MEM | 38 | 2 | 0 | 0 | 2,08897 | 2,65042 | 1,69692 | 1,6869 | 2,31087 | 0 | 1,51975 | 2,38487 | 1,8598 | 1,07121 | 1,35072 | PCO_18:0-20:2 |
| PCO_18:0-20:3 | PC O- | MEM | 38 | 3 | 0 | 1,1626 | 0,960546 | 1,64286 | 1,07864 | 1,42555 | 1,04529 | 0,655497 | 0,787335 | 1,78376 | 1,40015 | 1,19794 | 1,13481 | PCO_18:0-20:3 |
| PCO_18:0-20:4 | PC O- | MEM | 38 | 4 | 0 | 18,05 | 19,6377 | 16,8034 | 12,2144 | 17 | 14,8374 | 7,08484 | 9,28106 | 18,1293 | 18,8384 | 9,91161 | 9,99751 | PCO_18:0-20:4 |
| PCO_18:0-20:5 | PC O- | MEM | 38 | 5 | 0 | 0 | 0 | 0,472853 | 0 | 0 | 0,559076 | 0 | 0,42767 | 0 | 0,593724 | 0,488746 | 0 | PCO_18:0-20:5 |
| PCO_18:0-20:6 | PC O- | MEM | 38 | 6 | 0 | 0 | 1,17005 | 1,33576 | 0,836699 | 1,3725 | 1,10993 | 0,922434 | 0 | 1,63303 | 0 | 0,644705 | 1,16854 | PCO_18:0-20:6 |
| PCO_18:0-22:4 | PC O- | MEM | 40 | 4 | 0 | 2,11162 | 2,90482 | 2,40785 | 1,61121 | 2,11083 | 2,22672 | 0,918045 | 0,750945 | 1,02234 | 3,14054 | 0,859087 | 1,50554 | PCO_18:0-22:4 |
| PCO_18:0-22:5 | PC O- | MEM | 40 | 5 | 0 | 0 | 0 | 0 | 0,623827 | 0 | 2,26732 | 0 | 0 | 0,541682 | 0 | 0 | 0 | PCO_18:0-22:5 |
| PCO_18:1-14:0 | PC O- | MEM | 32 | 1 | 0 | 1,46293 | 2,02969 | 2,61145 | 1,50606 | 1,61536 | 1,34201 | 0,721343 | 1,60791 | 2,31761 | 2,3439 | 1,39318 | 1,51483 | PCO_18:1-14:0 |
| PCO_18:1-15:0 | PC O- | MEM | 33 | 1 | 0 | 0,040433 | 0,073806 | 0,103698 | 0,057868 | 0,050959 | 0,084094 | 0,023424 | 0,04025 | 0,09648 | 0 | 0,04134 | 0,034903 | PCO_18:1-15:0 |
| PCO_18:1-16:0 | PC O- | MEM | 34 | 1 | 0 | 57,3521 | 64,6886 | 70,9697 | 47,6295 | 53,6794 | 51,7229 | 29,4786 | 40,633 | 57,9868 | 69,4621 | 36,493 | 41,1897 | PCO_18:1-16:0 |
| PCO_18:1-16:1 | PC O- | MEM | 34 | 2 | 0 | 2,02083 | 2,11053 | 3,03649 | 1,71713 | 1,623 | 1,70322 | 1,01822 | 1,53561 | 2,0676 | 2,72496 | 1,3008 | 1,78614 | PCO_18:1-16:1 |
| PCO_18:1-17:0 | PC O- | MEM | 35 | 1 | 0 | 0 | 0 | 0 | 0 | 0 | 0 | 0 | 0 | 0,00394 | 0 | 0 | 0,001526 | PCO_18:1-17:0 |
| PCO_18:1-18:0 | PC O- | MEM | 36 | 1 | 0 | 0,253338 | 0,682282 | 0,505772 | 0,379 | 0,383043 | 0,393457 | 0,101043 | 0,312936 | 0,556778 | 0,521948 | 0,296199 | 0,26268 | PCO_18:1-18:0 |
| PCO_18:1-18:1 | PC O- | MEM | 36 | 2 | 0 | 5,48703 | 6,77609 | 8,73118 | 5,8277 | 5,60497 | 4,83791 | 2,68484 | 3,96665 | 8,20601 | 7,96291 | 4,24228 | 4,80636 | PCO_18:1-18:1 |
| PCO_18:1-18:2 | PC O- | MEM | 36 | 3 | 0 | 14,8953 | 23,548 | 23,1675 | 18,4839 | 18,0915 | 14,5824 | 8,15811 | 15,5815 | 26,5282 | 22,2481 | 14,7421 | 17,3138 | PCO_18:1-18:2 |
| PCO_18:1-20:2 | PC O- | MEM | 38 | 3 | 0 | 0,291462 | 0,636652 | 0,875788 | 0,474737 | 0,337583 | 0,496765 | 0,140016 | 0,189373 | 0,643591 | 0,346431 | 0,259289 | 0,235594 | PCO_18:1-20:2 |
| PCO_18:1-20:3 | PC O- | MEM | 38 | 4 | 0 | 3,37531 | 3,98427 | 4,31729 | 2,81315 | 3,85885 | 3,73812 | 2,15069 | 1,90505 | 5,01629 | 3,96238 | 2,57788 | 2,17828 | PCO_18:1-20:3 |
| PCO_18:1-20:4 | PC O- | MEM | 38 | 5 | 0 | 48,4005 | 48,1282 | 49,4183 | 34,0595 | 41,3459 | 42,0693 | 23,788 | 27,2661 | 37,1175 | 48,5327 | 24,1251 | 26,9466 | PCO_18:1-20:4 |
| PCO_18:1-22:4 | PC O- | MEM | 40 | 5 | 0 | 3,96233 | 4,23651 | 9,33018 | 2,32367 | 5,95304 | 8,87748 | 2,37139 | 3,31557 | 2,01084 | 5,55192 | 2,69122 | 2,03567 | PCO_18:1-22:4 |
| PCO_18:1-22:5 | PC O- | MEM | 40 | 6 | 0 | 0 | 0 | 0 | 0 | 3,62364 | 4,81405 | 0 | 0 | 0 | 4,8689 | 0 | 0 | PCO_18:1-22:5 |
| PCO_18:2-14:0 | PC O- | MEM | 32 | 2 | 0 | 0 | 0 | 0,573839 | 0,597836 | 0,737116 | 0,853047 | 0,338583 | 0,380949 | 1,38342 | 0,8341 | 0,64673 | 0,511745 | PCO_18:2-14:0 |
| PCO_18:2-16:0 | PC O- | MEM | 34 | 2 | 0 | 9,79006 | 11,9846 | 11,3593 | 8,35714 | 8,68493 | 9,91818 | 5,7988 | 8,97588 | 10,2006 | 10,5422 | 6,64182 | 7,19331 | PCO_18:2-16:0 |
| PCO_18:2-16:1 | PC O- | MEM | 34 | 3 | 0 | 0 | 0,367733 | 0,203194 | 0,207422 | 0 | 0,230074 | 0,11486 | 0,370321 | 0,24535 | 0,286141 | 0,162596 | 0,151086 | PCO_18:2-16:1 |
| PCO_18:2-17:0 | PC O- | MEM | 35 | 2 | 0 | 0 | 0 | 0 | 0 | 0 | 0 | 0 | 0 | 0 | 0 | 0,004507 | 0,005481 | PCO_18:2-17:0 |
| PCO_18:2-18:0 | PC O- | MEM | 36 | 2 | 0 | 1,31049 | 1,16281 | 1,19934 | 0,822037 | 1,14381 | 1,15454 | 0,665619 | 0,768766 | 1,16618 | 1,08007 | 0,64845 | 0,877579 | PCO_18:2-18:0 |
| PCO_18:2-18:1 | PC O- | MEM | 36 | 3 | 0 | 1,26895 | 2,3829 | 2,24604 | 1,96071 | 1,65582 | 1,79232 | 1,16253 | 1,66353 | 2,2165 | 2,357 | 1,33185 | 1,31617 | PCO_18:2-18:1 |
| PCO_18:2-18:2 | PC O- | MEM | 36 | 4 | 0 | 8,12966 | 11,3447 | 10,3064 | 8,32679 | 9,25284 | 10,2253 | 4,15114 | 9,59335 | 13,6046 | 9,53019 | 7,09076 | 7,86076 | PCO_18:2-18:2 |
| PCO_18:2-20:2 | PC O- | MEM | 38 | 4 | 0 | 0 | 0,293988 | 0 | 0 | 0 | 0,313481 | 0 | 0,216707 | 0 | 0 | 0 | 0 | PCO_18:2-20:2 |
| PCO_18:2-20:3 | PC O- | MEM | 38 | 5 | 0 | 0 | 0 | 0 | 0,43856 | 0 | 0 | 0,417858 | 0 | 0,610468 | 0,88487 | 0 | 0,348287 | PCO_18:2-20:3 |
| PCO_18:2-20:4 | PC O- | MEM | 38 | 6 | 0 | 8,25588 | 8,01653 | 6,71774 | 5,34979 | 6,01022 | 7,57576 | 4,87553 | 6,57948 | 7,02018 | 9,49682 | 4,3273 | 4,15726 | PCO_18:2-20:4 |
| PE_14:0-18:0 | PE | MEM | 32 | 0 | 0 | 0,12243 | 0 | 0,165923 | 0 | 0 | 0,148267 | 0 | 0 | 0,078885 | 0 | 0 | 0 | PE_14:0-18:0 |
| PE_14:0-18:1 | PE | MEM | 32 | 1 | 0 | 0 | 0,099396 | 0,246979 | 0 | 0 | 0,255512 | 0,160085 | 0,111299 | 0 | 0 | 0 | 0 | PE_14:0-18:1 |

| | | | | | | | | | | | | | | | | | | |
|---|---|---|---|---|---|---|---|---|---|---|---|---|---|---|---|---|---|---|
| PE_16:0-16:0 | PE | MEM | 32 | 0 | 0 | 1,29357 | 0,933893 | 1,20878 | 0,850354 | 0,962165 | 1,72349 | 1,30663 | 0,957614 | 0 | 1,20733 | 0,588463 | 0,66588 | PE_16:0-16:0 |
| PE_16:0-16:1 | PE | MEM | 32 | 1 | 0 | 0,973448 | 0,628147 | 1,1378 | 0,59253 | 0,794501 | 1,12837 | 0,902092 | 0,627603 | 0,33482 | 1,14109 | 0,382731 | 0,449798 | PE_16:0-16:1 |
| PE_16:0-18:0 | PE | MEM | 34 | 0 | 0 | 2,16196 | 1,81951 | 2,9628 | 1,31322 | 1,76658 | 3,36229 | 2,39062 | 1,38961 | 0,854873 | 2,06193 | 1,11457 | 1,14802 | PE_16:0-18:0 |
| PE_16:0-18:1 | PE | MEM | 34 | 1 | 0 | 11,4153 | 9,11057 | 14,5985 | 7,26165 | 9,06193 | 15,7136 | 12,7984 | 6,94205 | 4,7482 | 11,1091 | 5,23713 | 5,50821 | PE_16:0-18:1 |
| PE_16:0-18:2 | PE | MEM | 34 | 2 | 0 | 8,29804 | 7,63357 | 9,03928 | 5,43365 | 6,63317 | 11,041 | 8,32639 | 6,01167 | 4,67605 | 7,93641 | 3,98945 | 4,10659 | PE_16:0-18:2 |
| PE_16:0-18:3 | PE | MEM | 34 | 3 | 0 | 0,198267 | 0 | 0 | 0 | 0,123319 | 0 | 0,231537 | 0,199288 | 0 | 0 | 0 | 0 | PE_16:0-18:3 |
| PE_16:0-20:1 | PE | MEM | 36 | 1 | 0 | 0 | 0,506094 | 0,632019 | 0 | 0 | 0,674433 | 0,578074 | 0,311355 | 0,284185 | 0 | 0,235649 | 0 | PE_16:0-20:1 |
| PE_16:0-20:2 | PE | MEM | 36 | 2 | 0 | 1,29235 | 1,24466 | 1,42021 | 0,870525 | 1,12564 | 1,9159 | 1,11946 | 1,0504 | 0,664348 | 1,20226 | 0,688529 | 0,586325 | PE_16:0-20:2 |
| PE_16:0-20:3 | PE | MEM | 36 | 3 | 0 | 2,46957 | 2,0403 | 2,56264 | 1,35915 | 1,76515 | 3,08921 | 2,97908 | 1,77818 | 1,0809 | 2,22046 | 1,02626 | 0,983551 | PE_16:0-20:3 |
| PE_16:0-20:4 | PE | MEM | 36 | 4 | 0 | 35,3365 | 23,2688 | 38,8576 | 17,0801 | 25,6598 | 44,5563 | 41,204 | 19,7609 | 8,23993 | 29,3879 | 13,5592 | 13,6805 | PE_16:0-20:4 |
| PE_16:0-22:4 | PE | MEM | 38 | 4 | 0 | 15,0312 | 10,6198 | 14,1032 | 7,0773 | 10,0075 | 18,5701 | 13,468 | 9,00509 | 3,65489 | 10,4027 | 6,05845 | 5,73195 | PE_16:0-22:4 |
| PE_16:0-22:5 | PE | MEM | 38 | 5 | 0 | 15,7646 | 12,897 | 19,0884 | 10,5226 | 13,1651 | 21,6196 | 17,2158 | 10,7691 | 0 | 16,0028 | 7,77242 | 8,33915 | PE_16:0-22:5 |
| PE_16:0-22:6 | PE | MEM | 38 | 6 | 0 | 14,9176 | 8,88139 | 15,3766 | 8,19295 | 10,9579 | 21,0307 | 14,6355 | 8,2238 | 0 | 12,7781 | 5,49964 | 6,85282 | PE_16:0-22:6 |
| PE_16:1-18:0 | PE | MEM | 34 | 1 | 0 | 0,921168 | 0,851581 | 1,41872 | 0,629397 | 0,816077 | 1,07305 | 0,734808 | 0,586492 | 0,559372 | 0,935319 | 0,499654 | 0,393072 | PE_16:1-18:0 |
| PE_16:1-18:1 | PE | MEM | 34 | 2 | 0 | 1,44441 | 0,979895 | 1,94764 | 0,892099 | 1,00768 | 1,70116 | 1,35932 | 0,833317 | 0,663864 | 1,32165 | 0,674136 | 0,697342 | PE_16:1-18:1 |
| PE_16:1-18:2 | PE | MEM | 34 | 3 | 0 | 0,454817 | 0,414405 | 0,739806 | 0,345473 | 0,26609 | 1,08879 | 0,553818 | 0,53512 | 0 | 0,705154 | 0,324316 | 0,379744 | PE_16:1-18:2 |
| PE_16:1-20:3 | PE | MEM | 36 | 4 | 0 | 0,27927 | 0 | 0 | 0 | 0 | 0,802265 | 0,814901 | 0 | 0 | 0 | 0 | 0 | PE_16:1-20:3 |
| PE_16:1-20:4 | PE | MEM | 36 | 5 | 0 | 2,64068 | 1,70041 | 3,95128 | 1,52585 | 1,10401 | 4,4791 | 4,25638 | 2,01396 | 0,348728 | 2,48568 | 0,935628 | 1,13069 | PE_16:1-20:4 |
| PE_16:1-22:4 | PE | MEM | 38 | 5 | 0 | 1,18222 | 0 | 0,980008 | 0,547327 | 0,41195 | 1,35532 | 1,22634 | 0,744238 | 0 | 0 | 0 | 0,2895 | PE_16:1-22:4 |
| PE_17:0-18:2 | PE | MEM | 35 | 2 | 0 | 3,05253 | 0 | 0 | 0 | 0 | 0 | 0 | 2,68588 | 0 | 0 | 0 | 2,72563 | PE_17:0-18:2 |
| PE_17:0-20:2 | PE | MEM | 37 | 2 | 0 | 0 | 0 | 0,371961 | 0 | 0 | 0,088475 | 0 | 0 | 0 | 0 | 0 | 0 | PE_17:0-20:2 |
| PE_17:0-20:4 | PE | MEM | 37 | 4 | 0 | 1,80377 | 1,02121 | 1,69195 | 0,923727 | 1,02685 | 2,22241 | 1,64383 | 1,02489 | 0,549153 | 1,16154 | 0,664195 | 0,62446 | PE_17:0-20:4 |
| PE_17:1-17:1 | PE | MEM | 34 | 2 | 0 | 0 | 0 | 0 | 0,060667 | 0 | 0 | 0,066591 | 0,058531 | 0 | 0 | 0 | 0,051847 | PE_17:1-17:1 |
| PE_17:1-22:6 | PE | MEM | 39 | 7 | 0 | 0 | 0 | 0 | 0 | 0 | 0 | 0 | 0 | 0 | 0 | 0,440302 | 0,560019 | PE_17:1-22:6 |
| PE_18:0-18:1 | PE | MEM | 36 | 1 | 0 | 10,0468 | 9,0714 | 11,6874 | 7,22402 | 9,17252 | 9,48025 | 5,7213 | 5,36426 | 7,3947 | 10,923 | 5,06468 | 5,64579 | PE_18:0-18:1 |
| PE_18:0-18:2 | PE | MEM | 36 | 2 | 0 | 23,6261 | 24,5378 | 25,5516 | 18,1178 | 21,2056 | 22,9259 | 13,3705 | 16,2252 | 21,9062 | 23,3949 | 13,1141 | 13,7753 | PE_18:0-18:2 |
| PE_18:0-18:3 | PE | MEM | 36 | 3 | 0 | 0 | 0,188234 | 0,345012 | 0 | 0,286089 | 0,527061 | 0 | 0,180557 | 0 | 0 | 0 | 0,216085 | PE_18:0-18:3 |
| PE_18:0-20:1 | PE | MEM | 38 | 1 | 0 | 0 | 0 | 0,427734 | 0 | 0 | 0 | 0 | 0 | 0,4017 | 0 | 0 | 0 | PE_18:0-20:1 |
| PE_18:0-20:2 | PE | MEM | 38 | 2 | 0 | 0,436565 | 0 | 0,399933 | 0,288548 | 0,375496 | 0 | 0,40435 | 0,369309 | 0,318353 | 0,329602 | 0,195663 | 0,179456 | PE_18:0-20:2 |
| PE_18:0-20:3 | PE | MEM | 38 | 3 | 0 | 4,42325 | 4,25972 | 6,31193 | 3,4402 | 4,52405 | 4,59906 | 3,76067 | 3,30425 | 4,1748 | 4,87592 | 2,56832 | 2,43657 | PE_18:0-20:3 |
| PE_18:0-20:4 | PE | MEM | 38 | 4 | 0 | 134,608 | 123,12 | 145,902 | 83,1458 | 109,87 | 145,224 | 87,7717 | 73,9306 | 71,8238 | 125,184 | 60,7514 | 65,481 | PE_18:0-20:4 |
| PE_18:0-20:5 | PE | MEM | 38 | 5 | 0 | 0,869448 | 0 | 1,25699 | 0 | 0 | 1,49823 | 0 | 0,501922 | 0 | 0 | 0,412859 | 0 | PE_18:0-20:5 |
| PE_18:0-22:4 | PE | MEM | 40 | 4 | 0 | 13,9731 | 11,3456 | 10,3515 | 7,02598 | 10,4563 | 11,1695 | 6,68269 | 3,20912 | 4,42402 | 11,9111 | 4,876 | 5,79055 | PE_18:0-22:4 |
| PE_18:0-22:5 | PE | MEM | 40 | 5 | 0 | 5,01673 | 4,2396 | 6,10041 | 3,51082 | 4,12556 | 6,14713 | 4,16473 | 2,88798 | 2,16722 | 5,00963 | 2,14227 | 2,60218 | PE_18:0-22:5 |
| PE_18:0-22:6 | PE | MEM | 40 | 6 | 0 | 9,39248 | 7,60095 | 8,46621 | 5,05555 | 7,68993 | 11,4815 | 6,13923 | 4,42763 | 4,38034 | 8,24117 | 3,38483 | 4,15085 | PE_18:0-22:6 |
| PE_18:1-18:1 | PE | MEM | 36 | 2 | 0 | 5,78743 | 5,64971 | 8,65692 | 4,20764 | 5,68538 | 7,33571 | 5,62434 | 3,87613 | 4,08176 | 6,3401 | 2,94159 | 3,50408 | PE_18:1-18:1 |
| PE_18:1-18:2 | PE | MEM | 36 | 3 | 0 | 7,73262 | 6,5514 | 9,72433 | 5,23064 | 6,06684 | 10,7715 | 9,19722 | 5,73032 | 4,30723 | 7,47341 | 3,54885 | 4,14868 | PE_18:1-18:2 |
| PE_18:1-18:3 | PE | MEM | 36 | 4 | 0 | 1,73641 | 1,65278 | 1,89959 | 0,959825 | 0 | 2,08159 | 1,9074 | 1,06513 | 0 | 1,24302 | 0 | 0,614402 | PE_18:1-18:3 |
| PE_18:1-20:1 | PE | MEM | 38 | 2 | 0 | 0,206763 | 0 | 0,254839 | 0,148374 | 0,223062 | 0 | 0,234517 | 0 | 0,067156 | 0,195138 | 0,099864 | 0,085359 | PE_18:1-20:1 |
| PE_18:1-20:2 | PE | MEM | 38 | 3 | 0 | 0,609119 | 0,601808 | 0,952031 | 0,466525 | 0,539632 | 0,87124 | 0,662446 | 0,628252 | 0,455107 | 0,636702 | 0,336407 | 0,430741 | PE_18:1-20:2 |
| PE_18:1-20:3 | PE | MEM | 38 | 4 | 0 | 5,16368 | 4,97801 | 7,19303 | 3,09777 | 3,91725 | 6,84098 | 6,52356 | 3,19327 | 2,57326 | 4,64317 | 2,18705 | 2,51301 | PE_18:1-20:3 |
| PE_18:1-20:4 | PE | MEM | 38 | 5 | 0 | 37,8287 | 26,4729 | 46,7565 | 17,6631 | 24,5378 | 54,1963 | 56,268 | 19,5078 | 10,657 | 31,2034 | 12,4109 | 13,4618 | PE_18:1-20:4 |
| PE_18:1-22:3 | PE | MEM | 40 | 4 | 0 | 0 | 0 | 2,16807 | 0 | 0 | 2,5859 | 1,5037 | 3,19333 | 0 | 0 | 0 | 0 | PE_18:1-22:3 |
| PE_18:1-22:4 | PE | MEM | 40 | 5 | 0 | 6,81346 | 5,16625 | 8,08121 | 3,78585 | 4,84744 | 10,4767 | 8,57812 | 4,31414 | 1,44705 | 5,68602 | 2,79099 | 3,123 | PE_18:1-22:4 |
| PE_18:1-22:5 | PE | MEM | 40 | 6 | 0 | 8,31762 | 6,69077 | 9,78675 | 4,49186 | 6,59035 | 10,7254 | 8,79607 | 4,70326 | 0 | 7,19603 | 3,46165 | 3,82063 | PE_18:1-22:5 |
| PE_18:1-22:6 | PE | MEM | 40 | 7 | 0 | 13,0641 | 11,0959 | 19,4999 | 10,0056 | 11,5926 | 21,9845 | 19,9389 | 7,22427 | 0 | 13,7679 | 0 | 7,61571 | PE_18:1-22:6 |
| PE_18:2-18:2 | PE | MEM | 36 | 4 | 0 | 4,90468 | 4,3387 | 5,16338 | 3,48821 | 3,73002 | 6,9268 | 4,53062 | 3,81726 | 2,63535 | 4,01742 | 2,42624 | 2,25873 | PE_18:2-18:2 |
| PE_18:2-20:1 | PE | MEM | 38 | 3 | 0 | 0 | 0 | 0 | 0,202369 | 0 | 0,296929 | 0,282374 | 0,26482 | 0 | 0 | 0 | 0,184935 | PE_18:2-20:1 |
| PE_18:2-20:2 | PE | MEM | 38 | 4 | 0 | 1,77935 | 1,08874 | 1,32005 | 0,778288 | 1,08375 | 1,38491 | 0,603107 | 1,28199 | 0,828031 | 1,27487 | 0,846415 | 1,0457 | PE_18:2-20:2 |
| PE_18:2-20:3 | PE | MEM | 38 | 5 | 0 | 1,18389 | 1,30696 | 1,38242 | 0,331336 | 0 | 1,27765 | 1,25542 | 1,07265 | 0,510948 | 0,594455 | 0,402998 | 0,447068 | PE_18:2-20:3 |
| PE_18:2-20:4 | PE | MEM | 38 | 6 | 0 | 7,90878 | 6,85213 | 9,18515 | 4,02992 | 5,78745 | 10,6551 | 10,1001 | 5,64709 | 4,494 | 6,18749 | 2,92035 | 3,36739 | PE_18:2-20:4 |
| PE_18:2-22:4 | PE | MEM | 40 | 6 | 0 | 2,66171 | 2,19999 | 2,91701 | 1,43472 | 1,58701 | 3,53487 | 2,41926 | 2,20442 | 1,80433 | 2,29648 | 1,05476 | 1,14237 | PE_18:2-22:4 |
| PE_18:2-22:5 | PE | MEM | 40 | 7 | 0 | 4,72214 | 0 | 0 | 0 | 0 | 6,27694 | 0 | 3,3508 | 0 | 0 | 0 | 0 | PE_18:2-22:5 |
| PE_20:1-20:4 | PE | MEM | 40 | 5 | 0 | 0,563636 | 0,412012 | 0,538916 | 0 | 0,384519 | 0,846488 | 0,36864 | 0,471035 | 0 | 0,47369 | 0,231333 | 0,18794 | PE_20:1-20:4 |
| PE_20:2-20:4 | PE | MEM | 40 | 6 | 0 | 1,08859 | 0,572098 | 1,06955 | 0,700262 | 0,532695 | 1,34199 | 0,803415 | 0,720063 | 0,354303 | 0,571489 | 0,28997 | 0,247697 | PE_20:2-20:4 |

| | | | | | | | | | | | | | | | | | | |
|---|---|---|---|---|---|---|---|---|---|---|---|---|---|---|---|---|---|---|
| PE_20:3-20:4 | PE | MEM | 40 | 7 | 0 | 0 | 1,00138 | 0,618807 | 0 | 0 | 0,498633 | 0 | 0,460366 | 0 | 0 | 0 | 0 | PE_20:3-20:4 |
| PE_20:4-20:4 | PE | MEM | 40 | 8 | 0 | 2,90883 | 3,93844 | 3,09176 | 1,44292 | 2,07602 | 4,88014 | 3,08927 | 2,0401 | 0,634622 | 0 | 1,02207 | 1,29528 | PE_20:4-20:4 |
| PE_20:4-22:4 | PE | MEM | 42 | 8 | 0 | 1,02094 | 0,937013 | 1,29009 | 0 | 0 | 1,34492 | 0 | 0 | 0 | 0 | 0 | 0 | PE_20:4-22:4 |
| PEO_16:0-16:0 | PE O- | MEM | 32 | 0 | 0 | 0,576543 | 0,787474 | 0,987411 | 0,600685 | 0,71354 | 0,666137 | 0,249534 | 0,543807 | 1,21848 | 0,765585 | 0,541828 | 0,511995 | PEO_16:0-16:0 |
| PEO_16:0-16:1 | PE O- | MEM | 32 | 1 | 0 | 0,070312 | 0,104945 | 0,127981 | 0,059997 | 0,081461 | 0 | 0,051807 | 0,049309 | 0,098837 | 0,219285 | 0,069823 | 0,049221 | PEO_16:0-16:1 |
| PEO_16:0-18:0 | PE O- | MEM | 34 | 0 | 0 | 0 | 0 | 0 | 0 | 0 | 0,033908 | 0 | 0 | 0 | 0 | 0 | 0,015134 | PEO_16:0-18:0 |
| PEO_16:0-18:1 | PE O- | MEM | 34 | 1 | 0 | 2,20762 | 2,61313 | 3,48755 | 2,40743 | 2,85778 | 1,9758 | 1,34474 | 1,88567 | 3,9421 | 2,84754 | 2,08052 | 1,85968 | PEO_16:0-18:1 |
| PEO_16:0-18:2 | PE O- | MEM | 34 | 2 | 0 | 2,64785 | 3,58346 | 4,21351 | 3,26093 | 3,32672 | 2,46184 | 1,66484 | 2,87159 | 6,00156 | 3,25361 | 3,10658 | 2,48498 | PEO_16:0-18:2 |
| PEO_16:0-20:1 | PE O- | MEM | 36 | 1 | 0 | 0 | 0 | 0 | 0,062975 | 0 | 0 | 0,059649 | 0,039058 | 0 | 0,063931 | 0 | 0,042703 | PEO_16:0-20:1 |
| PEO_16:0-20:2 | PE O- | MEM | 36 | 2 | 0 | 0,296455 | 0,28183 | 0,098705 | 0,211506 | 0,10932 | 0,129162 | 0,116297 | 0,198434 | 0,278024 | 0,164009 | 0,177805 | 0,074943 | PEO_16:0-20:2 |
| PEO_16:0-20:3 | PE O- | MEM | 36 | 3 | 0 | 0,37232 | 0,32417 | 0,521475 | 0,383471 | 0,401662 | 0,385339 | 0,314971 | 0,274583 | 0,607449 | 0,398646 | 0,24736 | 0,1218 | PEO_16:0-20:3 |
| PEO_16:0-20:4 | PE O- | MEM | 36 | 4 | 0 | 2,91955 | 2,73203 | 3,44588 | 2,46244 | 2,84136 | 3,11843 | 1,85336 | 2,32815 | 3,34132 | 2,99469 | 1,76491 | 1,67981 | PEO_16:0-20:4 |
| PEO_16:0-20:5 | PE O- | MEM | 36 | 5 | 0 | 0,558305 | 0 | 0,476421 | 0,368777 | 0,582687 | 0,685841 | 0,427093 | 0,498221 | 0,563376 | 0,467835 | 0,286845 | 0,394702 | PEO_16:0-20:5 |
| PEO_16:0-22:3 | PE O- | MEM | 38 | 3 | 0 | 0 | 0 | 0 | 0 | 0 | 0 | 0,191896 | 0,179489 | 0 | 0 | 0 | 0 | PEO_16:0-22:3 |
| PEO_16:0-22:4 | PE O- | MEM | 38 | 4 | 0 | 5,03367 | 3,84483 | 4,27748 | 3,04699 | 4,51542 | 5,7369 | 2,65485 | 3,17013 | 1,98658 | 4,22632 | 1,58595 | 2,16735 | PEO_16:0-22:4 |
| PEO_16:0-22:5 | PE O- | MEM | 38 | 5 | 0 | 1,58401 | 1,34631 | 1,38504 | 1,02754 | 1,38348 | 1,955 | 1,05318 | 0,978992 | 0,495057 | 1,8109 | 0,207165 | 0,633303 | PEO_16:0-22:5 |
| PEO_16:0-24:4 | PE O- | MEM | 40 | 4 | 0 | 0 | 0,217013 | 0,2311 | 0 | 0,21065 | 0 | 0 | 0,177659 | 0 | 0 | 0,188369 | 0 | PEO_16:0-24:4 |
| PEO_16:1-16:0 | PE O- | MEM | 32 | 1 | 0 | 1,59617 | 1,92416 | 2,97855 | 1,76664 | 1,93015 | 1,92373 | 1,02678 | 1,55356 | 2,47957 | 2,1964 | 1,62331 | 1,4944 | PEO_16:1-16:0 |
| PEO_16:1-16:1 | PE O- | MEM | 32 | 2 | 0 | 0,466877 | 0,398365 | 0,54986 | 0,452453 | 0,448044 | 0,311081 | 0,238757 | 0,475931 | 0,894096 | 0,655383 | 0,303828 | 0,402936 | PEO_16:1-16:1 |
| PEO_16:1-18:1 | PE O- | MEM | 34 | 2 | 0 | 11,7367 | 13,4224 | 18,1738 | 11,5918 | 13,7568 | 9,40369 | 6,1556 | 9,13532 | 22,5612 | 15,8608 | 10,278 | 9,70689 | PEO_16:1-18:1 |
| PEO_16:1-18:2 | PE O- | MEM | 34 | 3 | 0 | 17,0911 | 22,9445 | 24,4678 | 20,4346 | 21,3004 | 16,2014 | 8,5675 | 17,8496 | 36,477 | 22,2651 | 18,4154 | 16,5273 | PEO_16:1-18:2 |
| PEO_16:1-18:3 | PE O- | MEM | 34 | 4 | 0 | 0 | 0 | 0,424776 | 0,261579 | 0,31857 | 0 | 0 | 0,323358 | 0,296642 | 0,262575 | 0,285969 | 0,228291 | PEO_16:1-18:3 |
| PEO_16:1-19:1 | PE O- | MEM | 35 | 2 | 0 | 0 | 0 | 0 | 0 | 0 | 0 | 0 | 0 | 0 | 0,004902 | 0 | 0,004001 | PEO_16:1-19:1 |
| PEO_16:1-20:1 | PE O- | MEM | 36 | 2 | 0 | 0,40114 | 0,429488 | 0,504227 | 0,240737 | 0,450388 | 0,382406 | 0,135861 | 0,195515 | 0,306077 | 0,366144 | 0,148174 | 0,102579 | PEO_16:1-20:1 |
| PEO_16:1-20:2 | PE O- | MEM | 36 | 3 | 0 | 1,54534 | 1,39597 | 1,83106 | 1,2323 | 1,4319 | 1,44973 | 0,830253 | 1,09372 | 1,88284 | 1,33669 | 1,02833 | 0,844245 | PEO_16:1-20:2 |
| PEO_16:1-20:3 | PE O- | MEM | 36 | 4 | 0 | 2,72219 | 3,12406 | 4,42207 | 2,5365 | 3,16173 | 3,04462 | 2,42074 | 2,36765 | 4,74323 | 3,42666 | 2,22431 | 2,03661 | PEO_16:1-20:3 |
| PEO_16:1-20:4 | PE O- | MEM | 36 | 5 | 0 | 94,4028 | 93,5682 | 110,916 | 70,5213 | 87,9623 | 95,0907 | 61,7547 | 63,6466 | 99,4739 | 94,4606 | 56,4679 | 54,4239 | PEO_16:1-20:4 |
| PEO_16:1-22:2 | PE O- | MEM | 38 | 3 | 0 | 0 | 0,401732 | 0,234138 | 0,238515 | 0,22146 | 0,209058 | 0,167138 | 0 | 0 | 0 | 0 | 0 | PEO_16:1-22:2 |
| PEO_16:1-22:3 | PE O- | MEM | 38 | 4 | 0 | 0 | 0 | 1,18025 | 0 | 0 | 0 | 1,04145 | 0 | 0 | 0 | 0 | 0 | PEO_16:1-22:3 |
| PEO_16:1-22:4 | PE O- | MEM | 38 | 5 | 0 | 35,5198 | 29,0593 | 35,66 | 19,9868 | 31,0551 | 34,2043 | 20,5721 | 17,1463 | 13,102 | 32,4553 | 12,0945 | 13,8402 | PEO_16:1-22:4 |
| PEO_16:1-22:5 | PE O- | MEM | 38 | 6 | 0 | 8,88503 | 7,43211 | 11,2259 | 6,22262 | 7,22221 | 10,0424 | 7,64436 | 5,46171 | 3,07452 | 7,73875 | 2,65232 | 4,13085 | PEO_16:1-22:5 |
| PEO_16:1-22:6 | PE O- | MEM | 38 | 7 | 0 | 23,2572 | 18,8572 | 22,3746 | 14,1428 | 18,9146 | 26,1615 | 15,7903 | 13,3217 | 13,5792 | 19,6439 | 9,31561 | 10,9928 | PEO_16:1-22:6 |
| PEO_16:1-24:4 | PE O- | MEM | 40 | 5 | 0 | 2,65975 | 2,25067 | 2,63427 | 1,40327 | 2,22183 | 2,61622 | 1,58183 | 1,73264 | 2,05686 | 2,64199 | 1,14412 | 0,999115 | PEO_16:1-24:4 |
| PEO_16:1-24:5 | PE O- | MEM | 40 | 6 | 0 | 0,648029 | 0 | 1,07706 | 0 | 0 | 0 | 0,375391 | 0 | 0 | 0 | 0 | 0 | PEO_16:1-24:5 |
| PEO_16:2-16:0 | PE O- | MEM | 32 | 2 | 0 | 0,300756 | 0,467436 | 0,472238 | 0,195334 | 0,238405 | 0,445481 | 0,155875 | 0,236999 | 0,369587 | 0,318741 | 0,230134 | 0,19322 | PEO_16:2-16:0 |
| PEO_16:2-18:0 | PE O- | MEM | 34 | 2 | 0 | 0,166143 | 0,64046 | 0,454481 | 0,254961 | 0,293595 | 0,472525 | 0,244536 | 0,255246 | 0,330481 | 0,276882 | 0,249632 | 0,324691 | PEO_16:2-18:0 |
| PEO_16:2-18:1 | PE O- | MEM | 34 | 3 | 0 | 0 | 0,205772 | 0,112292 | 0 | 0 | 0 | 0,093984 | 0 | 0,110349 | 0,191012 | 0,065949 | 0 | PEO_16:2-18:1 |
| PEO_16:2-18:2 | PE O- | MEM | 34 | 4 | 0 | 0 | 0 | 0 | 0 | 0 | 0 | 0 | 0 | 0,147097 | 0,113347 | 0 | 0 | PEO_16:2-18:2 |
| PEO_17:0-15:0 | PE O- | MEM | 32 | 0 | 0 | 0,051522 | 0,037017 | 0 | 0,051179 | 0,032553 | 0,05028 | 0,068074 | 0,103238 | 0,079762 | 0,080445 | 0,106593 | 0,051378 | PEO_17:0-15:0 |
| PEO_17:0-16:1 | PE O- | MEM | 33 | 1 | 0 | 0 | 0 | 0 | 0 | 0 | 0 | 0 | 0 | 0,086297 | 0,165629 | 0 | 0 | PEO_17:0-16:1 |
| PEO_17:0-18:2 | PE O- | MEM | 35 | 2 | 0 | 0,321513 | 0,425574 | 0,500767 | 0,302392 | 0,384202 | 0 | 0 | 0,356877 | 0,701348 | 0,354552 | 0,282467 | 0,248609 | PEO_17:0-18:2 |
| PEO_17:0-18:3 | PE O- | MEM | 35 | 3 | 0 | 0,016936 | 0 | 0 | 0 | 0,03625 | 0 | 0 | 0,037629 | 0 | 0 | 0 | 0 | PEO_17:0-18:3 |
| PEO_17:0-22:5 | PE O- | MEM | 39 | 5 | 0 | 0,476352 | 0,692487 | 0,6229 | 0,364222 | 0,550201 | 0,772384 | 0 | 0,445592 | 0 | 0,51913 | 0,274541 | 0,392789 | PEO_17:0-22:5 |
| PEO_17:1-16:0 | PE O- | MEM | 33 | 1 | 0 | 0 | 0 | 0 | 0 | 0 | 0 | 0 | 0 | 0,294947 | 0,161064 | 0 | 0 | PEO_17:1-16:0 |
| PEO_17:1-18:1 | PE O- | MEM | 35 | 2 | 0 | 0,067491 | 0,057305 | 0,107784 | 0,060373 | 0,066261 | 0 | 0 | 0,054701 | 0,125914 | 0,07142 | 0,054375 | 0,047611 | PEO_17:1-18:1 |
| PEO_17:1-18:2 | PE O- | MEM | 35 | 3 | 0 | 0,336742 | 0,647507 | 0,477523 | 0,456432 | 0,498154 | 0,46671 | 0 | 0,507306 | 0,839644 | 0,429202 | 0,416267 | 0,397388 | PEO_17:1-18:2 |
| PEO_17:1-20:4 | PE O- | MEM | 37 | 5 | 0 | 2,40643 | 2,29744 | 2,33138 | 2,18662 | 1,98558 | 2,88473 | 1,59158 | 2,06968 | 2,42716 | 2,10223 | 1,19659 | 1,23116 | PEO_17:1-20:4 |
| PEO_17:1-22:4 | PE O- | MEM | 39 | 5 | 0 | 0,343005 | 0 | 0,339509 | 0,333962 | 0,127476 | 0,388738 | 0 | 0,257923 | 0 | 0,180181 | 0,099938 | 0,096226 | PEO_17:1-22:4 |
| PEO_17:2-18:0 | PE O- | MEM | 35 | 2 | 0 | 0,004623 | 0,0241 | 0,028901 | 0,019719 | 0,008493 | 0 | 0 | 0,011058 | 0,048461 | 0,018885 | 0,016664 | 0,020735 | PEO_17:2-18:0 |
| PEO_17:2-18:1 | PE O- | MEM | 35 | 3 | 0 | 0,120373 | 0,116384 | 0,184006 | 0,141834 | 0,078428 | 0,209934 | 0 | 0,11967 | 0,097849 | 0,208515 | 0,111526 | 0,120933 | PEO_17:2-18:1 |
| PEO_17:2-20:3 | PE O- | MEM | 37 | 5 | 0 | 0,217118 | 0 | 0,158378 | 0 | 0 | 0,211 | 0 | 0 | 0 | 0,161672 | 0,096989 | 0,09062 | PEO_17:2-20:3 |
| PEO_18:0-14:0 | PE O- | MEM | 32 | 0 | 0 | 0 | 0,039018 | 0,033135 | 0,040781 | 0 | 0 | 0 | 0,024428 | 0,05373 | 0 | 0,021202 | 0 | PEO_18:0-14:0 |
| PEO_18:0-16:0 | PE O- | MEM | 34 | 0 | 0 | 0,924816 | 1,1526 | 1,78132 | 1,1751 | 1,02662 | 0,904377 | 0,835168 | 0,856019 | 1,76 | 1,36797 | 0,879996 | 0,717796 | PEO_18:0-16:0 |
| PEO_18:0-16:1 | PE O- | MEM | 34 | 1 | 0 | 0,098019 | 0,246277 | 0,228867 | 0,085923 | 0,125511 | 0,090337 | 0,04308 | 0,058143 | 0,19408 | 0,214398 | 0,134865 | 0,086943 | PEO_18:0-16:1 |
| PEO_18:0-18:1 | PE O- | MEM | 36 | 1 | 0 | 2,0827 | 2,58428 | 3,34114 | 2,12329 | 2,84731 | 1,94136 | 1,02231 | 1,29467 | 3,79658 | 2,61983 | 1,83519 | 1,90464 | PEO_18:0-18:1 |

| | | | | | | | | | | | | | | | | | | |
|---|---|---|---|---|---|---|---|---|---|---|---|---|---|---|---|---|---|---|
| PEO_18:0-18:2 | PE O- | MEM | 36 | 2 | 0 | 3,62289 | 5,1459 | 5,51757 | 4,09692 | 5,03993 | 3,41955 | 1,80436 | 3,37865 | 7,78052 | 5,04031 | 3,81775 | 3,46577 | PEO_18:0-18:2 |
| PEO_18:0-20:2 | PE O- | MEM | 38 | 2 | 0 | 1,05536 | 0,733765 | 1,79112 | 0,573827 | 0,479493 | 0,652954 | 0,331588 | 0 | 0,964188 | 1,24901 | 0,359396 | 0,453165 | PEO_18:0-20:2 |
| PEO_18:0-20:3 | PE O- | MEM | 38 | 3 | 0 | 0,581187 | 0 | 1,03696 | 0,707431 | 0,839908 | 0,635958 | 0,317471 | 0,589152 | 1,16286 | 0,984978 | 0,572903 | 0,563173 | PEO_18:0-20:3 |
| PEO_18:0-20:4 | PE O- | MEM | 38 | 4 | 0 | 7,38363 | 7,99303 | 7,82737 | 5,53692 | 6,77824 | 7,31808 | 3,54772 | 4,23353 | 7,65965 | 7,80542 | 4,42768 | 4,59105 | PEO_18:0-20:4 |
| PEO_18:0-20:5 | PE O- | MEM | 38 | 5 | 0 | 0,393892 | 0 | 1,01756 | 0 | 0 | 0 | 0 | 0,226909 | 0,498435 | 0,366772 | 0,24375 | 0,210498 | PEO_18:0-20:5 |
| PEO_18:0-22:4 | PE O- | MEM | 40 | 4 | 0 | 3,54823 | 3,16978 | 3,56272 | 2,28357 | 3,14756 | 3,7582 | 1,86567 | 2,0166 | 2,0506 | 3,43551 | 1,3591 | 1,99507 | PEO_18:0-22:4 |
| PEO_18:0-22:5 | PE O- | MEM | 40 | 5 | 0 | 0,580319 | 0,367332 | 0,463713 | 0,427582 | 0,303973 | 1,0632 | 0,63668 | 0,223059 | 0 | 0 | 0,235515 | 0 | PEO_18:0-22:5 |
| PEO_18:1-14:0 | PE O- | MEM | 32 | 1 | 0 | 0,092115 | 0,233577 | 0,32325 | 0,065855 | 0,0744 | 0,220088 | 0,045257 | 0,105543 | 0,239316 | 0,116121 | 0,157546 | 0,075511 | PEO_18:1-14:0 |
| PEO_18:1-16:0 | PE O- | MEM | 34 | 1 | 0 | 4,56414 | 4,70009 | 7,22331 | 4,10465 | 4,52326 | 4,82077 | 2,80906 | 3,49856 | 4,79051 | 5,51804 | 2,95899 | 3,31539 | PEO_18:1-16:0 |
| PEO_18:1-16:1 | PE O- | MEM | 34 | 2 | 0 | 0,781559 | 0,68501 | 1,21189 | 0,658005 | 0,767166 | 0,633861 | 0,349261 | 0,575929 | 1,04594 | 0,855477 | 0,569664 | 0,614514 | PEO_18:1-16:1 |
| PEO_18:1-17:1 | PE O- | MEM | 35 | 2 | 0 | 0 | 0 | 0 | 0,005544 | 0 | 0 | 0 | 0,005879 | 0 | 0 | 0 | 0,005068 | PEO_18:1-17:1 |
| PEO_18:1-18:1 | PE O- | MEM | 36 | 2 | 0 | 7,60853 | 9,86851 | 12,2366 | 7,95471 | 10,0455 | 6,86195 | 3,87848 | 5,53942 | 15,1395 | 11,2929 | 6,65119 | 6,92773 | PEO_18:1-18:1 |
| PEO_18:1-18:2 | PE O- | MEM | 36 | 3 | 0 | 16,0884 | 24,8864 | 24,1776 | 21,0091 | 23,5876 | 15,3175 | 6,95258 | 16,032 | 34,8921 | 23,0819 | 17,9235 | 17,3167 | PEO_18:1-18:2 |
| PEO_18:1-18:3 | PE O- | MEM | 36 | 4 | 0 | 0,122644 | 0,116178 | 0,162561 | 0,069035 | 0 | 0,086448 | 0 | 0 | 0,177666 | 0,102384 | 0,121502 | 0,104751 | PEO_18:1-18:3 |
| PEO_18:1-20:1 | PE O- | MEM | 38 | 2 | 0 | 0 | 0,566099 | 0 | 0,439017 | 0,699817 | 0,650011 | 0,376062 | 0 | 0,740469 | 0 | 0,520293 | 0,469186 | PEO_18:1-20:1 |
| PEO_18:1-20:2 | PE O- | MEM | 38 | 3 | 0 | 1,19924 | 2,27083 | 1,74093 | 1,18178 | 1,42306 | 1,38171 | 0,589467 | 0,954328 | 1,81145 | 1,45461 | 1,1802 | 1,13941 | PEO_18:1-20:2 |
| PEO_18:1-20:3 | PE O- | MEM | 38 | 4 | 0 | 3,4429 | 3,74565 | 5,00005 | 3,79439 | 3,89045 | 3,33637 | 2,13476 | 2,74657 | 5,68419 | 4,27504 | 2,32611 | 2,79967 | PEO_18:1-20:3 |
| PEO_18:1-20:4 | PE O- | MEM | 38 | 5 | 0 | 61,8573 | 72,9939 | 82,3415 | 54,498 | 73,1814 | 60,9745 | 29,9829 | 36,4217 | 81,0288 | 73,1489 | 37,8611 | 35,2576 | PEO_18:1-20:4 |
| PEO_18:1-20:5 | PE O- | MEM | 38 | 6 | 0 | 0,37733 | 0 | 0,387167 | 0 | 0 | 0,595831 | 0 | 0 | 0 | 0,386427 | 0 | 0,381745 | PEO_18:1-20:5 |
| PEO_18:1-22:4 | PE O- | MEM | 40 | 5 | 0 | 13,3831 | 11,8778 | 13,4288 | 8,24316 | 11,2526 | 14,256 | 7,74343 | 6,69075 | 6,9435 | 13,0763 | 5,82119 | 6,78062 | PEO_18:1-22:4 |
| PEO_18:1-22:5 | PE O- | MEM | 40 | 6 | 0 | 4,54646 | 5,03165 | 4,97801 | 3,08543 | 3,80248 | 5,56875 | 3,57902 | 3,336 | 2,85029 | 5,19879 | 2,14004 | 2,30811 | PEO_18:1-22:5 |
| PEO_18:1-22:6 | PE O- | MEM | 40 | 7 | 0 | 7,60377 | 8,33552 | 7,56222 | 4,32367 | 10,1505 | 10,0068 | 2,56659 | 2,29248 | 7,47664 | 6,82798 | 3,76221 | 6,6891 | PEO_18:1-22:6 |
| PEO_18:1-24:4 | PE O- | MEM | 42 | 5 | 0 | 1,06914 | 1,07556 | 0,993279 | 0,585047 | 0,840413 | 0,951928 | 0,646362 | 0,67692 | 0,944617 | 0,921452 | 0,548055 | 0,582635 | PEO_18:1-24:4 |
| PEO_18:1-24:5 | PE O- | MEM | 42 | 6 | 0 | 0 | 0 | 0 | 0 | 0 | 0,781734 | 0,476727 | 0 | 0 | 0 | 0 | 0 | PEO_18:1-24:5 |
| PEO_18:2-14:0 | PE O- | MEM | 32 | 2 | 0 | 0,045835 | 0,085737 | 0,281553 | 0,087854 | 0,062876 | 0,101948 | 0,064931 | 0,097624 | 0,124469 | 0,192283 | 0,119318 | 0,061446 | PEO_18:2-14:0 |
| PEO_18:2-16:0 | PE O- | MEM | 34 | 2 | 0 | 6,57583 | 7,02037 | 12,1459 | 5,83729 | 6,63751 | 7,71067 | 5,24365 | 5,69779 | 7,84485 | 8,41935 | 4,52646 | 4,96163 | PEO_18:2-16:0 |
| PEO_18:2-16:1 | PE O- | MEM | 34 | 3 | 0 | 0,356222 | 0,491256 | 0,844924 | 0,342962 | 0,407749 | 0,434268 | 0,18008 | 0,358702 | 0,599213 | 0,4747 | 0,28024 | 0,264851 | PEO_18:2-16:1 |
| PEO_18:2-18:0 | PE O- | MEM | 36 | 2 | 0 | 0,270531 | 0,280443 | 0,455934 | 0,181624 | 0,297236 | 0,15614 | 0,133853 | 0,108252 | 0,32974 | 0,384166 | 0,20328 | 0,227024 | PEO_18:2-18:0 |
| PEO_18:2-18:1 | PE O- | MEM | 36 | 3 | 0 | 4,19038 | 5,09325 | 8,45294 | 5,1155 | 5,22614 | 4,86045 | 3,00183 | 3,8609 | 8,47709 | 6,66535 | 4,16428 | 4,2955 | PEO_18:2-18:1 |
| PEO_18:2-18:2 | PE O- | MEM | 36 | 4 | 0 | 9,79969 | 12,4128 | 16,0992 | 11,6338 | 11,3292 | 10,5939 | 6,12108 | 10,5028 | 18,5134 | 13,47 | 9,85322 | 9,36206 | PEO_18:2-18:2 |
| PEO_18:2-18:3 | PE O- | MEM | 36 | 5 | 0 | 0 | 0 | 0 | 0 | 0 | 0 | 0 | 0,228586 | 0 | 0 | 0,206712 | 0 | PEO_18:2-18:3 |
| PEO_18:2-20:1 | PE O- | MEM | 38 | 3 | 0 | 0 | 0 | 0,183052 | 0 | 0 | 0 | 0 | 0 | 0 | 0,196803 | 0 | 0 | PEO_18:2-20:1 |
| PEO_18:2-20:2 | PE O- | MEM | 38 | 4 | 0 | 0,357044 | 0,72626 | 0,869987 | 0,633333 | 0,482736 | 0,63148 | 0,179065 | 0,729194 | 1,09264 | 0,896023 | 0,497755 | 0,39042 | PEO_18:2-20:2 |
| PEO_18:2-20:3 | PE O- | MEM | 38 | 5 | 0 | 2,07157 | 2,43428 | 4,02894 | 1,64504 | 2,66723 | 2,47595 | 1,40285 | 1,51986 | 2,86855 | 2,57588 | 1,41842 | 1,3695 | PEO_18:2-20:3 |
| PEO_18:2-20:4 | PE O- | MEM | 38 | 6 | 0 | 25,4539 | 28,5026 | 40,0668 | 23,0802 | 26,4518 | 30,5123 | 20,9904 | 22,4211 | 37,2853 | 31,7968 | 19,2334 | 19,6596 | PEO_18:2-20:4 |
| PEO_18:2-22:3 | PE O- | MEM | 40 | 5 | 0 | 0 | 0 | 0,203002 | 0 | 0 | 0 | 0 | 0 | 0 | 0,228948 | 0 | 0 | PEO_18:2-22:3 |
| PEO_18:2-22:4 | PE O- | MEM | 40 | 6 | 0 | 6,00589 | 4,91548 | 7,00929 | 4,22931 | 5,20301 | 7,8841 | 4,81804 | 3,7254 | 2,73713 | 5,9511 | 2,76863 | 3,72096 | PEO_18:2-22:4 |
| PEO_18:2-22:5 | PE O- | MEM | 40 | 7 | 0 | 8,3025 | 6,069 | 9,72065 | 6,77451 | 3,19384 | 7,80401 | 7,60562 | 7,07236 | 4,97025 | 7,55206 | 3,85993 | 2,4679 | PEO_18:2-22:5 |
| PEO_18:2-22:6 | PE O- | MEM | 40 | 8 | 0 | 0 | 0 | 6,4677 | 0 | 0 | 7,17224 | 0 | 3,32757 | 0 | 0 | 2,33209 | 0 | PEO_18:2-22:6 |
| PEO_18:2-24:4 | PE O- | MEM | 42 | 6 | 0 | 1,20904 | 1,49853 | 2,08039 | 0 | 0 | 0,738248 | 0,501932 | 0,939124 | 1,18624 | 0 | 0 | 0 | PEO_18:2-24:4 |
| PG_16:0-18:1 | PG | MEM | 34 | 1 | 0 | 5,82348 | 4,96803 | 8,48647 | 4,2921 | 4,74979 | 6,93341 | 5,2244 | 3,74932 | 3,41242 | 5,96761 | 2,99883 | 3,05855 | PG_16:0-18:1 |
| PG_16:0-18:2 | PG | MEM | 34 | 2 | 0 | 0,317616 | 0,764596 | 0,356509 | 0,25919 | 0,245482 | 0,437229 | 0,589418 | 0,269885 | 0,6391 | 0,366507 | 0,45538 | 0,157514 | PG_16:0-18:2 |
| PG_16:0-20:2 | PG | MEM | 36 | 2 | 0 | 0,908979 | 0,909658 | 0,879926 | 0,538599 | 0,878713 | 1,07984 | 0,635818 | 0,507332 | 0,490351 | 0,803028 | 0,460464 | 0,473028 | PG_16:0-20:2 |
| PG_16:0-20:3 | PG | MEM | 36 | 3 | 0 | 0,410478 | 0,405964 | 0,3187 | 0,178734 | 0,214012 | 0,308075 | 0,152591 | 0,3508 | 0,086786 | 0,230333 | 0 | 0,116095 | PG_16:0-20:3 |
| PG_16:0-20:4 | PG | MEM | 36 | 4 | 0 | 1,04584 | 0,927733 | 0,747867 | 0,42571 | 0,463675 | 0,836074 | 0,523058 | 0,625671 | 0,297716 | 0,597665 | 0,279995 | 0,278845 | PG_16:0-20:4 |
| PG_16:1-18:1 | PG | MEM | 34 | 2 | 0 | 0,538318 | 0 | 0,616804 | 0,343827 | 0,367723 | 0,527321 | 0 | 0,348373 | 0 | 0,525378 | 0 | 0,273927 | PG_16:1-18:1 |
| PG_16:1-22:4 | PG | MEM | 38 | 5 | 0 | 0 | 0 | 0 | 0,362719 | 0,588009 | 0 | 0 | 0,21132 | 0,177789 | 0 | 0,218345 | 0,241046 | PG_16:1-22:4 |
| PG_16:1-22:5 | PG | MEM | 38 | 6 | 0 | 0,261043 | 0,16806 | 0,183662 | 0,092631 | 0 | 0,358839 | 0,197976 | 0,100472 | 0 | 0,206324 | 0 | 0,113097 | PG_16:1-22:5 |
| PG_18:0-18:1 | PG | MEM | 36 | 1 | 0 | 0,921497 | 1,21637 | 1,50525 | 0,908833 | 1,06868 | 0,8338 | 0,514639 | 0,909132 | 2,0367 | 1,36809 | 0,96829 | 0,602394 | PG_18:0-18:1 |
| PG_18:0-18:2 | PG | MEM | 36 | 2 | 0 | 0,517414 | 0,645108 | 0,798905 | 0,524113 | 0,614016 | 0,617796 | 0,346142 | 0,528168 | 0,800957 | 0,683946 | 0,449117 | 0,470195 | PG_18:0-18:2 |
| PG_18:0-20:2 | PG | MEM | 38 | 2 | 0 | 0,205107 | 0,251652 | 0,205571 | 0 | 0 | 0 | 0 | 0,150447 | 0,233625 | 0 | 0,168816 | 0 | PG_18:0-20:2 |
| PG_18:0-20:3 | PG | MEM | 38 | 3 | 0 | 0 | 0,111926 | 0,113411 | 0 | 0,288419 | 0 | 0 | 0,080793 | 0,12292 | 0 | 0,077898 | 0 | PG_18:0-20:3 |
| PG_18:0-20:4 | PG | MEM | 38 | 4 | 0 | 0,357227 | 0,288274 | 0,377871 | 0,283902 | 0,30374 | 0,301147 | 0,242999 | 0,245986 | 0 | 0,399799 | 0,194416 | 0,218057 | PG_18:0-20:4 |
| PG_18:1-18:1 | PG | MEM | 36 | 2 | 0 | 1,10511 | 1,24586 | 1,96103 | 0,88554 | 1,09451 | 1,45111 | 1,20391 | 0,87027 | 0,627076 | 1,18444 | 0,552401 | 0,726013 | PG_18:1-18:1 |
| PG_18:1-18:2 | PG | MEM | 36 | 3 | 0 | 1,90529 | 2,19967 | 2,82774 | 1,82345 | 2,04401 | 2,8522 | 2,00603 | 1,72742 | 1,06614 | 2,15053 | 1,11195 | 1,28148 | PG_18:1-18:2 |

| | | | | | | | | | | | | | | | | | | |
|---|---|---|---|---|---|---|---|---|---|---|---|---|---|---|---|---|---|---|
| PG_18:1-20:2 | PG | MEM | 38 | 3 | 0 | 0,303714 | 0,171482 | 0,225788 | 0 | 0 | 0,326235 | 0,204484 | 0,125937 | 0,126312 | 0,278436 | 0,086606 | 0,190798 | PG_18:1-20:2 |
| PG_18:1-20:3 | PG | MEM | 38 | 4 | 0 | 0,420183 | 0,485127 | 0,543741 | 0,211064 | 0,312136 | 0,257998 | 0,336679 | 0,349493 | 0,600525 | 0,345223 | 0,220744 | 0,212552 | PG_18:1-20:3 |
| PG_18:1-20:4 | PG | MEM | 38 | 5 | 0 | 1,67648 | 1,53032 | 2,22544 | 0,957746 | 1,04249 | 2,29238 | 1,48234 | 0,93902 | 0,377573 | 1,83341 | 0,579293 | 0,650648 | PG_18:1-20:4 |
| PG_18:1-22:4 | PG | MEM | 40 | 5 | 0 | 0,625984 | 0,697078 | 0,922218 | 0,475891 | 0,584922 | 0,878474 | 0,61858 | 0,474985 | 0 | 0,617999 | 0,353424 | 0,429216 | PG_18:1-22:4 |
| PG_18:2-18:2 | PG | MEM | 36 | 4 | 0 | 0,783377 | 0,785992 | 1,25785 | 0,815118 | 1,01875 | 1,29961 | 0,950222 | 0,620138 | 0,526001 | 0,940016 | 0,470544 | 0,646581 | PG_18:2-18:2 |
| PG_18:2-20:2 | PG | MEM | 38 | 4 | 0 | 0,120535 | 0,118223 | 0,127106 | 0,067431 | 0,150308 | 0,187335 | 0,102529 | 0,088637 | 0 | 0,114982 | 0,065151 | 0,079469 | PG_18:2-20:2 |
| PG_18:2-20:3 | PG | MEM | 38 | 5 | 0 | 0,336745 | 0,384928 | 0,519281 | 0,201424 | 0,332983 | 0,372159 | 0,351602 | 0,293049 | 0,138758 | 0 | 0,127322 | 0,14377 | PG_18:2-20:3 |
| PG_18:2-20:4 | PG | MEM | 38 | 6 | 0 | 1,5857 | 1,46943 | 2,13471 | 1,31796 | 1,65737 | 2,45754 | 1,2864 | 1,36904 | 0,647687 | 1,25609 | 0,777116 | 0,932943 | PG_18:2-20:4 |
| PG_18:2-22:4 | PG | MEM | 40 | 6 | 0 | 1,24683 | 1,35324 | 0,62638 | 0,970285 | 1,34553 | 1,34336 | 1,1211 | 0,810059 | 0,577329 | 1,22921 | 0,659254 | 0,691811 | PG_18:2-22:4 |
| PG_18:2-22:5 | PG | MEM | 40 | 7 | 0 | 0 | 0 | 2,12136 | 1,51953 | 0 | 1,77252 | 1,23864 | 0 | 0 | 0 | 0 | 0 | PG_18:2-22:5 |
| PG_20:2-20:4 | PG | MEM | 40 | 6 | 0 | 0 | 0 | 0,110309 | 0 | 0 | 0,142108 | 0 | 0,089858 | 0 | 0 | 0 | 0 | PG_20:2-20:4 |
| PG_20:3-20:4 | PG | MEM | 40 | 7 | 0 | 1,99914 | 2,2915 | 0,551832 | 0 | 0 | 0,445063 | 0,295084 | 1,28849 | 0 | 2,03857 | 0 | 0 | PG_20:3-20:4 |
| PG_20:4-20:4 | PG | MEM | 40 | 8 | 0 | 2,48486 | 2,64009 | 2,96754 | 1,89258 | 2,17475 | 2,98813 | 1,72346 | 1,56735 | 1,51947 | 2,19818 | 1,27155 | 1,44511 | PG_20:4-20:4 |
| PG_20:4-22:4 | PG | MEM | 42 | 8 | 0 | 0,824895 | 0,38561 | 1,18941 | 0,64362 | 0,855673 | 1,32457 | 0,710347 | 0,556245 | 0 | 0 | 0,490714 | 0,491617 | PG_20:4-22:4 |
| PG_22:4-22:4 | PG | MEM | 44 | 8 | 0 | 0,176039 | 0 | 0 | 0 | 0,184018 | 0,252975 | 0,182598 | 0 | 0 | 0 | 0,122091 | 0 | PG_22:4-22:4 |
| PG_22:6-22:6 | PG | MEM | 44 | 12 | 0 | 1,10098 | 1,30457 | 1,15587 | 0,776143 | 0 | 0 | 0,494818 | 0,605853 | 1,64988 | 0 | 0 | 0,653965 | PG_22:6-22:6 |
| PI_16:0-18:0 | PI | MEM | 34 | 0 | 0 | 1,2466 | 0,443975 | 1,11732 | 0,372052 | 0,425994 | 1,36873 | 1,4328 | 0,303822 | 0 | 1,93262 | 1,19298 | 0,298982 | PI_16:0-18:0 |
| PI_16:0-18:1 | PI | MEM | 34 | 1 | 0 | 1,71159 | 1,03761 | 3,41967 | 1,06279 | 1,27858 | 2,79701 | 1,97423 | 1,8226 | 0 | 4,1222 | 2,54686 | 1,28566 | PI_16:0-18:1 |
| PI_16:0-18:2 | PI | MEM | 34 | 2 | 0 | 2,21271 | 2,03847 | 2,91845 | 1,97331 | 2,04759 | 2,28153 | 1,00163 | 1,66027 | 0 | 4,68036 | 3,26638 | 1,22863 | PI_16:0-18:2 |
| PI_16:0-20:2 | PI | MEM | 36 | 2 | 0 | 0,376947 | 0,352372 | 0 | 0 | 0,278866 | 0 | 0,214371 | 0 | 0 | 0,781312 | 0,519049 | 0,206267 | PI_16:0-20:2 |
| PI_16:0-20:3 | PI | MEM | 36 | 3 | 0 | 1,55544 | 0,970555 | 2,11545 | 0,958174 | 1,01454 | 1,41756 | 0,907708 | 0,763578 | 0 | 2,36135 | 1,67257 | 0,611053 | PI_16:0-20:3 |
| PI_16:0-20:4 | PI | MEM | 36 | 4 | 0 | 16,1601 | 10,0929 | 16,5729 | 7,53415 | 9,03565 | 15,6609 | 8,79593 | 6,32557 | 0 | 24,5477 | 14,0991 | 4,76954 | PI_16:0-20:4 |
| PI_16:0-22:4 | PI | MEM | 38 | 4 | 0 | 3,30705 | 1,56736 | 2,64785 | 1,0048 | 1,53176 | 4,10713 | 1,61317 | 0,930903 | 0 | 3,93868 | 2,15211 | 0 | PI_16:0-22:4 |
| PI_16:0-22:5 | PI | MEM | 38 | 5 | 0 | 0 | 0 | 0 | 0 | 0 | 0 | 0,580227 | 0 | 0 | 1,23559 | 0,797457 | 0 | PI_16:0-22:5 |
| PI_16:1-18:0 | PI | MEM | 34 | 1 | 0 | 1,71414 | 1,09992 | 2,13771 | 1,11917 | 1,2629 | 2,08158 | 1,38714 | 0 | 0 | 2,80952 | 1,87664 | 0 | PI_16:1-18:0 |
| PI_16:1-18:1 | PI | MEM | 34 | 2 | 0 | 0,702472 | 0,563896 | 1,10345 | 0 | 0,570784 | 0,831856 | 0,520131 | 0,438487 | 0 | 1,62783 | 1,05048 | 0,29403 | PI_16:1-18:1 |
| PI_16:1-18:2 | PI | MEM | 34 | 3 | 0 | 0 | 0 | 0,339308 | 0 | 0 | 0 | 0,272116 | 0 | 0 | 0 | 0,483957 | 0 | PI_16:1-18:2 |
| PI_16:1-20:4 | PI | MEM | 36 | 5 | 0 | 0 | 0 | 0,465045 | 0 | 0 | 0 | 0,27783 | 0 | 0 | 0 | 0 | 0 | PI_16:1-20:4 |
| PI_17:0-17:0 | PI | MEM | 34 | 0 | 0 | 0,244742 | 0,143854 | 0,286954 | 0,176545 | 0,13026 | 0,425356 | 0,246395 | 0,137487 | 0 | 0,267941 | 0,323226 | 0,11247 | PI_17:0-17:0 |
| PI_17:0-20:4 | PI | MEM | 37 | 4 | 0 | 0,914555 | 0,510302 | 0,950001 | 0,47633 | 0,453845 | 1,07497 | 0,483955 | 0,354068 | 0 | 1,04051 | 0,792717 | 0,294796 | PI_17:0-20:4 |
| PI_17:1-17:1 | PI | MEM | 34 | 2 | 0 | 0 | 0 | 0 | 0,028542 | 0,033693 | 0 | 0,014536 | 0 | 0 | 0 | 0,04645 | 0 | PI_17:1-17:1 |
| PI_18:0-18:1 | PI | MEM | 36 | 1 | 0 | 2,12181 | 1,44117 | 3,37992 | 1,74229 | 1,91842 | 1,82274 | 0,847566 | 0,903183 | 0 | 3,81487 | 3,27363 | 1,14256 | PI_18:0-18:1 |
| PI_18:0-18:2 | PI | MEM | 36 | 2 | 0 | 9,98338 | 8,37259 | 14,1854 | 7,88861 | 8,13253 | 9,9111 | 4,6297 | 5,99949 | 0 | 17,4021 | 14,6563 | 5,6297 | PI_18:0-18:2 |
| PI_18:0-18:3 | PI | MEM | 36 | 3 | 0 | 0 | 0,217538 | 0 | 0 | 0 | 0 | 0,437057 | 0 | 0 | 0,642637 | 0 | 0 | PI_18:0-18:3 |
| PI_18:0-20:3 | PI | MEM | 38 | 3 | 0 | 0 | 2,34691 | 6,03285 | 2,38288 | 2,13363 | 2,60554 | 1,7801 | 1,48712 | 0 | 4,00485 | 3,09749 | 1,17215 | PI_18:0-20:3 |
| PI_18:0-20:4 | PI | MEM | 38 | 4 | 0 | 231,567 | 141,808 | 239,507 | 109,951 | 134,574 | 226,961 | 111,554 | 87,0697 | 0 | 324,377 | 201,621 | 75,0759 | PI_18:0-20:4 |
| PI_18:0-20:5 | PI | MEM | 38 | 5 | 0 | 0,974147 | 0 | 0 | 0 | 0 | 1,378 | 0,750807 | 0 | 0 | 1,44833 | 0 | 0 | PI_18:0-20:5 |
| PI_18:0-22:4 | PI | MEM | 40 | 4 | 0 | 15,6053 | 10,5887 | 16,8648 | 16,3717 | 11,7048 | 16,3431 | 12,4005 | 10,6309 | 0 | 16,346 | 18,983 | 13,6412 | PI_18:0-22:4 |
| PI_18:0-22:5 | PI | MEM | 40 | 5 | 0 | 3,31703 | 1,5514 | 2,91438 | 1,80699 | 1,53089 | 3,65445 | 1,79944 | 2,13536 | 0 | 2,89826 | 2,90713 | 1,35275 | PI_18:0-22:5 |
| PI_18:0-22:6 | PI | MEM | 40 | 6 | 0 | 3,55863 | 1,44862 | 3,44391 | 0 | 0 | 4,76806 | 2,5026 | 0 | 0 | 3,73739 | 0 | 0 | PI_18:0-22:6 |
| PI_18:1-18:1 | PI | MEM | 36 | 2 | 0 | 2,49714 | 1,92348 | 4,70749 | 2,12986 | 2,14515 | 2,56674 | 1,14615 | 1,23073 | 0 | 5,64772 | 3,97907 | 1,32637 | PI_18:1-18:1 |
| PI_18:1-18:2 | PI | MEM | 36 | 3 | 0 | 4,16138 | 3,40589 | 6,6262 | 3,58525 | 3,58641 | 4,73009 | 1,76927 | 2,83474 | 0 | 7,7788 | 6,27432 | 2,25075 | PI_18:1-18:2 |
| PI_18:1-20:2 | PI | MEM | 38 | 3 | 0 | 0 | 0,065466 | 0,211859 | 0,07414 | 0,032965 | 0,080792 | 0,0348 | 0 | 0 | 0,133482 | 0 | 0,030543 | PI_18:1-20:2 |
| PI_18:1-20:3 | PI | MEM | 38 | 4 | 0 | 3,23797 | 2,07171 | 4,20478 | 1,67314 | 1,82086 | 3,23081 | 1,88547 | 1,14735 | 0 | 5,00382 | 3,37178 | 0,999749 | PI_18:1-20:3 |
| PI_18:1-20:4 | PI | MEM | 38 | 5 | 0 | 14,6976 | 9,24127 | 21,016 | 7,79269 | 8,73259 | 16,8466 | 8,64073 | 6,19037 | 0 | 24,2841 | 12,4261 | 4,63741 | PI_18:1-20:4 |
| PI_18:1-22:4 | PI | MEM | 40 | 5 | 0 | 1,11725 | 0,679748 | 1,75096 | 0,628927 | 0,552049 | 1,48517 | 0,822245 | 0 | 0 | 1,62647 | 1,55779 | 0,424361 | PI_18:1-22:4 |
| PI_18:2-18:2 | PI | MEM | 36 | 4 | 0 | 0,917865 | 0,663658 | 0,938113 | 0,487094 | 0,245788 | 0,73048 | 0,362699 | 0,409053 | 0 | 1,48289 | 0,867455 | 0,149757 | PI_18:2-18:2 |
| PI_18:2-20:1 | PI | MEM | 38 | 3 | 0 | 0 | 0,025911 | 0 | 0 | 0 | 0 | 0 | 0 | 0 | 0 | 0,051306 | 0 | PI_18:2-20:1 |
| PI_18:2-20:2 | PI | MEM | 38 | 4 | 0 | 0,440985 | 0,384929 | 0 | 0 | 0 | 0 | 0 | 0 | 0 | 0 | 0,588356 | 0 | PI_18:2-20:2 |
| PI_18:2-20:4 | PI | MEM | 38 | 6 | 0 | 1,12669 | 0,596248 | 1,27356 | 0,511905 | 0 | 0 | 0,782257 | 0,444215 | 0 | 1,58755 | 1,00184 | 0 | PI_18:2-20:4 |
| PI_20:1-20:4 | PI | MEM | 40 | 5 | 0 | 0 | 0,365353 | 0,514176 | 0,331866 | 0,520929 | 0 | 0,380838 | 0 | 0 | 0,96445 | 0 | 0 | PI_20:1-20:4 |
| PI_20:2-20:4 | PI | MEM | 40 | 6 | 0 | 0 | 0,472235 | 1,04045 | 0 | 1,93626 | 0 | 0 | 1,53071 | 0 | 1,2901 | 0 | 0 | PI_20:2-20:4 |
| PS_16:0-16:0 | PS | MEM | 32 | 0 | 0 | 1,69165 | 1,04199 | 2,04861 | 0,930503 | 0,903371 | 1,80283 | 1,23847 | 0,728296 | 0 | 2,18375 | 1,6571 | 0,627414 | PS_16:0-16:0 |
| PS_16:0-16:1 | PS | MEM | 32 | 1 | 0 | 0 | 0,482774 | 1,10842 | 0 | 0 | 0 | 0,381607 | 0 | 0 | 0 | 0 | 0 | PS_16:0-16:1 |
| PS_16:0-18:0 | PS | MEM | 34 | 0 | 0 | 5,18869 | 3,65032 | 6,16517 | 3,34915 | 3,31021 | 6,3687 | 3,67053 | 2,09888 | 2,85322 | 8,13438 | 5,66119 | 2,20681 | PS_16:0-18:0 |

| | | | | | | | | | | | | | | | | | | |
|---|---|---|---|---|---|---|---|---|---|---|---|---|---|---|---|---|---|---|
| PS_16:0-18:1 | PS | MEM | 34 | 1 | 0 | 6,26303 | 5,64101 | 9,98966 | 4,80869 | 5,05517 | 6,7125 | 3,73396 | 2,5963 | 9,26377 | 11,6093 | 8,32764 | 2,87845 | PS_16:0-18:1 |
| PS_16:0-18:2 | PS | MEM | 34 | 2 | 0 | 2,75446 | 2,74327 | 4,37455 | 2,23113 | 2,22892 | 2,87739 | 1,89541 | 1,67406 | 0 | 5,16395 | 5,16161 | 1,35824 | PS_16:0-18:2 |
| PS_16:0-20:2 | PS | MEM | 36 | 2 | 0 | 0 | 0,773889 | 0 | 0 | 0,603213 | 0,739512 | 0 | 0 | 0 | 0 | 0 | 0 | PS_16:0-20:2 |
| PS_16:0-20:3 | PS | MEM | 36 | 3 | 0 | 0 | 1,02507 | 1,54635 | 0,748473 | 0 | 0 | 0,995449 | 0,784163 | 0 | 1,76503 | 0 | 0 | PS_16:0-20:3 |
| PS_16:0-20:4 | PS | MEM | 36 | 4 | 0 | 12,0842 | 6,73008 | 15,7197 | 5,5301 | 5,80643 | 15,6655 | 14,0907 | 4,40846 | 0 | 17,5789 | 13,3604 | 3,25782 | PS_16:0-20:4 |
| PS_16:0-22:4 | PS | MEM | 38 | 4 | 0 | 20,0231 | 20,3826 | 25,4175 | 0 | 16,8701 | 24,2561 | 17,7826 | 0 | 0 | 28,6697 | 24,9261 | 0 | PS_16:0-22:4 |
| PS_16:0-22:5 | PS | MEM | 38 | 5 | 0 | 0 | 0 | 8,74025 | 0 | 0 | 0 | 5,35245 | 0 | 0 | 7,92453 | 0 | 0 | PS_16:0-22:5 |
| PS_16:0-22:6 | PS | MEM | 38 | 6 | 0 | 8,75885 | 0 | 11,2232 | 0 | 4,88645 | 0 | 8,58733 | 0 | 0 | 0 | 8,64435 | 0 | PS_16:0-22:6 |
| PS_16:1-18:0 | PS | MEM | 34 | 1 | 0 | 2,09165 | 2,00361 | 3,40751 | 1,59352 | 1,53558 | 1,82601 | 0,998459 | 0,821515 | 2,27019 | 3,91312 | 3,02055 | 1,06925 | PS_16:1-18:0 |
| PS_17:0-20:4 | PS | MEM | 37 | 4 | 0 | 0 | 0 | 0,749225 | 0 | 0 | 0 | 0,856088 | 0 | 0 | 0,928048 | 0 | 0 | PS_17:0-20:4 |
| PS_17:0-22:4 | PS | MEM | 39 | 4 | 0 | 0 | 0,248018 | 0 | 0 | 0 | 0 | 0 | 0 | 0 | 0,328586 | 0 | 0 | PS_17:0-22:4 |
| PS_17:1-17:1 | PS | MEM | 34 | 2 | 0 | 0 | 0,156223 | 0 | 0 | 0 | 0 | 0 | 0 | 0 | 0 | 0 | 0,0894 | PS_17:1-17:1 |
| PS_18:0-18:1 | PS | MEM | 36 | 1 | 0 | 41,558 | 37,6283 | 58,9156 | 31,5493 | 33,3361 | 35,8383 | 17,652 | 14,458 | 0 | 74,0567 | 46,1355 | 18,8039 | PS_18:0-18:1 |
| PS_18:0-18:2 | PS | MEM | 36 | 2 | 0 | 50,7291 | 57,7422 | 70,4299 | 44,9737 | 43,6913 | 45,0235 | 21,5159 | 24,4878 | 0 | 95,6473 | 84,8105 | 27,7975 | PS_18:0-18:2 |
| PS_18:0-20:2 | PS | MEM | 38 | 2 | 0 | 1,39698 | 1,15047 | 2,09851 | 0 | 1,05122 | 1,67039 | 0,586812 | 0,502046 | 0 | 1,99798 | 1,62264 | 0,669909 | PS_18:0-20:2 |
| PS_18:0-20:3 | PS | MEM | 38 | 3 | 0 | 10,4646 | 10,1469 | 17,7651 | 9,48849 | 9,09459 | 10,4122 | 6,49399 | 5,58288 | 0 | 18,8752 | 15,7455 | 5,52683 | PS_18:0-20:3 |
| PS_18:0-20:4 | PS | MEM | 38 | 4 | 0 | 81,2748 | 59,0978 | 112,803 | 45,189 | 46,3311 | 80,7193 | 35,851 | 23,9725 | 0 | 135,142 | 87,9317 | 0 | PS_18:0-20:4 |
| PS_18:0-22:4 | PS | MEM | 40 | 4 | 0 | 14,4734 | 10,5237 | 17,2163 | 7,69916 | 9,6007 | 14,9521 | 6,58837 | 4,65934 | 0 | 20,5732 | 13,6742 | 4,84773 | PS_18:0-22:4 |
| PS_18:0-22:5 | PS | MEM | 40 | 5 | 0 | 7,40552 | 4,70828 | 9,9738 | 4,56471 | 4,67732 | 8,56048 | 4,68618 | 2,96751 | 0 | 10,345 | 7,68378 | 2,436 | PS_18:0-22:5 |
| PS_18:0-22:6 | PS | MEM | 40 | 6 | 0 | 29,0737 | 19,97 | 36,2407 | 15,1136 | 17,308 | 34,6514 | 16,4864 | 8,41614 | 0 | 40,3463 | 26,6047 | 0 | PS_18:0-22:6 |
| PS_18:0-24:4 | PS | MEM | 42 | 4 | 0 | 0 | 0 | 1,42465 | 0 | 0,505123 | 0 | 0 | 0 | 0 | 1,70026 | 0 | 0 | PS_18:0-24:4 |
| PS_18:1-18:1 | PS | MEM | 36 | 2 | 0 | 2,03087 | 2,22842 | 4,01661 | 1,83651 | 1,96285 | 2,07556 | 1,05742 | 0,920809 | 0 | 4,92293 | 2,96911 | 1,21224 | PS_18:1-18:1 |
| PS_18:1-18:2 | PS | MEM | 36 | 3 | 0 | 3,07214 | 1,01576 | 1,72475 | 0,999076 | 1,96497 | 2,78522 | 0,673942 | 0,605773 | 0 | 2,06675 | 3,14318 | 1,25318 | PS_18:1-18:2 |
| PS_18:1-20:2 | PS | MEM | 38 | 3 | 0 | 0 | 0,267237 | 0,317266 | 0 | 0 | 0 | 0 | 0 | 0 | 0 | 0 | 0 | PS_18:1-20:2 |
| PS_18:1-20:4 | PS | MEM | 38 | 5 | 0 | 7,74229 | 4,15208 | 2,97156 | 3,74322 | 3,48293 | 10,0269 | 3,35268 | 2,83889 | 0 | 2,60779 | 8,24113 | 1,91922 | PS_18:1-20:4 |
| PS_18:1-22:6 | PS | MEM | 40 | 7 | 0 | 0 | 3,25276 | 0 | 0 | 0 | 0 | 5,8626 | 0 | 0 | 0 | 0 | 0 | PS_18:1-22:6 |
| PS_18:2-18:2 | PS | MEM | 36 | 4 | 0 | 0,113643 | 0,062528 | 0,175571 | 0,158976 | 0 | 0 | 0,08895 | 0,064336 | 0 | 0,207247 | 0 | 0 | PS_18:2-18:2 |
| PS_18:2-20:0 | PS | MEM | 38 | 2 | 0 | 0 | 0,081941 | 0 | 1,07026 | 0 | 0 | 0 | 0 | 0 | 0 | 0,322632 | 0 | PS_18:2-20:0 |
| PS_18:2-20:4 | PS | MEM | 38 | 6 | 0 | 0 | 5,3162 | 0 | 4,61532 | 0 | 0 | 0,37734 | 0 | 0 | 13,2325 | 0,5288 | 0 | PS_18:2-20:4 |
| PS_19:0-20:4 | PS | MEM | 39 | 4 | 0 | 0 | 0,43374 | 0 | 0 | 0 | 1,06554 | 1,01646 | 0 | 0 | 0,939862 | 0 | 0 | PS_19:0-20:4 |
| SM_32:1;2 | SM | MEM | 32 | 1 | 2 | 0 | 0 | 0,713295 | 0,449076 | 0 | 0 | 0 | 0 | 0,417089 | 0 | 0 | 0 | SM_32:1;2 |
| SM_34:0;2 | SM | MEM | 34 | 0 | 2 | 9,0707 | 11,6071 | 14,8508 | 7,7377 | 10,0435 | 8,80038 | 4,84978 | 5,19586 | 20,9589 | 10,5234 | 10,7267 | 7,5865 | SM_34:0;2 |
| SM_34:1;2 | SM | MEM | 34 | 1 | 2 | 105,775 | 121,906 | 117,69 | 93,5266 | 104,207 | 90,1478 | 55,7987 | 84,8821 | 123,525 | 115,71 | 79,4955 | 78,572 | SM_34:1;2 |
| SM_34:1;3 | SM | MEM | 34 | 1 | 3 | 0,588974 | 0 | 0 | 0,488363 | 0,672297 | 0 | 0 | 0 | 1,24386 | 0 | 0,539237 | 0,390987 | SM_34:1;3 |
| SM_34:2;2 | SM | MEM | 34 | 2 | 2 | 1,1076 | 1,14409 | 1,12899 | 0,818926 | 1,04851 | 1,14912 | 0,838085 | 0,754133 | 0,90412 | 1,08653 | 0,579982 | 0,626031 | SM_34:2;2 |
| SM_36:0;2 | SM | MEM | 36 | 0 | 2 | 0 | 0,726562 | 0 | 0 | 0,48056 | 0 | 0 | 0 | 0,884019 | 0 | 0,3974 | 0 | SM_36:0;2 |
| SM_36:1;2 | SM | MEM | 36 | 1 | 2 | 6,23058 | 6,25922 | 6,19588 | 4,1869 | 5,38889 | 6,14798 | 3,19312 | 3,54839 | 4,05126 | 6,30399 | 3,17583 | 3,58402 | SM_36:1;2 |
| SM_38:1;2 | SM | MEM | 38 | 1 | 2 | 2,44934 | 2,19743 | 2,4116 | 2,04676 | 2,23825 | 2,12584 | 1,5161 | 1,47186 | 1,55715 | 2,52219 | 1,27372 | 1,66927 | SM_38:1;2 |
| SM_40:1;2 | SM | MEM | 40 | 1 | 2 | 10,6152 | 12,2483 | 12,5571 | 8,87143 | 9,91101 | 10,4443 | 7,25341 | 7,29631 | 11,5141 | 11,792 | 7,03377 | 6,94399 | SM_40:1;2 |
| SM_40:2;2 | SM | MEM | 40 | 2 | 2 | 2,11018 | 2,23243 | 2,61983 | 1,90337 | 1,77332 | 2,38696 | 1,62187 | 1,93955 | 1,99046 | 2,3789 | 1,55766 | 1,51912 | SM_40:2;2 |
| SM_42:1;2 | SM | MEM | 42 | 1 | 2 | 13,8986 | 16,8212 | 20,359 | 11,1889 | 13,0979 | 13,8684 | 12,1948 | 8,87773 | 20,5042 | 14,9297 | 12,5153 | 9,2838 | SM_42:1;2 |
| SM_42:2;2 | SM | MEM | 42 | 2 | 2 | 48,3928 | 63,47 | 66,8172 | 47,605 | 56,0235 | 49,002 | 33,1294 | 39,8842 | 56,9522 | 59,1713 | 42,3604 | 41,0959 | SM_42:2;2 |
| SM_42:2;3 | SM | MEM | 42 | 2 | 3 | 0 | 0,664268 | 0 | 0 | 0,829406 | 0 | 0 | 0,497067 | 1,01339 | 0 | 0,617753 | 0 | SM_42:2;3 |
| TAG_50:1 | TAG | STO | 50 | 1 | 0 | 1,21906 | 1,53716 | 2,43027 | 0 | 1,84069 | 2,18658 | 1,3096 | 0 | 0 | 1,59979 | 0 | 1,08655 | TAG_50:1 |
| TAG_50:2 | TAG | STO | 50 | 2 | 0 | 1,05779 | 1,36808 | 2,22165 | 1,24184 | 1,48469 | 1,90462 | 0 | 1,36899 | 0 | 1,49469 | 1,03546 | 1,09171 | TAG_50:2 |
| TAG_50:3 | TAG | STO | 50 | 3 | 0 | 0 | 0 | 1,11345 | 0 | 0 | 0 | 0 | 1,23604 | 0 | 0 | 0 | 0 | TAG_50:3 |
| TAG_52:2 | TAG | STO | 52 | 2 | 0 | 1,27858 | 2,02347 | 2,59565 | 1,4035 | 1,7247 | 2,52734 | 1,25282 | 1,19039 | 0 | 2,03797 | 1,14196 | 1,17871 | TAG_52:2 |
| TAG_52:3 | TAG | STO | 52 | 3 | 0 | 1,50793 | 1,66153 | 2,93628 | 1,79871 | 2,03811 | 2,47961 | 1,20405 | 1,17854 | 0 | 1,95097 | 1,44891 | 1,50701 | TAG_52:3 |
| TAG_52:4 | TAG | STO | 52 | 4 | 0 | 0 | 0 | 1,4655 | 0 | 1,04887 | 1,40533 | 0,922623 | 0,968101 | 0 | 0,994619 | 0 | 0 | TAG_52:4 |
| TAG_54:2 | TAG | STO | 54 | 2 | 0 | 1,03708 | 0 | 1,12917 | 1,00444 | 1,02568 | 1,2505 | 0 | 1,02736 | 0 | 0 | 1,4218 | 0 | TAG_54:2 |
| TAG_54:3 | TAG | STO | 54 | 3 | 0 | 0,86575 | 1,20562 | 1,21918 | 1,18852 | 1,29521 | 1,17862 | 0 | 0,78991 | 0 | 0,960864 | 1,00765 | 0 | TAG_54:3 |
| TAG_54:4 | TAG | STO | 54 | 4 | 0 | 1,36213 | 1,16768 | 2,15115 | 1,50588 | 1,42919 | 1,76981 | 0,765739 | 0,825521 | 0 | 1,62372 | 1,12148 | 1,38899 | TAG_54:4 |
| TAG_54:5 | TAG | STO | 54 | 5 | 0 | 0 | 0 | 1,35235 | 0,77887 | 0,860836 | 1,11221 | 0 | 0 | 0 | 1,01524 | 0 | 0 | TAG_54:5 |
